# Amplification is the Primary Mode of Gene-by-Sex Interaction in Complex Human Traits

**DOI:** 10.1101/2022.05.06.490973

**Authors:** Carrie Zhu, Matthew J. Ming, Jared M. Cole, Michael D. Edge, Mark Kirkpatrick, Arbel Harpak

## Abstract

Sex differences in complex traits are suspected to be in part due to widespread gene-by-sex interactions (GxSex), but empirical evidence has been elusive. Here, we infer the mixture of ways polygenic effects on physiological traits covary between males and females. We find that GxSex is pervasive but acts primarily through systematic sex differences in the magnitude of many genetic effects (“amplification”), rather than in the identity of causal variants. Amplification patterns account for sex differences in trait variance. In some cases, testosterone may mediate amplification. Finally, we develop a population-genetic test linking GxSex to contemporary natural selection and find evidence for sexually antagonistic selection on variants affecting testosterone levels. Taken together, our results suggest that the amplification of polygenic effects is a common mode of GxSex that may contribute to sex differences and fuel their evolution.

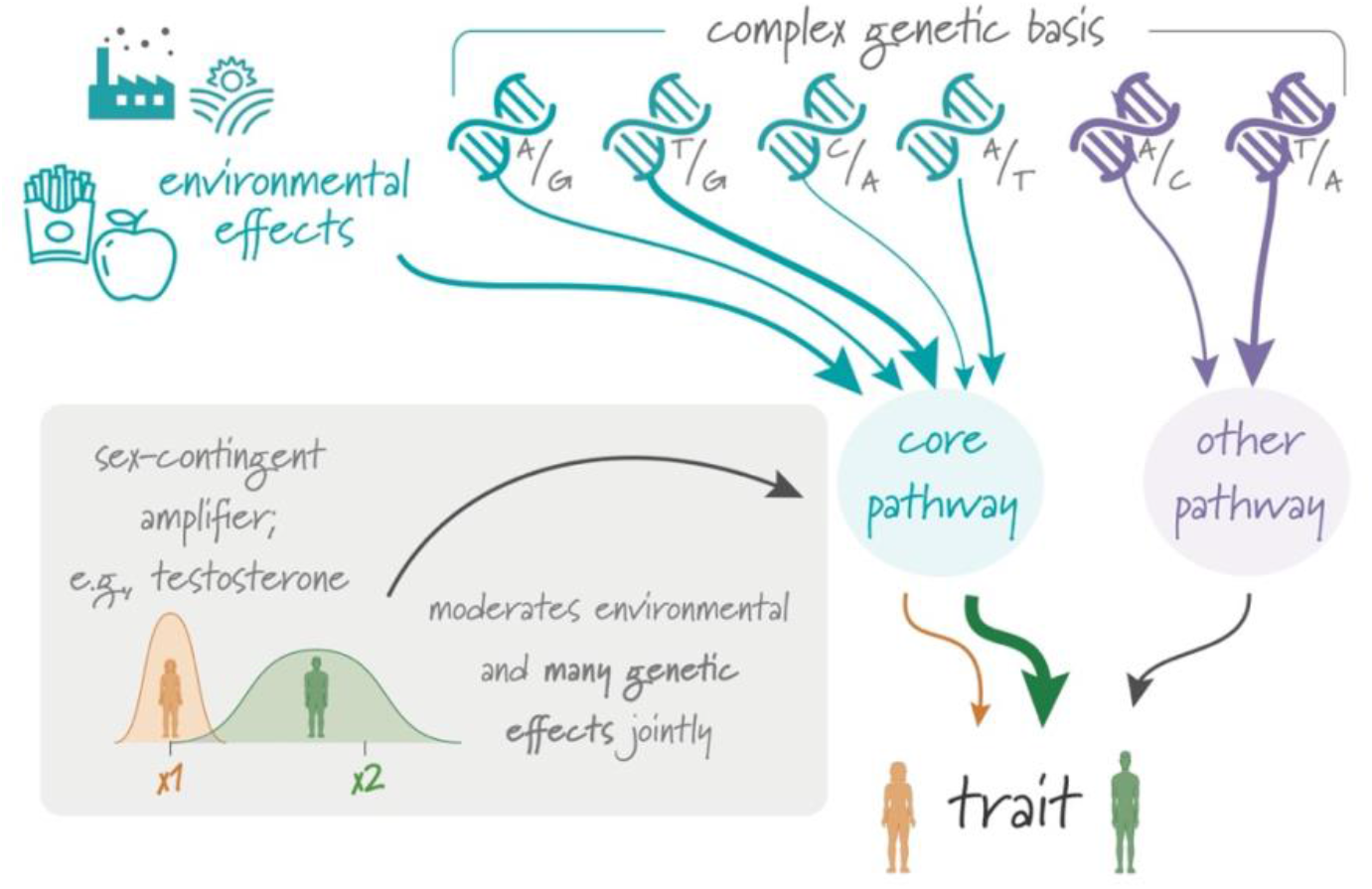

## Introduction

Genetic effects can depend on context. If the distribution of contexts differs between groups of people, as they do for males and females, so should the average genetic effects on traits^1,2^. In particular, such gene-by-sex interaction (GxSex) may be a result of sex differences in bodily, environmental and social contexts or epistatic interaction with sex chromosomes^3–9^. Sex differences in genetic effects on complex traits are clearly of high evolutionary^8,10–14^ and translational^9,15–22^ importance. Yet with the exception of testosterone levels^23–26^, the basis of sexual dimorphism in complex traits is not well understood^19^. To date, empirical evidence for GxSex in GWAS data—whether focused on identifying large GxSex effects at individual loci or by estimating genetic correlations between the sexes for polygenic traits—has been lacking.

Here, we set out to study governing principles of GxSex in complex human traits and explain why current approaches for characterizing GxSex may be lacking for this goal. We then suggest a mode of GxSex that may have gone largely underappreciated: A systematic difference in the magnitude of effect of many variants between the sexes, which we refer to as “amplification”^27^. Amplification can happen for a large set of variants regulating a specific pathway if the pathway responds to a shared cue^28–31^. In classic hypothesis-testing approaches that test for a GxSex effect separately in each variant, the signal of amplification may be crushed under the multiple hypothesis burden. On the other hand, even state-of-the-art tools designed with complex traits in mind may miss amplification signals: They often treat genetic correlation (between GWAS estimates based on samples from two environments) as a litmus test for whether effects are the same in two groups^32–36^, but correlations are scaleless and thus may entirely miss amplification effects.

We developed a new approach for flexibly characterizing a mixture of male-female genetic covariance relationships and applied it to 27 physiological traits in the UK Biobank. We found that amplification is pervasive across traits, and that considering amplification helps explain sex differences in phenotypic variance. Finally, we consider an implication of polygenic GxSex for sexually antagonistic selection: Our model confirms that variants that affect traits may be subject to sexually antagonistic selection when male and female trait optima are very different or, surprisingly, even if the trait optima are very similar. We developed a novel test for sexually antagonistic polygenic selection, which connects GxSex to signals of contemporary viability selection. Using this test, we find subtle evidence of sexually antagonistic selection on variants affecting testosterone levels.

## Results

### The limited scope of single-locus analysis

We conducted GWASs stratified by sex chromosome karyotype for 27 continuous physiological traits in the UKB using a sample of ~150K individuals with two X chromosomes and another sample of ~150K individuals with XY, and a combined sample that included both the XX and XY samples. We chose to analyze traits with SNP heritabilities over 7.5% in the combined sample, to have higher statistical power. While there is not a strict one-to-one relationship between sex chromosome karyotype and biological sex, we label XX individuals as females and XY individuals as males, and view these labels as capturing group differences in distributions of contexts for autosomal effects, rather than as a dichotomy^9,22,37^. Throughout, we analyze GWAS on the raw measurement units as provided by UKB. (See note on the rationale behind this choice in the section **Amplification of genetic effects is the primary mode of GxSex**).

Among the 27 traits, we observed substantial discordance between males and females in associations with the trait only for testosterone and waist:hip ratio (whether or not it is adjusted for BMI; **Fig. S1**). For testosterone, as noted in previous analyses, associated genes are often in separate pathways in males and females^23,25^. This is reflected in the small overlap of genes neighboring top associations in our GWAS. For example, in females, the gene CYP3A7 is involved in the hydroxylation of testosterone, resulting in its inactivation. In males, FKBP4 plays a role in the downstream signaling of testosterone on the hypothalamus. Both genes, to our knowledge, do not affect testosterone levels in the other sex.

For waist:hip ratio, we saw multiple associations in females only, such as variants near ADAMTS9, a gene involved in insulin sensitivity^38^. As previous work established^23,25,26^, testosterone and waist:hip ratio are the exception, not the rule: Most traits did not display many sex differences in top associations. For instance, arm fat-free mass, a highly heritable dimorphic trait, showed near-perfect concordance in significant loci (**Fig. S1**). A previous study^26^ examining the concordance in top associations between males and females found few uniquely-associated SNPs (<20) across the 84 continuous traits they studied; waist:hip ratio was an exception with 100 associations unique to one sex. Considering the evidence for the polygenicity of additive genetic variation affecting many complex traits^39–41^, it stands to reason that looking beyond lead associations, through a polygenic prism, may aid in the characterization of non-additive effects (such as GxSex) as well.

### The limited scope of analyzing GxSex via heritability differences and genetic correlations

We therefore turned to consider the polygenic nature of GxSex, first by employing commonly-used approaches: comparing sex-specific SNP heritabilities and examining genetic correlations. We used LD Score Regression (LDSC)^36,42^ to estimate these for each trait. In most traits (17/27), males and females had a genetic correlation greater than 0.9. Testosterone had the lowest genetic correlation of 0.01, which suggests very little sharing of signals between males and females (see similar results by Flynn et al.^25^ and Sinnott-Armstrong et al.^23^).

For the majority of traits (18/27), male and female heritabilities were both greater than the heritability in a sample that included both sexes. For instance, in arm fat-free mass (right), the heritability in the both-sex sample was 0.232 (± 0.009), while the heritabilities for male and female were 0.279 (± 0.012) and 0.255 (± 0.011), respectively. In particular, all body mass-related traits, excluding BMI-adjusted waist:hip ratio, had greater sex-specific heritabilities **(Fig. 1)**.

**Figure 1:**
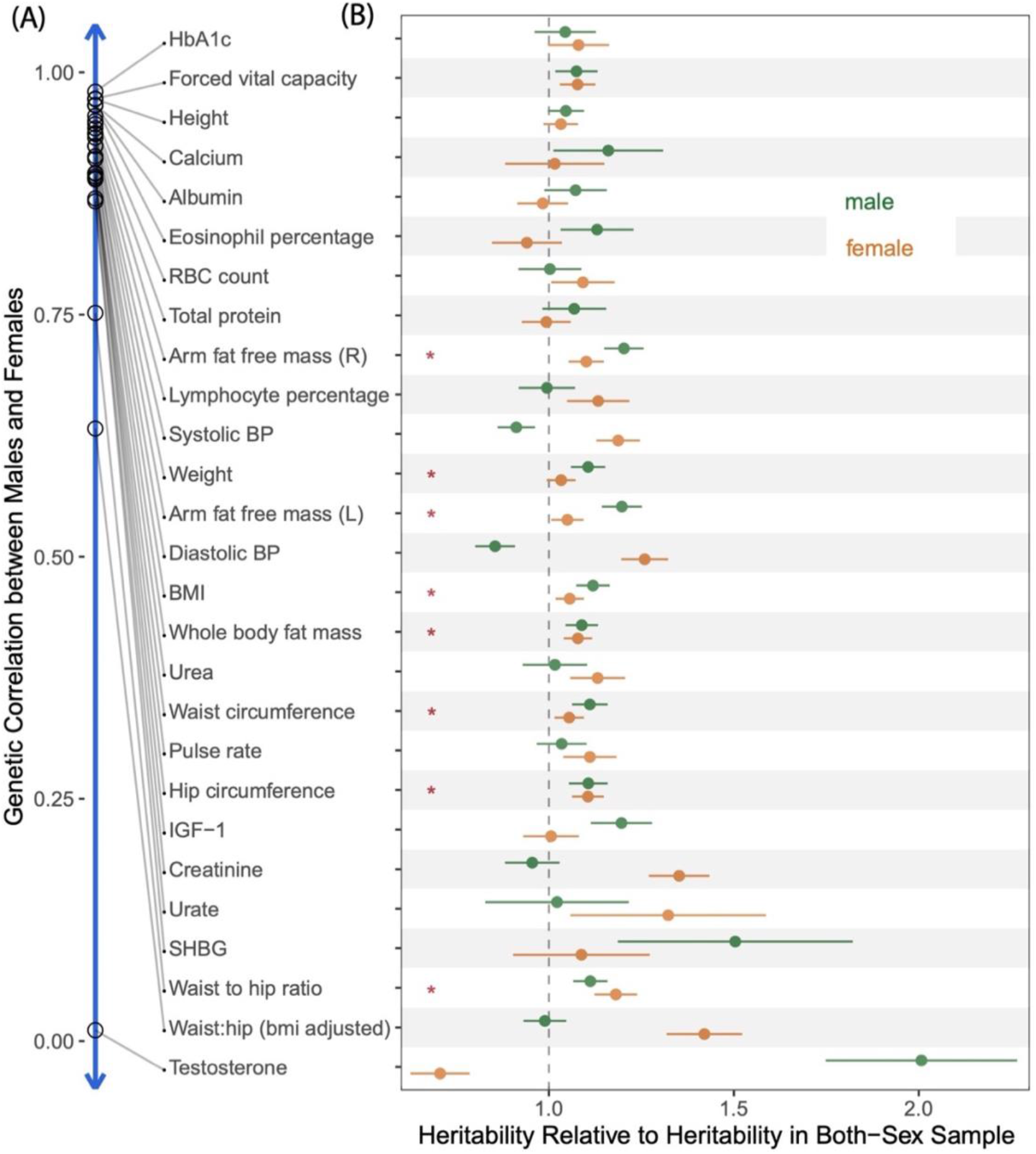
Heritabilities and Genetic Correlations Cannot Fully Distinguish Models of GxSex. **(A)** Genetic correlations between the male and females, estimated using bi-variate LD Score Regression, are shown in descending order. **(B)** The x-axis represents the relative heritability, i.e., the SNP heritability divided by the SNP heritability estimated in the sample with both sexes combined. Red asterisks show body-mass related traits with greater heritabilities in both sex-specific samples than in the sample combining both sexes.

In addition, we noticed a trend in which, as the genetic correlation decreased, the difference between the heritabilities within each sex and in the sample combining both sexes tended to become larger (Pearson r = −0.88, paired t-test p-value = 10^−10^, **Fig. 1**). Nonetheless, several traits with genetic correlation above 0.9 also present relatively large sex differences in heritability: For example, diastolic blood pressure and arm fat-free mass (left) had differences of 5.2% (two-sample t-test p-value = 3 · 10^−6^) and 3.4% (two-sample t-test p-value = 0.04), respectively. These examples are incompatible with a model of pervasive uncorrelated genetic effects driving sex-specific genetic contributions to variation in the trait (**Table 1**, second model).

**Table 1.** Polygenic Models of GxSex. We examine different models of the nature of GxSex in complex traits that link to previous studies and motivations. Each model leads to different expectations from the analysis of heritability and genetic correlations (**Fig. 1**). The illustrations in the third column depict examples of directions and magnitudes of genetic effects, corresponding to each model. 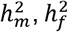 and *h*^2^ denote narrow-sense heritabilities in males, females, and a combined sample, respectively.

| Model | Motivation | Illustration of Effect Covariance | Expectation from Heritability Analysis |
| --- | --- | --- | --- |
| No GxSex | Little previous evidence for GxSex | | (a) $h_m^2$ can only differ from $h_f^2$ through environmental variance differences<br>(b) $h_m^2 < h^2$ or $h_f^2 < h^2$ |
| Weakly or negatively correlated genetic effects | Sexual dimorphism is pervasive and heritable contribution is expected to lie primarily in autosomes | | (a) Low or negative genetic correlation<br>(b) $h_m^2, h_f^2 > h^2$, and the larger the difference, the lower the genetic correlation |
| Highly correlated effects, difference in magnitude ("amplification") | Response to cues such as testosterone; evidence for GxE in non-human organisms | | (a) High genetic correlation<br>(b) $h_m^2 < h^2$ or $h_f^2 < h^2$ |
| Mixture of covariance relationships | Heritability analysis often incompatible with either model or cannot distinguish between models |  | Compatible with all observations; motivates:<br>(a) Direct estimation of genetic effect covariance, rather than sole reliance on heritability estimates<br>(b) Modeling mixture components |

We therefore considered two other alternative hypotheses under a simple additive model of variance in a trait. Differences in heritability are either due to sex differences in genetic variance, in environmental variance, or both. If genetic effects are similar, differences in environmental variance alone could cause heritability differences (**Table 1**, first model). But as we show in the **Methods** section, under such a model, the heritability in the combined sample cannot be smaller than both sex-specific heritabilities.

Therefore, the observation of higher sex-specific heritabilities for most traits suggests that the genetic variance must differ between males and females. Given the random segregation of autosomal alleles, independent of an individual’s sex chromosome karyotype, and assuming, further, that there is little-to-no interaction of sex and genotype affecting participation in the UKB^43^, UKB allele frequencies in males and females are expected to be very similar. Thus, this observation suggests that causal genetic effects differ between males and females for most traits analyzed.

A last hypothesis that might tie together the observations in **Table 1** is a less appreciated mode of GxSex, amplification. Namely, that the identity and direction of effects are largely shared between sexes (leading to high genetic correlation), but the magnitude of genetic effects differs—e.g., larger genetic effects on blood pressure in females—which in turn lead to differences in genetic variance (**Table 1**, third model).

We can test the hypothesis that amplification acts systematically—across a large fraction of causal variants—by examining the effects of polygenic scores (PGSs), genetic predictors of a complex trait. Under this hypothesis, regardless of whether the PGS is estimated in a sample of males, females, or a combined sample of both males and females, it should be predictive in both sexes, since the causal variants and the direction of their effects are shared and the magnitude is correlated (**Table 1**, third model). At the same time, in the sex for which genetic effects are larger, the effect of the PGS is expected to be larger. To evaluate evidence for the systematic amplification model, we estimated PGSs based on our sex-specific GWASs, and examined their effect in both sexes. For some traits, like albumin and lymphocyte percentage, the effects of the same PGS on trait value in males and females were statistically indistinguishable (**Fig. 2A,E,I,J**). In a few other traits, such as diastolic blood pressure, the result was contingent on the sample in which the PGS was estimated (**Fig. 2C,G,I,J**). However, for roughly half of the traits examined, regardless of the sample in which the PGS was derived, the effect of the PGS was predictive in both sexes yet significantly larger in one of the sexes (17/27 traits with t-test p-value < 0.05 using the PGS derived from the males sample; 13/27 using the PGS derived from the females sample; **Fig. 2B,D,F,H,I,J**). These observations are consistent with systematic amplification.

**Figure 2:**
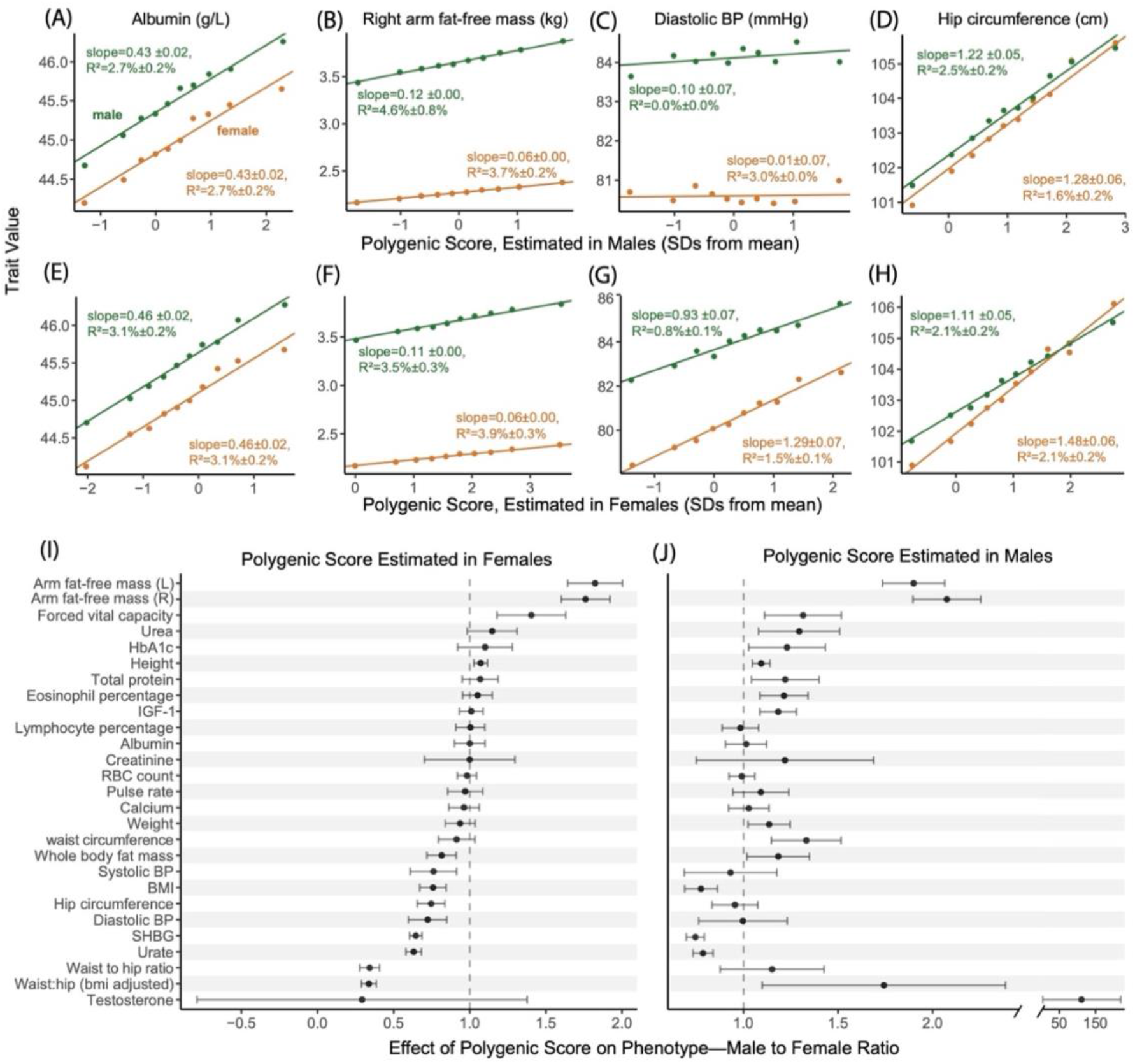
Evaluating evidence for systematic amplification. **(A-D)** We regressed trait values in males (green) and separately in females (orange) on a polygenic score estimated in an independent sample of males. Points show mean values in one decile of the polygenic score; the fitted line and associated effect estimate and *R*^2^ correspond to regressions on the raw, non-binned data. In some traits, like Albumin (A), the polygenic score has a similar effect on the trait in both sexes. In other traits (B,D), the estimated effect of the polygenic score differs significantly, consistent with a substantial difference in the magnitude of genetic effects of sites included in the polygenic score. **(E-H)** Same analysis as A-D, but with a polygenic score pre-estimated in an independent sample of females. **(I-J)** Summary of the ratio of the effect of the polygenic score on the trait (±2 SE) in males to the effect in females across physiological traits. See results for other traits in **Fig. S12**.

The results presented in **Figs. 1,2** suggested to us that various modes of polygenic GxSex ought to be jointly evaluated. None of the hypothesized rules of thumb (**Table 1**) for interpreting genetic correlations and sex differences in heritability worked across all traits (see also relevant discussion in Khramtsova et al.^9^). This motivated us to estimate the covariance between genetic effects in males and females directly. Another reason to treat covariance of genetic effects themselves as the estimand of interest is that multiple, distinct GxSex patterns may exist across subsets of genetic factors affecting a trait (**Table 1**, fourth model).

### Flexible model of sex-specific genetic effects as arising from a mixture of covariance relationships

We set to infer the mixture of covariance relationships of genetic effects among the sexes directly. We analyzed all traits in their raw measurement units as provided by the UKB. In particular, we did not normalize or standardize phenotypes within each sex before performing the sex-stratified GWAS, because sex differences in trait variance may be partly due to amplification. Standardization would have therefore resulted in masking amplification signals that may exist in the data. In some cases, this is indeed the purpose of standardization^44^. More generally, while each scaling choice has it merits, we view the measurement of genetic effects in their raw units as the most biologically interpretable.

We used multivariate adaptive shrinkage (*mash*)^45^, a tool that allows the inference of genome-wide frequencies of genetic covariance relationships. Namely, we model the marginal SNP effect estimates as sampled (with SNP-specific, sex-specific noise) from a mixture of zero-centered Normal distributions with various prespecified covariance relationships (2×2 Variance-Covariance matrices for male and female effects; Eq. 1 in Urbut et al.^45^). Our prespecified covariance matrices (“hypothesis matrices”) span a wide array of amplification and correlation relationships, and use *mash* to estimate the mixture weights. Loosely, these weights can be interpreted as the proportion of variants that follow the pattern specified by the covariance matrix (**Fig. 3A**). Our covariance matrices ranged from −1 to 1 in between-sex correlation, and 10 levels of relative magnitude in females relative to males, including matrices corresponding to no effect in one or both sexes (**Fig. S2**).

**Figure 3:**
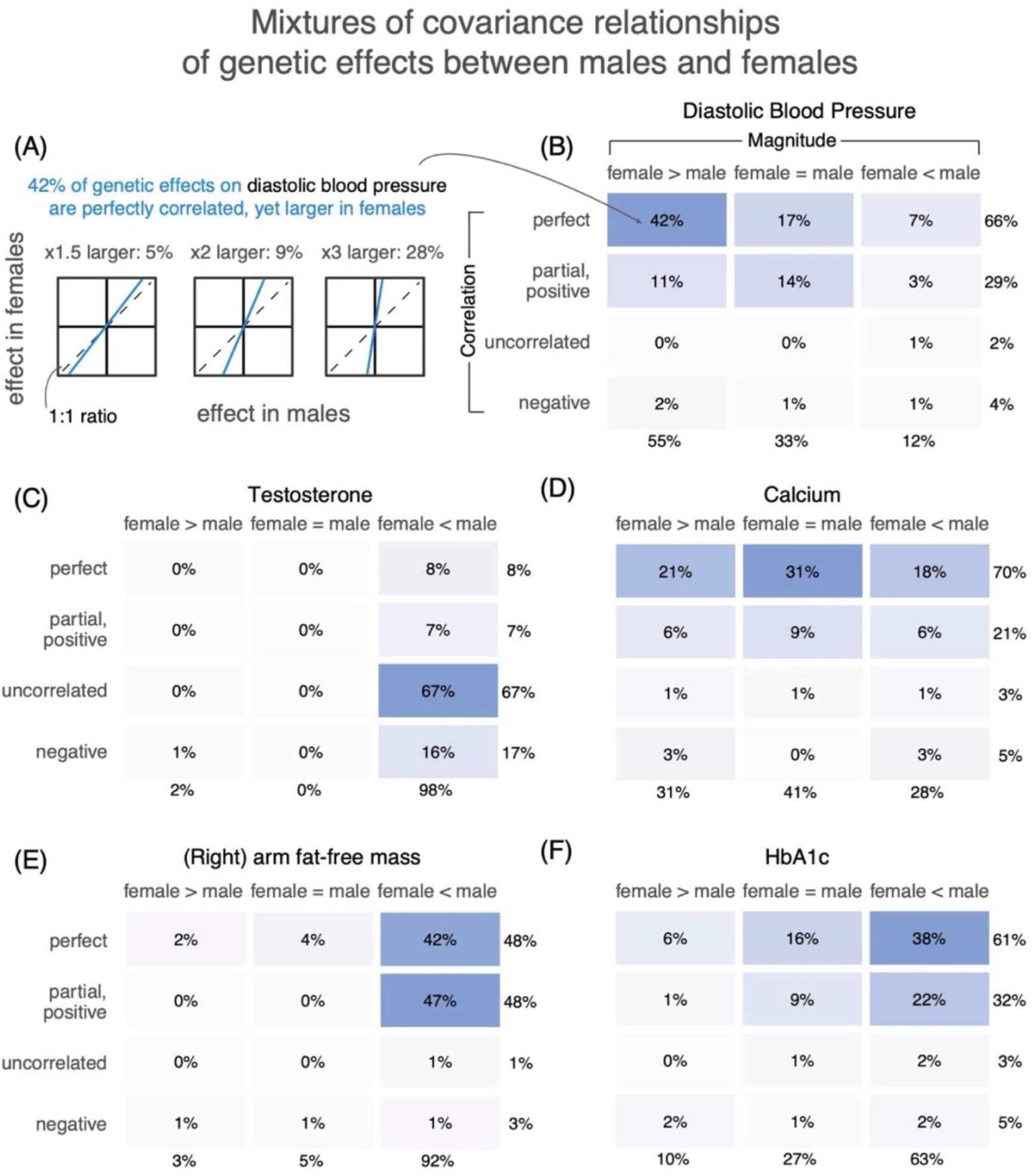
Polygenic covariance structure between males and females. **(A)** Our analysis of the polygenic covariance between males and females is based on sex-stratified GWAS. We modelled the sex-stratified GWAS estimates as sampled with error from true effects arising from a mixture of possible covariance relationships between female and male genetic effects. As an example, shown are illustrations for three possible relationships of the same qualitative nature—perfectly correlated effects which are also larger in females—and the mixture weights estimated for each in the case of diastolic blood pressure. **(B-F)** Each box shows the sum of weights placed on all covariance relationships of the same qualitative nature, as specified by relative magnitude (horizontal axis) and correlation (vertical axis) between male and female effects. The full set of pre-specified covariance matrices is shown in **Fig. S2**, and the weights placed on each of them for each trait are shown in **Fig. S5**. All weights shown are percentages of non-null weights, i.e., the weight divided by the sum of all weights except for the one corresponding to no effect in either sex.

We first focus on testosterone, for which previous research sets the expectation for polygenic male-female covariance. In terms of magnitude, the vast majority of effects should have much greater effect in males. In terms of correlation, we expect a class of genetic effects acting through largely independent and uncorrelated pathways alongside a class of effects via shared pathways^23^. Independent pathways include the role of hypothalamic-pituitary-gonadal axis in male testosterone regulation and the contrasting role of the adrenal gland in female testosterone production. Shared pathways involve sex hormone-binding globulin (SHBG), which decreases the amount of bioavailable testosterone in both males and females. As expected, we found that mixture weights for testosterone concentrated on greater magnitudes in males and largely uncorrelated effects. Out of the 32% total weights on matrices with an effect in at least one sex, 98% of the weights were placed on matrices representing larger effects in males, including 20.4% (± 0.7%) having male-specific effects (**Fig. 3, S5**).

### Amplification of genetic effects is the primary mode of GxSex

The only trait of the 27 where a large fraction (≥10%) of non-zero effects were negatively correlated was testosterone (17%). Most effects were instead perfectly or near-perfectly correlated. For example, diastolic blood pressure and eosinophil percentage had 66% (**Fig. 3**) and 68% (**Fig. S5**) of effects being perfectly correlated, respectively. Overall, the low weights on matrices representing negative correlation do not support opposite directions of effects being a major mode of GxSex (**Fig. S8**).

In some traits, such as hemoglobin A1C or diastolic blood pressure, previously considered non-sex-specific because of high genetic correlations between sexes and a concordance in top GWAS hits, we find evidence for substantial GxSex through amplification (**Fig. 3B,F; Fig. S5**)^25,26^. Furthermore, about half (13/27) of the traits analyzed had the majority of weights placed on greater effects in just one of the sexes (x-axis in **Fig. 4A**). For instance, 92% of effects on BMI-adjusted waist:hip ratio were greater in females and 92% of effects on (right) arm fat-free mass were greater in males. Both traits had mixture weights concentrated on highly correlated effects (**Fig. 3**). We confirmed, using a simulation study, that this summary of sex-biased amplification indeed captures sex differences in the magnitude of genetic effects and that it is not due to differences in the extent of estimation noise (e.g., variation in environmental factors independent of genetic effects; **Figs. S6-7**; **Methods**).

**Figure 4.**
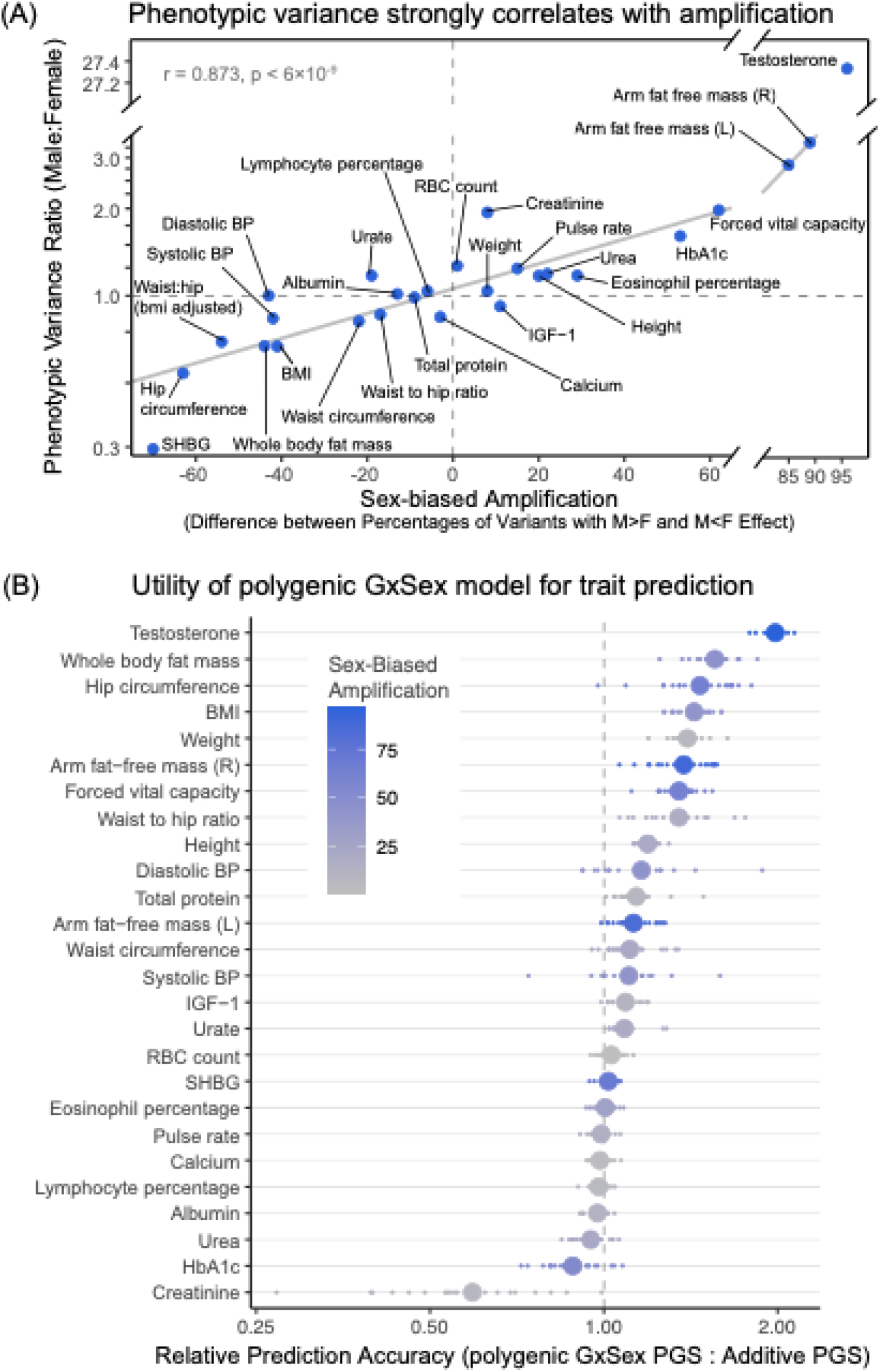
**(A) Phenotypic variance strongly correlates with amplification.** The x-axis summarized the “sex-biased amplification” of polygenic effects and is calculated by taking the difference between the sum of weights on matrices with male effects greater in magnitude than female effects (M>F) and the sum of weights of M<F matrices. The solid gray line shows a linear fit across traits, excluding testosterone as an outlier. **(B) Utility of polygenic GxSex model for trait prediction.** The x-axis shows the relative prediction accuracy estimated from the incremental R^2^ ratio of a GxSex model informed by polygenic covariance patterns and an additive model. For each trait, smaller points show relative prediction accuracy across 20 cross-validation folds, and larger points show the average across the 20 folds. The phenotypes are ordered by the relative prediction accuracy. The color of each point corresponds to the degree of sex-biased amplification as described in (A).

Across traits, the difference between the fraction of male-larger effects and the fraction of female-larger effects correlates strongly with male-to-female phenotypic variance ratio (Pearson r = 0.873, p-value = 6·10^−9^ after removing testosterone as an outlier; **Fig. 4A**). This observation is consistent with our hypothesis of amplification leading to differences in genetic variance between sexes and thereby contributing substantially to sex differences in phenotypic variance. Together, these observations point to amplification, rather than uncorrelated effects, as a primary mode of polygenic GxSex.

Another important question about the implication of pervasive amplification is whether it is a major driver of mean phenotypic differences. The ratio between male and female phenotypic means is correlated with the difference between male-larger and female-larger amplification (Pearson r = 0.75; p-value = 2 · 10^−5^ after removing testosterone and BMI-adjusted waist:hip ratio as outliers). Though this correlation is intriguing, within-sex GWAS aims to explain individual differences from the mean of the sex, and such GWAS results do not dictate the values of the sex means. Further, both the ratio of mean trait values between sexes and the difference in amplification are strongly correlated with phenotypic variance ratios (**Fig. 4A**; **Fig. S9**; see also Karp et al.^8^),, and many different causal accounts could explain these correlations.

Finally, the pervasiveness of GxSex, alongside the mixture of covariance relationships across the genome for many traits, may be important to consider in phenotypic prediction. We compared the prediction accuracy of PGSs that consider the polygenic covariance structure to that of additive models, that ignore GxSex, as well as models that include GxSex but do not consider the polygenic covariance structure (**Supplementary Materials; Fig. S13**). Indeed, for most traits (20/27 traits; **Fig. 4B**), models that consider the polygenic covariance structure outperform all other models evaluated. Traits which showed better prediction accuracy using the model that considered polygenic covariance structure included many body mass-related traits such as BMI and whole body fat mass that also tended to have higher sex-based amplification (**Fig. 4B;** Pearson *r* = 0.54, *p* = 0.004 between sex-biased amplification and prediction accuracy ratio). These results point to the utility of considering polygenic covariance structure in polygenic score prediction.

### Testosterone as an amplifier

Thus far, we treated the genetic interaction as discretely mediated by biological sex. One mechanism that may underlie GxSex is a cue or exposure that modulates the magnitude (and less often, the direction) of genetic effects, and varies in its distribution between the sexes. A plausible candidate is testosterone. Testosterone may be a key instigator since the hormone is present in distinctive pathways and levels between the sexes and a known contributor to the development of male secondary characteristics, so therefore could modulate genetic causes on sex-differentiated traits.

To test this idea, we first binned individuals of each sex by their testosterone levels. Then, for each trait and within the bin, we quantified the magnitude of total genetic effect as the linear regression coefficient of trait to a PGS for the trait (**Methods**; see **Fig. S15** for results obtained using sex-specific PGS). For BMI, testosterone (mean per bin) and the magnitude of genetic effect were correlated for both males and females (Pearson p-value < 0.05; **Fig. 5A**). For all body-mass-related traits, there was a negative correlation between the magnitude of genetic effect and testosterone levels for males and a positive correlation for females (**Fig. 5B**). Since the relationship with testosterone remains contingent on sex, a model of testosterone as the sole driver of the observed sex-specificity would be invalid. These observations may help explain previous reports of positive correlations between obesity and free testosterone in women, and negative correlations in men^46^. We conclude that in body-mass related traits, testosterone may be modulating genetic effects in a sexually antagonistic manner.

**Figure 5.**
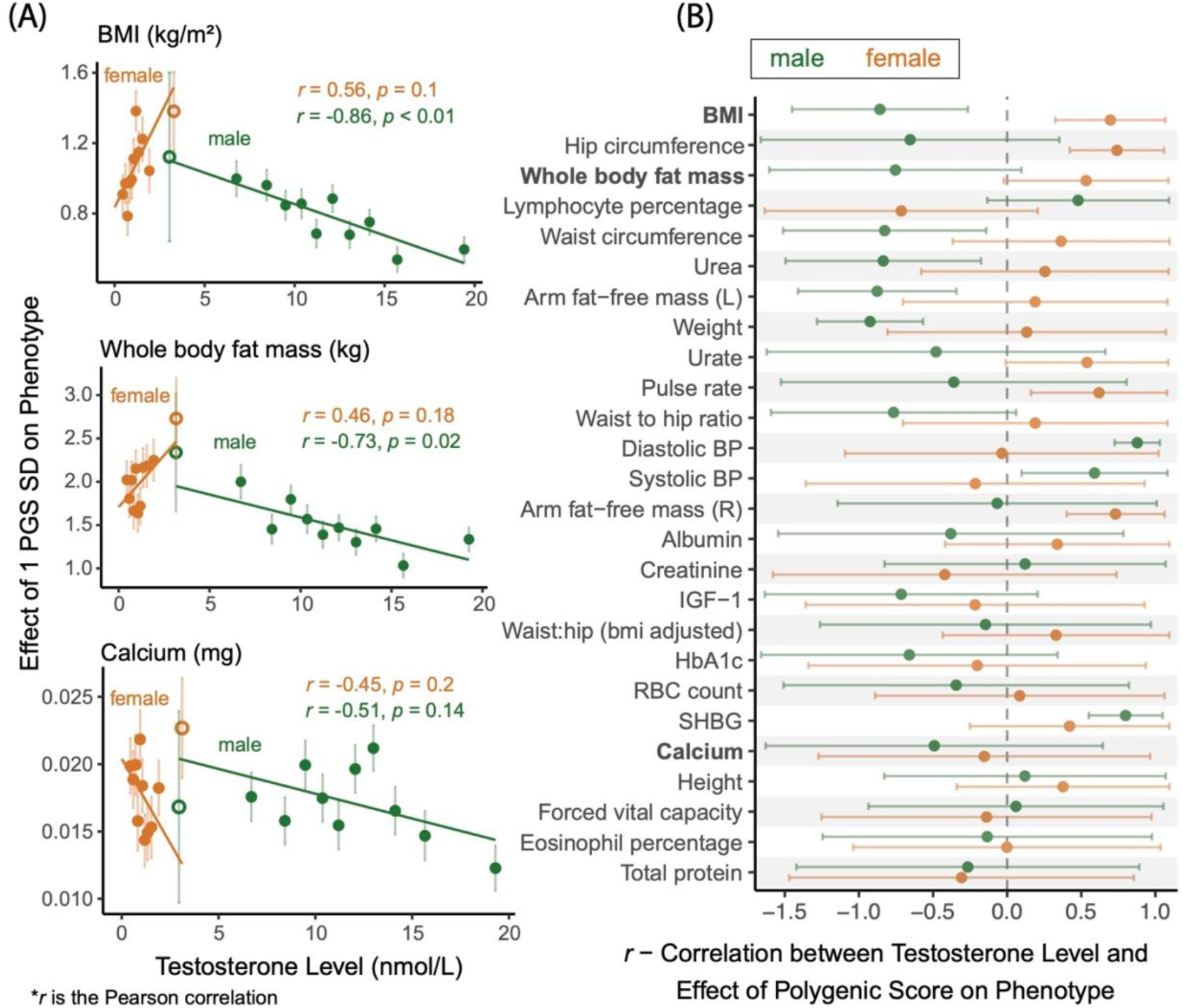
Amplification of total genetic effect in relation to testosterone levels. **(A)** The relationship between testosterone level bins and estimated magnitude of genetic effect on traits is shown for three traits. The magnitude of genetic effect is estimated using the slope of the regression of phenotypic values to polygenic scores in that bin. The units on the y-axis are effect per standard deviations (SD) of the polygenic scores across all individuals in all bins. The hollow data points are bins with overlapping testosterone ranges between males and females; these are based on fewer individuals (~800 compared to ~2200 in other bins) and not included in the regression. **Fig. S14** show all other traits analyzed. **(B)** The correlation for each sex (90% CI) are shown for all 27 traits. Traits are ordered in descending order of male-female differences in Pearson correlation.

We performed two additional analyses designed to control for possible caveats to the association of testosterone and the magnitude of polygenic effect: An association test that controls for possible confounding with age (**Fig. S17**) and a test that mitigates confounding with other variables or reverse causality (wherein the magnitude of genetic effect affecting the focal trait causally affecting testosterone levels; **Fig. S16**). The evidence for an effect of testosterone on the magnitude of polygenic effect did not remain significant in either of these tests. It is possible, however, that this was due to low statistical power (**Methods**).

### Are polygenic and environmental effects jointly amplified?

Our results thus far suggest that polygenic amplification across sexes is pervasive across traits; and that the ratio of phenotypic variance scales with amplification (**Fig. 4A**). An immediate question of interest is whether the same modulators that act on the magnitude of genetic effects act on environmental effects as well (see also relevant discussion by Domingue et al.^47^). Consider the example of human skeletal muscle. The impact of resistance exercise varies between males and females. Resistance exercise can be considered as an environmental effect since it upregulates multiple skeletal muscle genes present in both males and females such as IGF-1, which in turn is involved in muscle growth^48^. However, after resistance exercise at similar intensities, upregulation of such genes is sustained in males, while levels return sooner to the resting state in females (**Fig. S18**). It is plausible that modulators of the effect of IGF-1, such as insulin^49^ or sex hormones^50,51^, drive a difference in the magnitude of effect of core genes such as IGF-1 in a sex-specific manner. To express this intuition with a model: If amplification mechanisms are shared, then amplification may be modeled as having the same scalar multiplier effect on genetic and environmental effects (**Fig. 6A**). In the **Methods** section, we specify the details of a null model of joint amplification, which yields the prediction that the male-female ratio of genetic variances should equal the respective ratio of environmental variances (blue line in **Fig 6B**).

**Figure 6.**
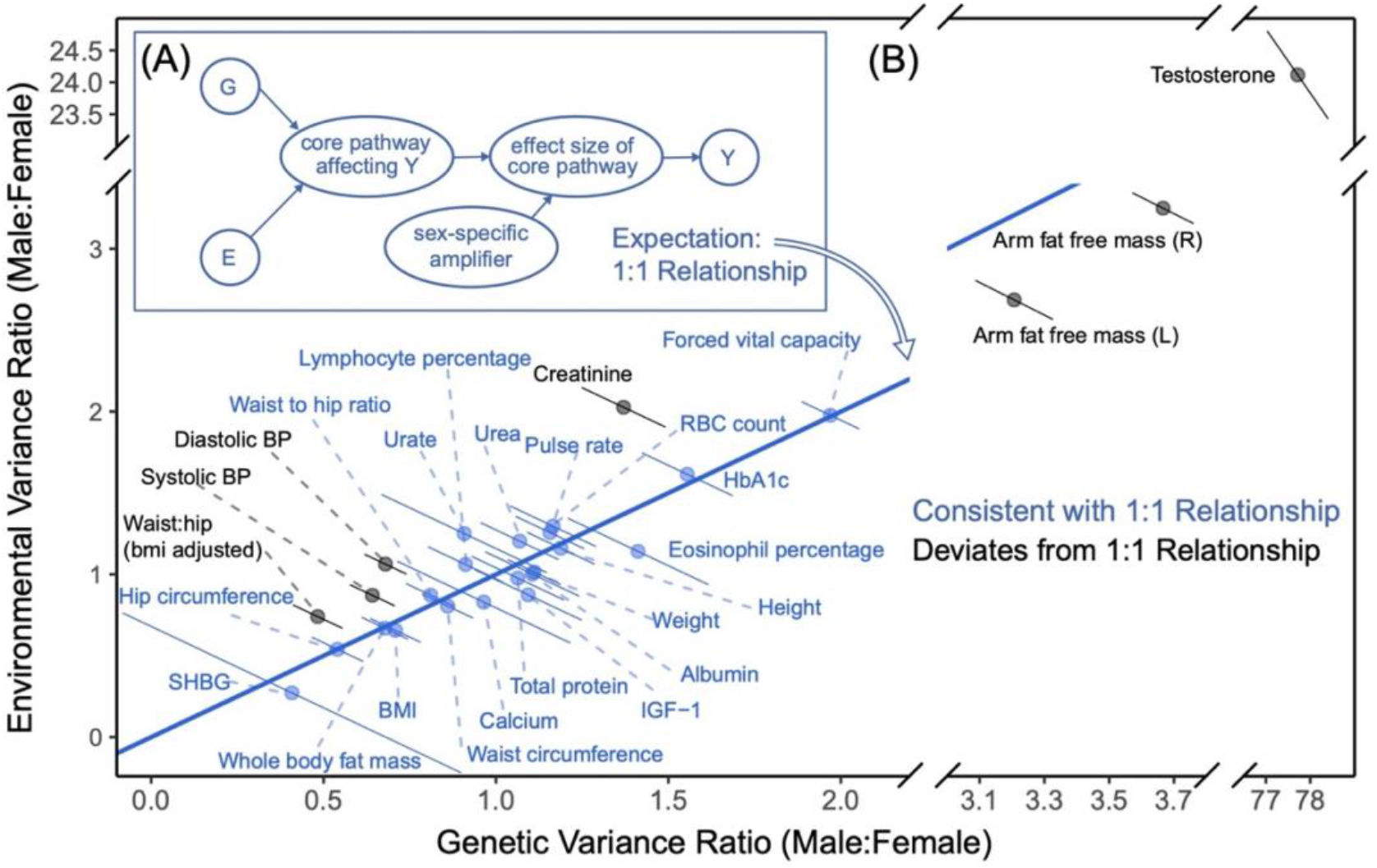
Testing a model of pervasive, joint amplification of environmental and polygenic effects. **(A)** A model of equal amplification of genetic (G) and environmental (E) effect, that produces the sex differences in the distribution of the phenotype, Y. G and E both act through a core pathway that is amplified in a sex-specific manner. **(B)** The blue 1:1 line depicts the theoretical expectation under a simple model of equal amplification of genetic and environmental effects in males compared to females. Error bars show 90% confidence intervals. Traits in blue are consistent (within their 90% CI) with the theoretical prediction. **Fig. S19** shows the same data alongside the predictions under other theoretical models of male-female variance ratios.

This expectation is qualitatively different from those of two longstanding theoretical “rules of thumb” predictions for sex differences in trait variance (**Supplementary materials**; **Fig. S19A;** of. Zajitschek et al.^52^): The “greater male variability” and “estrus-mediated variability” models, which provide a poor fit across the 27 physiological traits analyzed (**Fig. S19B**).

We tested the fit of the theoretical prediction under pervasive joint amplification across traits. We used our estimates of sex-specific phenotypic variance and SNP heritabilities to estimate the ratios of genetic and environmental variances. We note that environmental variance is proxied here by all trait variance not due to additive genetic effects, and caution is advised with interpretation of this proxy. Twenty of the 27 traits were consistent with the null model of pervasive joint amplification (within 90% CI; **Fig. 6B**). This finding may suggest a sharing of pathways between polygenic and environmental effects for these traits (**Fig. 6A**). Interesting exceptions include diastolic blood pressure—which was the strongest outlier (p-value = 3.06 · 10^−12^, single-sample z-test), excluding testosterone.

### Sexually antagonistic selection

A hypothesized cause of sexual dimorphism is sexually antagonistic selection, in which some alleles are beneficial in one sex yet deleterious in the other^11,12,14,53,54^. Sexually antagonistic selection is difficult to study using traditional population genetics methods because Mendelian inheritance equalizes autosomal allele frequencies between the sexes at conception, thereby erasing informative signals. One way around this limitation is to examine allele-frequency differences between the sexes in the current generation, known as “selection in real time”^14,55,56^. In this section, we consider a model of sexually antagonistic selection acting on a polygenic trait and use it to estimate the strength of contemporary viability selection acting on the 27 traits we analyzed.

Most theoretical models of sexually antagonistic selection on a trait under stabilizing selection usually posit either highly distinct male and female fitness optima or genetic variants affecting traits antagonistically. Our findings on pervasive amplification suggest that variant effects on traits tend to have concordant signs. Yet, under pervasive amplification, a somewhat surprising intuition arises: Alleles affecting a trait may frequently experience sexually antagonistic selection—both in the case in which trait optima for males and females are very distinct (**Fig. 7B**) and for the case in which they are similar (**Fig. 7A**).

**Figure 7.**
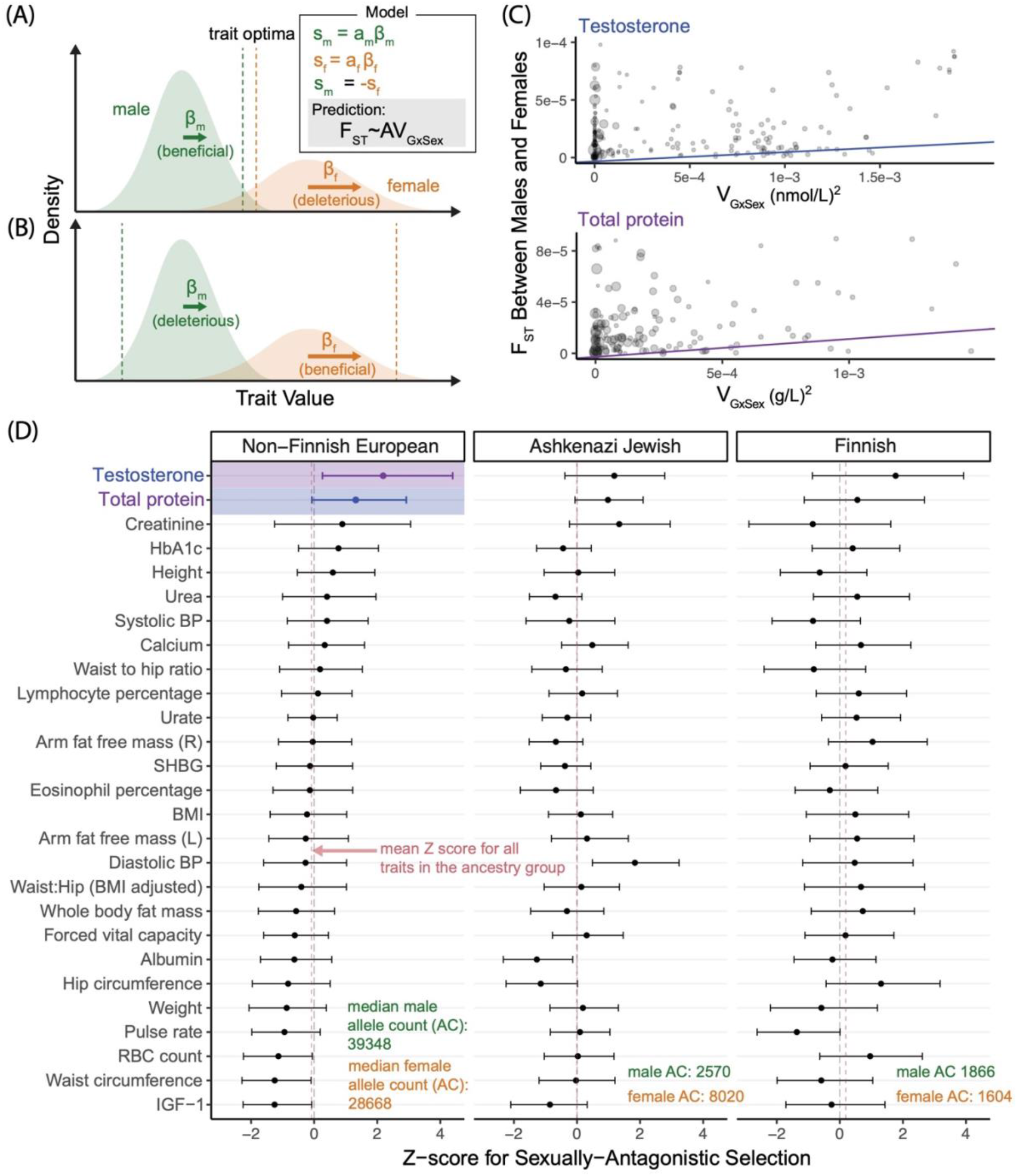
Testing for sexually antagonistic selection. **(A,B)** A model of sexually antagonistic selection. Selection coefficients, *s_m_* and *s_f_*, are linear with the additive effect on the trait in each sex. Sexually antagonistic selection acts such that *s_m_* = −*s_f_*. The model yields the prediction of **Eq. 1**. In (A), the effect of an allele tends to drive trait values towards the optimum in males, and away from the optimum in females. In (B), the fitness optima are farther in males and females; in both examples, selection on acts antagonistically (i.e., in opposite directions). **(C)** Two examples of the weighted least-squares linear regression preformed to estimate the strength of sexually antagonistic selection on variants associated with a trait (A in panel A and **Eq. 1**). Each point shows one SNP. Size is proportional to each point’s regression weight. **(D)** Z-scores (90% non-parametric bootstrap CI) estimated through 1000 resampling iterations of the weighted linear regression of panel B for each trait. The two colored estimates correspond to the examples in (B).

We developed a theoretical model of sexually antagonistic viability selection on a single trait that builds on this intuition. The model relates sex-specific effects on a complex trait to the divergence in allele frequency between males and females (measured as *F_ST_*^57,58^) due to viability selection “in real time”, i.e., acting in the current generation between conception and the time of sampling. We derive the expected relationship for each site *i*,

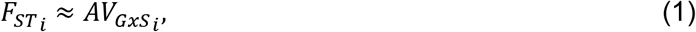

where

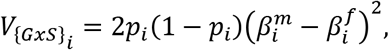

and *p_i_*, 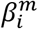 and 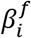 are the allele frequency of an allele at site *i*, its effect on the trait in males and its effect in females, respectively. *A* is a constant parameter shared across all variants and can therefore be interpreted as the effect of sexually antagonistic selection on male-female divergence at variants associated with the trait (**Methods**). We estimated *F_ST_i__* for all sites *i* across subsamples of various ancestry groups in the gnomAD dataset ^59^. To estimate *V*_{*GxS*}_*i*__ at each site and for each trait, we used our sex-stratified GWAS results. Since there is large heterogeneity in uncertainty of GxSex-genetic variance estimates, we use a variance-weighted linear regression to estimate *A* (see **Methods** for the derivation of the variance of *V*_{*GxS*}_*i*__ estimates and **Supplementary Materials** for further details).

Recent work has shown that apparent sex differences in autosomal allele frequencies within a sample are often due to a bioinformatic artifact: The mismapping of sequencing reads from autosomes to sex chromosomes or vice versa^53,60,61^. We identified and excluded sites which are potentially vulnerable to this artifact (**Supplementary Materials**). In **Fig. 7D**, we only show results for gnomAD subsamples that are the closest in their genetic ancestry to our UKB sample^62^ (results for other subsamples are shown in **Fig. S20,21**). Furthermore, given the concerns of study recruitment biases ^43,60^, we place higher confidence in results that replicate qualitatively across different subsamples, even though we note that subsample-specific selection signals may be real since sexually antagonistic selection may act heterogeneously across groups.

With these conservative criteria considered, we only find evidence for sexually antagonistic polygenic selection on testosterone. In the non-Finnish sample, the largest of the three samples, the null hypothesis *H*_0_:*A* = 0 in **Eq. 1** is rejected (p-value < 0.05) only for testosterone (Z score = 2.2). Testosterone is among the three strongest signals in the two other samples as well, though none of the traits are statistically significant in these samples.

## Discussion

Departing from previous studies that sought GxSex through single loci or heritability analyses, we modelled GxSex as a mixture of polygenic relationships across the genome. Our analysis supports pervasive context-dependency of genetic effects on complex traits, acting largely through amplification. Surprisingly, even for some traits such as red blood cell count, previously considered non-sex-specific because of high genetic correlations between sexes and a concordance in top GWAS hits, we find evidence for substantial GxSex. The strong relationships we find between amplification, environmental variance and phenotypic variance further points to its potential importance for sex differences.

We have shown that considering the polygenic covariance structure, including amplification signals, improves phenotypic prediction for most traits. Its incorporation in polygenic scores is straightforward. We therefore recommend its broad application and further building on our approach to improve clinical risk stratification and other applications of polygenic scores.

Our findings may seem at odds with previous reports of GxSex primarily consisting of sex-limited effects (i.e., no effect in one of the sexes) or antagonistic effects (differences in sign)^63^. In the **Supplementary Materials** and **Table S6**, we illustrate that these apparent discrepancies may be rooted in ascertainment biases. Therefore, limiting analyses to variants with outsized sex differences provides a clouded picture of polygenic GxSex.

Localization of GxSex signals can provide clues into the modulators underlying amplification. Here, we proposed one such modulator, testosterone, and found a correlation between testosterone levels and the magnitude of genetic effect on whole body fat mass. The opposite signs of these correlations in females and males may reflect the discrepant relationship between testosterone and these traits at the phenotypic level.

Our approach for studying GxSex in complex physiological traits can be adopted to study the moderation of polygenic effects by other environments. Starting out with sex as an environmental variable offers a methodological advantage. The study of context-dependency in humans is often complicated by study participation biases, leading to genetic ancestry structure that confounds genotype-phenotype associations^43,64–66^, reverse causality between the phenotype and environment variable, collider bias, gene-by-environment correlation and other problems^67–69^. Focusing on sex as a case study circumvents many of these “usual suspects” problems: For example, problems involving the phenotype causally affecting sex are unlikely. This is an important benchmark for future studies of environmental modulation, both because of the methodological advantage of sex as an environmental variable and because sex is almost always measured; so insight into sex differences in genetic effects can be incorporated straightforwardly in future studies and in clinical risk prediction. Here, we showed that for most of the traits considered, modeling polygenic GxSex (as opposed to individually estimating sex-specific effects at each site; **Fig. S13** yields sex-specific predictors that outperform standard additive polygenic scores.

Finally, we developed a model to consider how GxSex may fuel sexually antagonistic selection in contemporary populations. Over long evolutionary timescales, the two scenarios depicted in **Fig. 7A,B** may lead to different predictions about the long-term maintenance of GxSex genetic variance. Regardless, in both cases, alleles that underlie GxSex may experience sexually antagonistic selection.

We found suggestive signals of sexually antagonistic selection on variation associated with testosterone levels (also see related results by Ruzicka et al.^56^). The signal for our inference of selection is systematic allele frequency differences between adult males and females, which are consistent with contemporary viability selection. The severity, age of onset and prevalence of nearly all diseases are sexually dimorphic^70^. These signals may therefore point to a related disease that differentially affects lifespan in the two sexes, such as immune system suppression, diabetes, cancers, and hypertension^71–74^. Recently, high testosterone levels have been linked to increased rates of mortality and cancer in women, but decreased rates in men^75,76^. However, the testosterone result is also consistent with other accounts, such as testosterone having opposing effects on propensity to participate in a study in the two sexes. Further validation is therefore required to better test hypotheses of sexually antagonistic selection, for example in studies with no recruitment biases (or at least distinct recruitment biases).

In this work, we have shown that amplification of the magnitude of polygenic effects may be important to consider as a driver of sex differences and their evolution. Our approach included the flexible modelling of genetic effect covariance among the sexes, as well as various subsequent analyses exploring the implications of these covariance structures. We hope this study can inform future work on the context-specificity of genetic effects on complex traits.

## Limitations of the Study

Study participation in large biobanks like the UK Biobank (UKB) differs by sex^77^; and work by Piratsu et al. further argued that allele frequency differences between males and females may reflect sex-specific recruitment biases^60^. However, a recent study by Benonisdottir and Kong found no evidence of sex-specific genetic associations with UKB participation^43^, and another by Kasimatis et al. showed that many apparent associations of autosomal genotypes and biological sex in the UKB were instead primarily due to a bioinformatic artifact—the mis-hybridization of autosomal genotyping probes with sex chromosomes^53^. Even still, subtle recruitment biases affecting male and female participation differently remain a possible caveat to our conclusions. For the analysis of natural selection in particular, while the replication of signals of selection in multiple samples may lend credence to our inference, medical datasets based on recruitment of participants via referring physicians, participation biases may still plausibly be shared across studies.

## Acknowledgments

We thank the Harpak Lab and Edge Lab members, Ziyue Gao, Tom Juenger, Jonathan Pritchard, Molly Przeworski, Guy Sella, Jeff Spence and Elliot Tucker-Drob for helpful comments on the manuscript. We also thank Brian Dilkes, Andrés Bendesky and Jim Fleet for insightful discussions. We thank Abin Abraham and Tony Capra for their help in implementing code from Kasimatis et al.^53^, and Michelle Traglia and Lauren Weiss for useful discussions of the relationship between the results reported here and Traglia et al.^63^. This work was supported by NIH GM116853-07 to M.K. and NIH GM137758 to M.D.E. This study has been conducted using the UK Biobank resource under application Number 61666, as approved by the University of Texas at Austin Institutional Review Board, protocol 2019-02-0125.

## Author Contributions

C.Z., M.J.M. and A.H. designed the experiments. C.Z. and M.J.M. performed the experiments. C.Z., M.J.M. and A.H. wrote the paper with assistance from all authors. J.M.C., M.D.E and M.K. provided expertise and feedback.

## Methods

### RESOURCE AVAILABILITY

#### Lead contact

Further information and requests for resources should be directed to and will be fulfilled by the lead contact, Arbel Harpak

#### Materials availability

This study did not generate new unique reagents.

#### Data and code availability

This study used genotype and phenotype data from the UK Biobank https://www.ukbiobank.ac.uk/.

Sex-specific GWAS summary statistics are available at [Zenodo](https://doi.org/10.5281/zenodo.7222725) and are publicly available as of the data of publication. DOIs are listed in the key resources table.

All original code has been deposited at https://github.com/harpak-lab/amplification_gxsex and is publicly available as of the date of publication. DOIs are listed in the key resources table.

Any additional information required to reanalyze the data reported in this paper is available from the lead contact upon request.

### METHOD DETAILS

#### UK Biobank sample characteristics

The UK Biobank is an extensive database that contains a wide variety of phenotypic and genotypic information of around half a million participants aged 40-69 at recruitment^78^.

In this study, we considered 337,111 individuals who passed quality control (QC) checks, which included the removal of samples identified by the UK Biobank with sex chromosome aneuploidy or self-reported sex differing from sex determined from genotyping analysis. We excluded related individuals (3^rd^-degree relatives or closer) as identified by the UK Biobank in data field 22020. To reduce potential population structure confounding, we further limited our sample to individuals identified by the UK Biobank as “White British” in data field 22006. These are individuals who both self-identified as White and as British and were additionally very tightly clustered in the genetic principal component space^78,79^. Individuals who had withdrawn from the UK Biobank by the time of this study were removed. For each phenotype, we also removed individuals who had missing data for the specified phenotype. These procedures left us with between 255,426 to 336,551 individuals in the analysis for each trait.

#### Expectations for sex-specific heritabilities with no GxSex

In the section “The limited scope of analyzing GxSex via heritability differences and genetic correlations,” we report our observation that, for most traits examined, sex-specific heritabilities (i.e., estimated independently from sex-stratified GWAS) were both higher than the heritability in the combined sample. Here, we explain why this observation is inconsistent with a simple model in which genetic effects are the same across the sexes.

Under a simple additive model of variance in a trait *Y* within each sex *Z*,

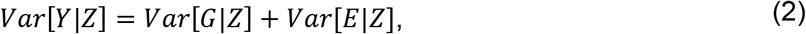

where *Y, G, E* represent the trait value, additive effect, and environmental effect (including all non-genetic context aside from sex), respectively. Under this model, the sex-specific heritability 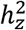 is

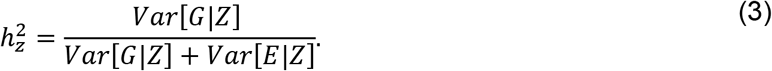

Therefore, sex differences in heritability are either due to sex differences in genetic variance, in environmental variance, or both. If genetic effects are equal, differences in environmental variance alone could cause heritability differences (**Table 1**, first model). But as we show below, the heritability in the combined sample cannot be smaller than both sex-specific heritabilities.

We assume as before that allele frequencies are highly similar between males and females. Since genetic effects are equal, this implies

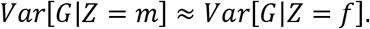

For the environmental variance, we have that

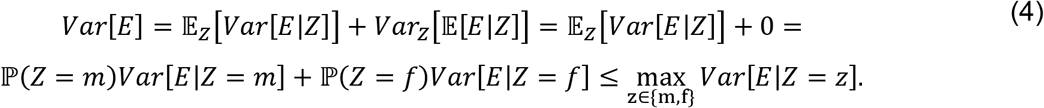

The first equality follows from the law of total variance. In the second equality, we have assumed that there are no mean sex differences in the environmental effects (or, in practice in our analysis and as routine in other analyses, that mean phenotypic sex differences have been subtracted out), giving

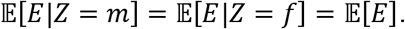

**Eq. 4** shows that the combined environmental variance cannot be greater than the larger of the two sex-specific environmental variances. It follows that if the genetic variance is equal in both sexes, then the heritability in the combined sample cannot be smaller than both of the sex-specific heritabilities,

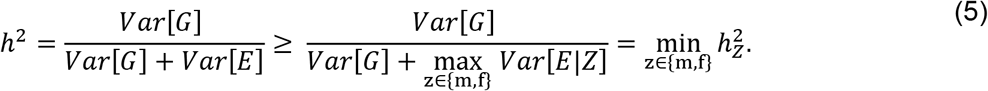

#### Multivariate adaptive shrinkage (mash)

We used multivariate adaptive shrinkage (*mash*) to examine correlation and differences in magnitude of SNP effects between males and females ^45^. *mash* is an adaptive shrinkage method^80^ that improves upon previous methods of estimating and comparing effects across multiple conditions by flexibly allowing for a mixture of effect covariance patterns between conditions and requiring only summary statistics from each condition (including a point estimate of the effect and corresponding standard error for each SNP and condition). The method adapts to patterns of sparsity, sharing, and correlation among the conditions to compute improved effect estimates.

In this study, we set two conditions, male and female, and provided effect estimates and corresponding standard errors from our male-specific and female-specific GWAS. *mash* learns from the data by estimating mixture proportions of various predefined covariance matrices representing different patterns in effects. Using maximum likelihood, mash assigns low weights to matrices that capture fewer patterns in the data, and higher weights to those that capture more.

#### Mixture weights for covariance structure between male and female effects

To interpret patterns of SNP effects between males and females, we inputted 66 hypothesis-based covariance matrices (**Fig. S2**) spanning a range of correlations and relative magnitudes of effects between males and females. We used a random subset of all SNPs for mash to learn the covariance mixture weights. In order for the random subset to contain approximately independent SNPs and capture the weight of SNPs with no effect (**Fig. S2**), we created a subset of SNPs for each trait by taking a random SNP from each of 1703 approximately independent LD blocks estimated for Europeans^81^. *mash* can also generate data-driven covariance matrices that capture SNP effects in the data, but we did not use this feature since the data-driven matrices had negligible differences from our hypothesized matrices (in terms of ℓ2 norm) and were less interpretable.

For each trait, we repeat this weight-learning step 100 times, sampling the SNPs from the 1703 LD blocks without replacement to fit the mash model and generate mixture proportions. We then take the average proportion for each covariance matrix as an estimate of its weight, effectively treating each of the 100 samples as i.i.d. draws.

#### Choice of SNPs used to estimate male-female effect covariance

We examined the effect of using a random subset taken from different p-value thresholds [1, 5e-2, 1e-5, 5e-8] while selecting from LD blocks. By doing so, we can examine differences in the distribution of weights across the p-value thresholds. We performed this test on height, BMI, testosterone, and BMI-adjusted waist:hip ratio. For each trait, weight placed on the no-effect matrix decreased as we reduced the p-value threshold (**Fig. S4A**). Patterns of weights for non-null effect matrices varied across the traits (**Fig. S4B,C**). Since *mash* considers the proportion of null effects and sex-specific, SNP-specific noise; together with the fact that for complex traits, less significant associations may still reflect valuable signal, we decided on using the whole set of SNPs to sample from when estimating mixture proportions.

#### Simulating equal genetic effects and heterogeneous estimation noise among the sexes

To ensure that *mash* was not mistaking sex differences in estimation noise (e.g. via differences in the extent of environmental variance) to be differences in the magnitude of genetic effects, we performed a simulation study. In short, samples of males and females were generated under the model given by **Eq. 2**. Genetic effects were set as equal, but the environmental variance differed among the sexes. We then perform a GWAS on both samples and input the simulated GWAS results into *mash*, and test whether the estimated mixture weights spuriously suggest the presence of GxSex. We performed this simulation on a grid of parameters, including heritabilities in males set to either 5% or 50%, female to male environmental variance ratio of 1, 1.5 or 5; and 100, 1,000 or 10,000 causal SNPs.

First, we created a sample of 300K individuals with randomly assigned sex. We then sampled genotypes for all individuals consisting of 20K SNPs by sampling from the observed distribution of allele frequencies from UK Biobank’s imputed data^82^, assuming linkage equilibrium. From the 20K SNPs, we portioned out the predetermined number of causal SNPs and assigned effect sizes by sampling from a Standard Normal distribution. We estimated the male environmental variance for each causal SNP using the equation,

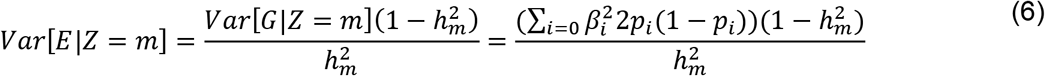

where *Var*[*E*||*Z* = *m*] is the simulated environmental variance for males, *G*|*Z* = *m* is a vector of the genetic effects in males, 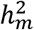 is the heritability in males and *β_i_* and *p_i_* are the effect size and allele frequency at site *i*, which are equal for males and females. We multiplied *Var*[*E*||*Z* = *m*] by the predetermined environmental variance ratio to obtain the environmental variance for females *Var*[*E*||*Z* = *f*]. Afterwards, for each individual *j* with sex *z_j_*, we sampled the environmental effect *E_j_* as

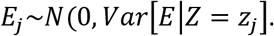

Phenotypes were then set using the following additive model,

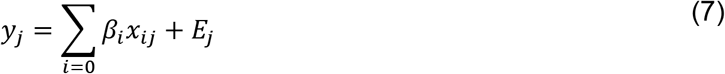

where *y_j_* is the phenotypic value for individual *j* and *x_ij_* is the number of effect allele copies at the *i^th^* causal SNP for the *j^th^* individual. With the phenotype, genotype and environmental effect set, we obtained the estimated effect sizes, 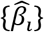, using least squares simple linear regression for all 20K SNPs and used the estimated effect sizes and corresponding standard errors as input into *mash*.

For nearly all parameters, out of the weights on matrices other than the null matrix, the vast majority was placed on the matrix for perfect correlation, equal magnitude (**Fig S6**). As the number of causal SNPs increased, the weight on the no-effect covariance matrix decreased accordingly. These results suggest that *mash* was not grossly mistaking differences in environmental variance as amplification.

#### Simulating sex-biased amplification

To evaluate whether *mash* accurately captures sex-biased amplification of genetic effects (a measure we have used in the x-axis of **Fig. 4A,B**), we followed the same simulation procedure described in the Section “Simulating equal genetic effects and heterogeneous estimation noise among the sexes”. However, instead of using equal genetic effects in males and females, we sampled genetic effects from pre-specified covariance matrices (**Fig. S7** left-hand panel). We set the female to male environmental variance ratio as 1.2 and the heritability as 0.5. We generated data from (A) a model in which all genetic effects are sampled from a matrix where male and female effects are equal, (B) a model in which 86% of the genetic effects are sampled from a matrix where effects between the sexes are equal, and 14% of the effects are sampled from a matrix where the female effect size magnitude is 4 times that of males, and (C) a model in which 86% of effects are sampled from a matrix where effects between sexes are equal, and 14% of effects are sampled from a matrix of only female-specific effects. After simulating sex-specific GWAS on the three models, we input the results into *mash* to estimate mixture weights. We repeated this simulation procedure 100 times for each model.

For model (A), the equal effect matrix received 78% of the weight, and the difference between male-larger and female-larger magnitude was 1% (**Fig. S7**). For model (B), 67% of the weight was placed on the matrix for equal effects. The weight difference between male-larger and female-larger magnitude was 13%. In model (C), 69% of the weight was on the matrix for equal effects, and the difference between male-larger and female-larger magnitude was 16%. These simulation results therefore suggest some overestimation of the proportion of SNPs with magnitude differences. However, the measure of “sex-biased amplification” matched that of the pre-specified generative models up to an error of 2%. Therefore, the simulations suggest “sex-biased amplification” is measured accurately in our estimation procedure.

#### Testosterone as an amplifier

We tested a model of testosterone as a modulator of magnitude differences in males and females. We first split individuals by sex and for each sex, created 10 bins of testosterone levels. We adjusted one of the 10 bins to have testosterone levels overlap between males and females. The overlapping testosterone bin was based on fewer individuals (~800) compared to the other bins (~2200). For each trait, each of the sexes, and within each bin, we performed a simple linear regression of trait values to the PGS for the trait (using a PGS based on both-sex summary statistics (**Supplementary Materials**)). We interpret the estimated coefficient for the effect of the PGS as a proxy for the magnitude of polygenic effect. Finally, we summarized the relationship between testosterone level and magnitude of polygenic effect across bins using the Pearson correlation between the two.

To mitigate the possible effects of confounding (of testosterone and magnitude of polygenic effect) or reverse causation (the magnitude of polygenic effect on the focal trait causally affecting testosterone levels) we employed a version of Mendelian Randomization^83,84^ of the same analysis (**Fig. S16**). Namely, we replaced testosterone levels of each individual with their PGS for testosterone. Here, given the near-zero genetic correlation between males and females, we used our sex-specific PGS for each sex; otherwise, the analysis is unchanged.

We also examined whether participants’ age may have confounded the relationship between testosterone and polygenic effect. In this analysis, instead of using the polygenic effect as the response variable across bins, we used the polygenic effect residualized for mean age in the bin and examined the effect of an individual’s polygenic score on the residual (**Fig. S17**).

#### Model of Shared Amplification

Here, we suggest a null model in which amplification is shared between genetic and environmental effects. We then suggest a prediction that the model yields and explain how we tested this prediction across traits (**Fig. 6**).

If an amplifier is shared, it may be modeled as having the same scalar multiplier effect on genetic and environmental effects. Consider the within-sex additive model of **Eq. 1** in the section “The limited scope of analyzing GxSex via heritability differences and genetic correlations” above. For a phenotype value *Y_z_* in sex *z* ∈ {m, f}

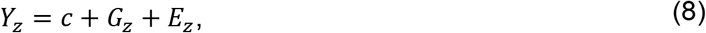

Where *c* is a constant, *E_z_* is the environmental effect and

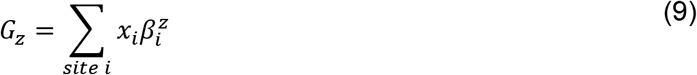

is the polygenic effect where 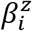 is the effect of an allele at site *i* (say the minor allele) in sex *Z* and *x_i_* is the number of copies of the allele. We assume here for simplicity that male genetic effects relate to female effects solely through a shared polygenic amplification constant, α,

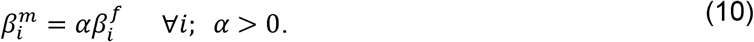

Allele frequencies are once again assumed to be close to equal between males and females, since due to random segregation of alleles during meiosis, genotype frequencies at autosomal sites are independent of sex; and further assuming no substantial interaction between genotype and sex affecting participation in UKB^43^. Consequently, differences in polygenic effect distributions between males and females are solely based on GxSex, and thus:

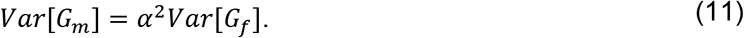

The model we would like to test is one where the amplification of environmental effects can also be simplified to the same scalar multiplier,

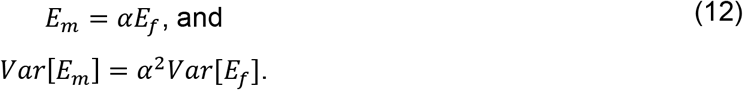

Hence, with equal amplification,

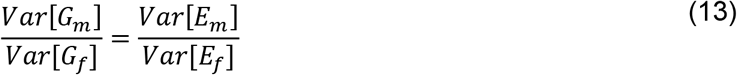

To test the model of shared amplification between environmental and polygenic effects (**Eq. 8**) we obtained the genetic and environmental variance for males and females based on the following relationships,

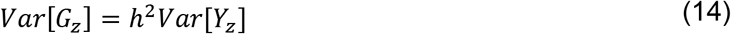

and

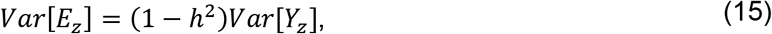

where *Var*[*G_z_*], *Var*[*E_z_*], and *Var*[*G_z_*] are the additive genetic, environmental, and phenotype variances, respectively. Estimates of the sex-specific heritabilities, 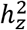, were obtained from previous estimates using LD Score Regression (**Supplementary Materials**).

Representing male genetic or environmental variance as *x*, and the corresponding female variance as y, we derived standard errors for the ratio of male to female variance using the 2 ^nd^-order Taylor approximation for the standard error of a ratio of estimators of *x* and *y*,

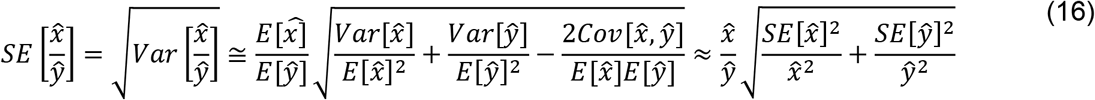

assuming independence between 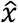 and 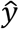 since they are statistics of independent sampling distributions (independent samples of males and females). The standard errors of the genetic and environmental variance were estimated using the law of total variance for a product of two random variables. For 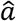 and 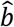, unbiased estimators of the two parameters *a* and *b*, respectively, we get

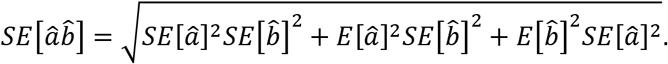

Plugging in the point estimate 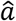 for 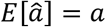 and the point estimate 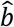 for 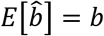,

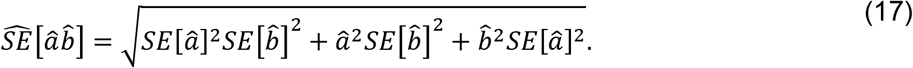

In this case, *a* represents the phenotypic variance for a sex, *Var*[*Y_z_*], and *b* represents either 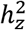 for estimation of genetic variance or 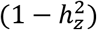 for estimation of environmental variance. Lastly, to obtain the standard error of the phenotypic variance, we used 100 bootstrapped samples *Var*[*Y_z_*]_*i*_ of estimates of the phenotypic variance in sex *z*,

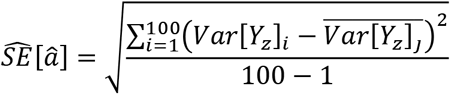

Finally, for each trait, we estimated 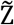, the ratio of the two male-female ratios (environmental and genetic, y and x axes in **Fig. 6**, respectively), and its standard error, 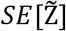, using the same method as in **Eq. 16**. Under the null hypothesis of equal environmental and genetic amplification (**Eq. 8**),

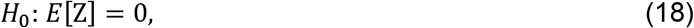

where

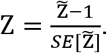

In **Fig. 6**, we approximated 90% confidence intervals on Z by treating it as a Z score, i.e., further treating Z as a Standard Normal.

#### A Model of Sexually antagonistic Selection

We developed a model relating sex differences in additive effects on a trait at a biallelic locus (*β_m_* and *β_f_*) and divergence in allele frequencies. Our model resembles that of Cheng and Kirkpatrick^14^ who developed a similar model relating allele-frequency differences and sex bias in gene expression. In short, we modelled sexually antagonistic, post-conception viability selection on a focal complex trait. We assumed allele frequencies in adult males, *p_m_*, and adult females, *p_f_*, are at equilibrium, i.e. do not change in consecutive generations. Under these conditions, we derive the relationship

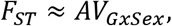

where *F_ST_*^57^ is the fixation index with respect to the male and female subpopulations, i.e., the proportion of heterozygosity in the population that is due to allelic divergence between the sexes. *V_GxSex_* is defined as

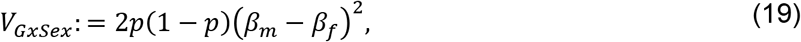

where *p* is the allele frequency in zygotes. *A* is a parameter that, importantly, is shared across all variants affecting the trait and can be thought of as the intensity of sexually antagonistic selection acting on genetic variation for the trait in question.

In our model, allele frequencies at the autosomal locus are assumed to be equal in males and female zygotes. *F_ST_* at adulthood takes the form

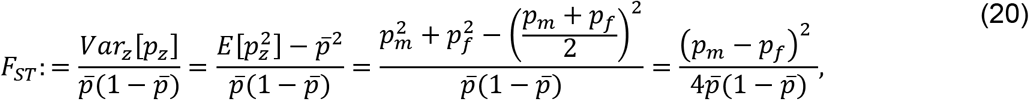

where

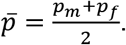

If we further assume a near-1:1 sex ratio such that 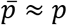,

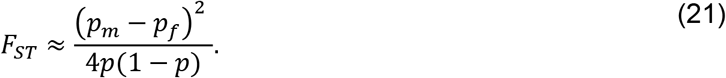

Sexually antagonistic selection acting on viability will cause divergence in allele frequencies between adult males and females. We write the relative viabilities of the homozygote for the reference allele, the heterozygote and the homozygote for the effect allele as 1:: 1 + *d_z_S_z_* :: 1 + *S_z_* for each sex *z* ∈ {*m,f*}. The selection coefficient *S_z_* and dominance coefficient *d_z_* can be frequency-dependent, in which case these coefficients take their values at equilibrium. We can write the additive selection coefficient of the effect allele as

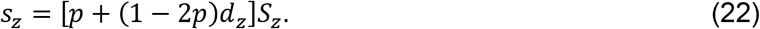

Assuming that zygotes are at Hardy-Weinberg equilibrium, the allele frequency in each sex at adulthood is

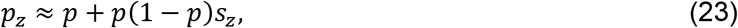

where we neglected terms of order 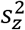 ^85^. Plugging **Eq. 23** into Eq. 21, the divergence between males and females post-selection is

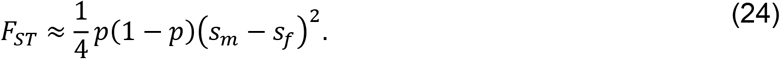

We model the strength of viability selection acting on males and females as linear with the additive effect on a focal trait in each sex,

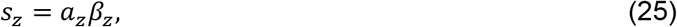

and recalling the simplifying assumption that allele frequencies are at equilibrium under sexually antagonistic viability selection at the locus, such that selection favoring an allele in one sex is balanced by selection against that allele in the other sex,

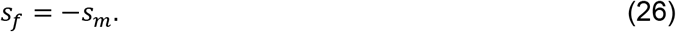

If *β_m_* = *β_f_*, then **Eq. 24** simplifies to

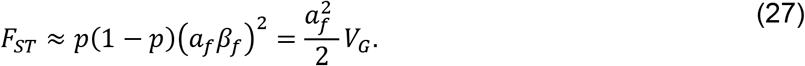

where

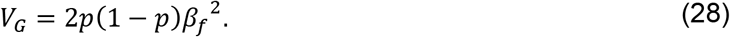

is the additive genetic variance. However, when *β_m_* does not strictly equal *β_f_*, **Eq. 25**, **26** together imply

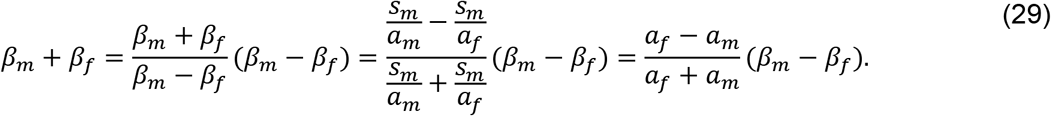

Finally, using **Eq. 25**,

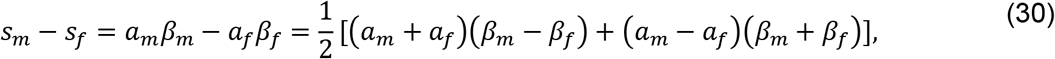

which together with **Eq. 29** gives

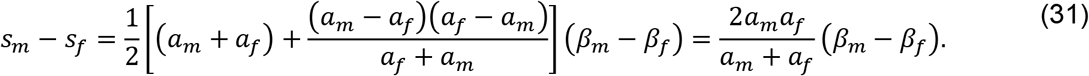

We denote the heritability due to GxSex at the locus as 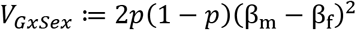 and the parameter relating this contribution to the differentiation in allele frequencies as

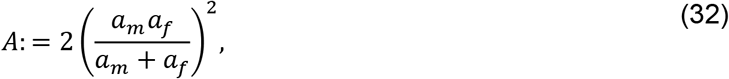

and plug **Eq. 31** into **Eq. 24**, we get

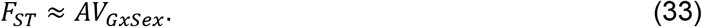

as given by **Eq. 3** in **Results**.

#### Estimating the potential for sexually antagonistic selection on standing variation (A)

For each trait and gnomAD subsample (**Supplementary Materials**), we estimated A using weighted least squares linear regression of our estimate of 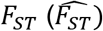 to our estimate of 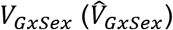, with weight w inversely proportional to our site-specific estimate of noise in the estimate of *V_GxSex_*,

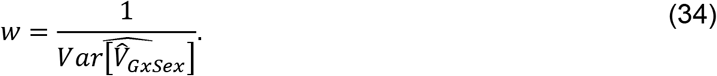

To simplify the estimation of 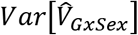, we treated the allele frequency *p* as perfectly estimated, and as independent of the allele frequency in the GWAS sample—as different data are used in the GWAS (UK Biobank) and in the allele frequency estimation (gnomAD). Under these assumptions,

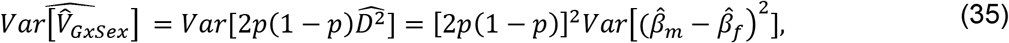

and thus the task at hand is estimating 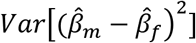. Using the law of total variance,

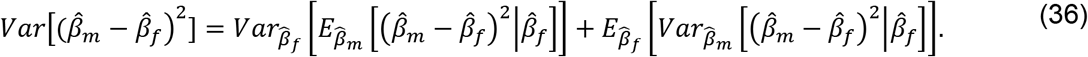

We begin with the argument of the first term,

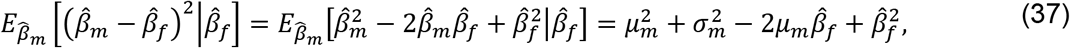

where we denote

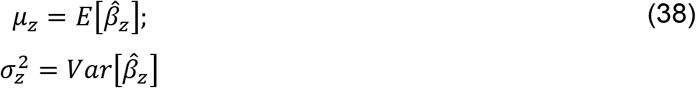

for each sex *z* ∈ {*m,f*}. Plugging **Eq. 37** into the first term of **Eq. 36**,

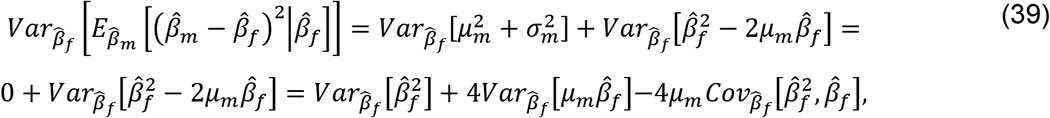

where the first and second step follow from the fact that 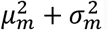 is a constant. We can take note of the fact that 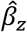 is Normally distributed around *β_z_*, and in particular that it has no skewness. Therefore,

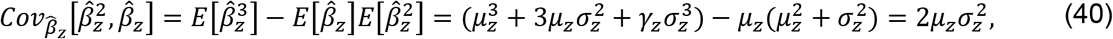

where *γ_z_* = 0 is the skewness of 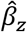. We can also note that

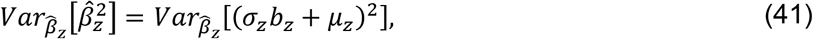

where we defined

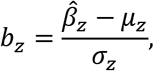

and therefore *b_z_* is a Standard Normal and therefore 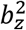 is Chi-squared with one degree of freedom. **Eq. 41** now gives

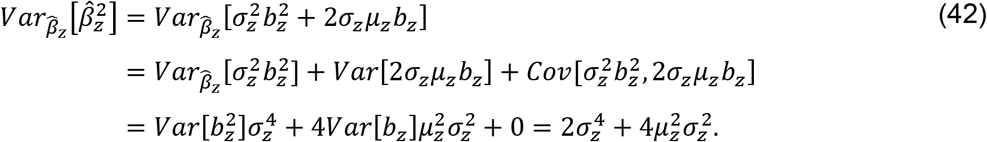

Plugging **Eq. 40** and **Eq. 42** into **Eq. 39**, we find

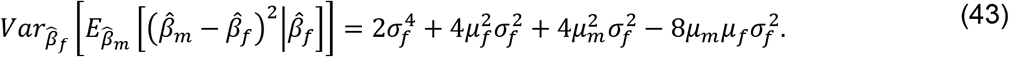

We now turn to the second term of **Eq. 36**. First,

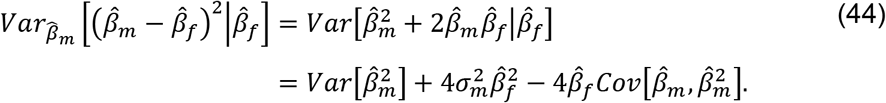

**Eq. 40** and **42** again give us

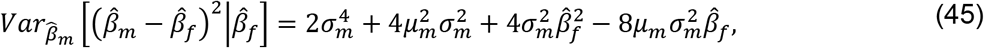

which then gives

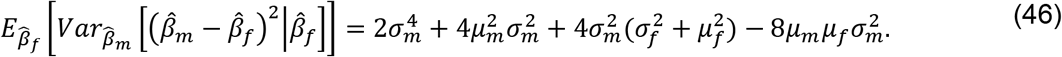

Plugging **Eq. 43** and **Eq. 46** into **Eq. 36**, we obtain

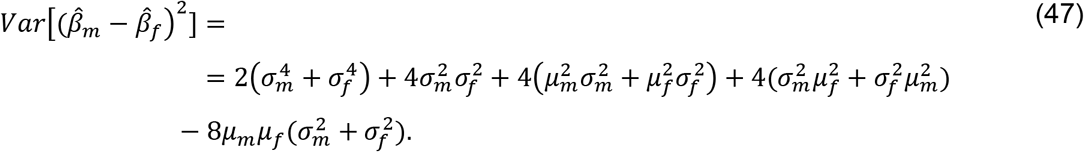

Finally, we estimate *μ_z_* with the GWAS-derived point estimate of the effect 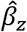 and *σ_z_* with its standard error, 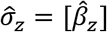. Plugging back into **Eq. 35**, we obtain

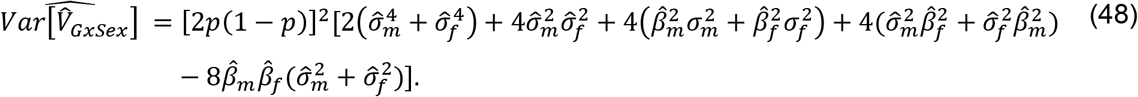

Using **Eq. 33**, we estimate *F_st_* with the estimator

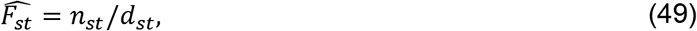

where

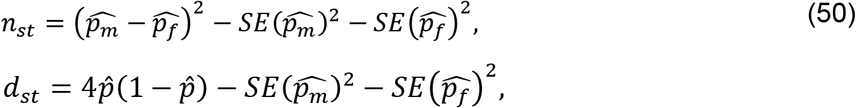

and noting that

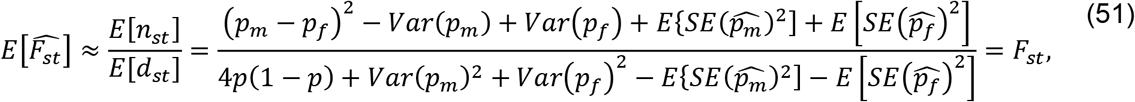

where in the first equality we approximated the expectation of a ratio with the ratio of expectations. Therefore, **Eq. 49** provides an approximately unbiased estimator of *F_st_* despite the absence of genotype frequencies.

To perform this estimation of A on the GWAS and *F_st_* data, we used paired *v* and *V_GxSex_* points for all sites which passed all previous stages of filtering. Weights were set by **Eq. 34** and follow **Eq. 48** where 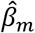 and 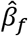 are the GWAS effect estimates as above, and 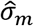 and 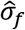 are the GWAS standard errors (SE) estimates for the effect size of each site per trait.

To minimize the possibility of LD between sites used in the analysis as much as possible, we used the approximately independent LD blocks in Europeans^81^ as in Section “Mixture weights for covariance structure between male and female effects”. Namely, we subdivided the genome into 1703 approximately independent LD blocks as before. We iterated over the 1703 blocks and sampling one site per block in a given iteration, using a sample of (up to) 1703 post-filtering sites to perform the weighted linear regression of *F_ST_* on *V_G×Sex_*. The slope of this regression was used as an estimate of *A*. We perform this estimation procedure 1,000 times and take an average of Z scores (slope point estimates divided by their SE) as the final estimate of *A*. In each replicate, we sample with replacement m LD blocks from the m LD blocks which had at least one site within them post-filtering (Supplementary Materials); we then sample one site per resampled block. In **Fig. 7D**, each point is the mean of the 1,000 samples of one site per LD block and 90% confidence intervals show the range between the 5th and 95th percentile of the 1000 replicates.

In the main text, we focus on the results performed this estimation for Ashkenazi Jewish, Finnish, and Non-Finnish European populations as the other ancestry group-stratified subsamples in *gnomAD* are further diverged from the UKB White British sample and therefore our GWAS estimates are expected to be less portable^62,86^. We also performed a similar analysis using UKB data to measure differentiation in allele frequencies between males and females, rather than an independent dataset (*gnomAD*) as in the main text. Since individual level data was available in this case, we replaced *F_st_* with *L_ST_*, a measure developed by Ruzicka et al.^56^. *L_st_* can be thought of as site-specific *F_st_* controlled for major axes of population structure differentiating males and females **(Fig. S21)**.

