## Supplementary Materials for "Amplification is the Primary Mode of Gene-by-Sex Interaction in Complex Human Traits"

Carrie Zhu[1,2], Matthew J. Ming[1,2], Jared M. Cole[1,2], Michael D. Edge[3], Mark Kirkpatrick[2], Arbel Harpak[1,2,+]
[1] Department of Population Health, The University of Texas at Austin, Austin, TX
[2] Department of Integrative Biology, The University of Texas at Austin, Austin, TX
[3] Department of Quantitative and Computational Biology, University of Southern California, Los Angeles, CA

### Supplementary Text

##### Text S1: Genotype data

We focused on autosomal bi-allelic SNPs with an INFO score greater than 0.8. Using *plink 2.0 alpha*[1–6], we further retained variants with calling rate > 0.95, Hardy-Weinberg equilibrium test p-value > 10-9, minor allele frequency (MAF) > 0.001 and MAF < 0.999, following a QC procedure used by the Neale Lab on version 2 of UK Biobank[7]. These were computed on the aforementioned quality-controlled sample set (**Methods "UK Biobank sample characteristics"**) and resulted in 9,607,691 SNPs.

##### Text S2: Phenotype data

Our analysis consisted of 27 continuous traits for their relatively high SNP heritability estimates, based on LD Score regression[7,8]. BMI-adjusted waist:hip ratio (WHR) was calculated by regressing WHR on BMI and obtaining the residuals, as obtained using the following commands in R:

```
model <- lm(<WHR>~<BMI>, data=<dataframe>)
residuals <- summary(model)$residuals
```

##### Text S3: GWAS

We performed all GWAS using *plink 2.0 alpha*, adjusting for birth year, sex, and the first 10 principal components (PC) provided by the UK Biobank as covariates. Covariates were standardized to mean 0, variance 1 (using the flag --covar-variance-standardize). We generated sex-specific GWAS summary statistics for each trait by separating the sample by males and females and applying the same regression model for each sex independently. Any variants with missing values in the summary statistics were removed from further analysis.

##### Text S4: Miami plots

We used the *Ensembl Variant Effect Predictor (VEP)*[9] based on the GRCh38 genome build to annotate SNPs with p-value < $5x10^{-8}$ in Miami plots (**Figs S1**). Using the --nearest flag, we retrieved the gene with the closest protein-coding transcription start site within 5,000 bp up- and down-stream each SNP.

**Text S5: Heritability and genetic correlation estimation using LD Score Regression**

We estimated the SNP heritability of each trait for both-sex, female-specific, and male-specific GWAS, as well as the genetic correlation between sexes using LD Score Regression[8,10]. Since our GWAS summary statistics were based on “White British” individuals of primarily European ancestry, we used the precalculated LD scores computed by Bulik-Sullivan et al.[8].

**Text S6: Qualitative differences between our conclusions and approaches based on independent analysis of individual sex-heterogenous SNPs**

A common approach for detecting and characterizing GxSex based on sex-stratified GWAS data is to test the hypothesis of sex differences in genetic effects at each site independently. As an example, Traglia et al.[11] meta-analyzed sex-stratified GWAS from various sources, many of which standardized effects within-sex. Traglia et al. defined sex-heterogeneous (“sex-het”) SNPs as ones where a t-test testing a null hypothesis of equal effects in males and females was significant at a level of 0.05. They then characterized the pervasiveness of different modes of GxSex focusing on this subset. In particular, they categorized sex-het SNPs where the marginal association p-value was significant at a level of 0.05 as “having an effect in only one sex”. They then categorized the remaining sex-het SNPs as having opposite signs of effects in the two sexes or the same sign and different magnitudes. With these categorizations at hand, they argued that the vast majority of sex-het SNPs have an effect in only one sex.

In this study, we analyzed the polygenic covariance of genetic effects genome-wide, rather than focusing on significant individual SNPs. We proposed that sex differences were largely due to differences in magnitude rather than opposite or sex-private effects. Here, we show that the seemingly discrepant arguments may be a direct result of the different analysis approaches. We show that when we generate data from a generative model adhering to pervasive amplification and use Traglia’s et al.’s approach to characterize GxSex based on the simulated data, their classification suggests the same qualitative result—that GxSex manifests primarily via effects that are private to one of the sexes.

We performed a simulation study following a procedure similar to that in the section “**Simulating equal genetic effects and heterogenous estimation noise among sexes**” of the main text. Here, however, we sample genetic effects using the covariance mixtures estimated for five traits – height, BMI, creatinine, IGF-1 and systolic BP (**Fig. S5**). We set the female to male environmental variance ratio as 1.2 and the heritability as 0.05. We simulated genotypes and phenotypic values, and then standardized phenotypic values within-sex and performed a sex-stratified GWAS. Using the effect estimates and corresponding standard errors for males and females, we calculated a t-statistic for each SNP, where

$$t = \frac{\hat{\beta}_{female} - \hat{\beta}_{male}}{\sqrt{SE_{female}^2 + SE_{male}^2}},$$

and identified SNPs with p-value < 0.05 as sex-het SNPs. We further categorized sex-het SNPs as having an effect in only one sex if the effect was only significant in one sex (t-test p-value < 0.05). For SNPs where the sex-specific effect was significant for both sexes or for neither, the SNPs were categorized based on if the effects were of opposite or same sign.

For data generated based on the covariance structures, we indeed find that the classification of sex-het SNPs qualitatively mirrors that of Traglia et al., with the majority of the sex het SNPs categorized as having an effect in just one sex (**Table S6**). Importantly, this is despite having simulated traits with pervasive differences in the magnitude of effects. We therefore conclude that such an approach, including the within-sex standardization of effect sizes (which heavily dilutes signals of pervasive magnitude differences), the ascertainment bias resulting from a focus on individually-significant sex-heterogenous SNPs, and the use of the above-described classification result in mis-characterization of the mode of GxSex, can lead to the mischaracterization of GxSex in complex traits.

**Text S7: Posterior estimates of sex-specific effect sizes**

*mash* can also apply adaptive shrinkage after learning patterns in effect sizes to improve marginal point estimates—estimated using the posterior mode—and measures of significance. These posterior estimates could reduce noise by shrinking effects towards zero and possibly reveal greater or lesser variation in effect sizes between males and females. Therefore, using the average of the fitted mixture model over 100 repetitions (**Methods "Mixture weights for covariance structure between male and female effects"**), we computed posterior estimates for each trait be used in further analysis (**Text S8**). *mash* calculates a local false sign rate (lfsr) for each effect, which is similar to a local false discovery rate and is defined as the cumulative density of the posterior distribution of values with a sign that differs from that of the posterior mean. We mapped lfsr to "pseudo p-values" by ranking the SNPs according to lfsr values, and then ascribing a p-value of the same rank (**Fig. S11**).

**Text S8: Improved utility of sex-specific models for complex phenotype prediction.**

The pervasiveness of GxSex that we infer, alongside the mixture of covariance relationships across the genome for each trait, may mean that GxSex is important to consider in phenotypic prediction. To test this possibility, we compared the prediction accuracy of three polygenic scores (PGS) for various traits, together with covariates (**Fig. S13**):

1. An additive PGS, assuming no GxSex, based on both-sex GWAS summary statistics
2. A sex-specific, but polygenic covariance-naïve PGS, based on stratified GWAS, fit independently for each sex.
3. A sex-specific, covariance-aware PGS based on posterior sex-specific effect estimates (**Text S7**)

The estimation, prediction and evaluation pipeline are illustrated in **Fig. S11**, and, for models (a) and (b), parallels the procedure described by Choi[11].

First, for each phenotype, we split the sample of unrelated "White British" individuals into a test set of 25K individuals randomly sampled from each sex and a training set with the remaining individuals. Second, we re-ran GWAS on the training set following the same procedure above (**Text S3**), generating both-sex, female-specific, and male-specific summary statistics. Third, to obtain posterior estimates, we first input the effect sizes and standard errors from the male-specific and female-specific GWAS, and took the average of 100 resampling estimates of mixture proportion vectors estimated from a random subset of SNPs with p-value < 1e-5 available in each of 1703 LD blocks[12]. We used the 1e-5 p-value threshold to create the random subset to provide stronger signals and patterns for *mash* to learn from. We then feed estimated mixture proportion vectors into *mash* to perform the refined, covariance-aware estimation of effects. The output from *mash* included posterior mean and a local false sign rate (lfsr) for each SNP. We sorted SNPs by lfsr and matched it to a ranked list of p-values from the additive GWAS summary statistics to create a "pseudo p value" which we use later for thresholding.

Afterwards, we performed clumping separately for the additive both-sex, male-specific additive, female-specific additive, male covariance-aware and female covariance aware models to produce a subset of significant SNPs that are approximately independent using *plink 1.9 beta*'s --clump command, removing SNPs with pairwise LD threshold $r^2 > 0.1$ or within 250kb. To estimate pairwise LD values, we used a sample of 187 unrelated individuals (population codes GBR and CEU) from 1000 Genomes phase 3[13].

Using the resulting subset of SNPs, we estimated PGS for the individuals in the test set by summing the number of effect alleles an individual has weighted by the allelic effect sizes. We used *plink 2.0 alpha's* --score command along with the --q-score-range flag to repeat the PGS computation over a range of p-value thresholds [1, 0.01, 1e-5, 1e-8]. Therefore, a total of 20 PGS runs were performed for each trait over the combination of five models and four p-value thresholds (**Fig. S11**).

Finally, to assess the prediction accuracy, we computed $R^2$ for the following,

$$y \sim birth\ year + sex + principal\ componenets + (sex \times M) + ((1 - sex) \times F)$$

$$sex: \{0(female), 1(male)\}$$

$$M: male\ PGS \qquad F: female\ PGS$$

The covariates used for the regression are the same as those used in our GWAS: sex, birth year, and the first 10 PCs of the genotype matrix (UKB data field 22009). We performed regressions both separately by sex and together with both sexes in the test set. For each of the three predictors—additive model, sex-specific additive model, and sex-specific polygenic covariance-aware model—we selected the p-value threshold with the greatest $R^2$. We also calculated the incremental $R^2$, which is the increment in $R^2$ after adding the PGS to a null model with only the covariates (**Fig. S11**).

We performed 20-fold cross validation for our PGS procedure. Each prediction was performed on a different randomly sampled 5K female and 5K male test set, with the remaining used at the training set. For each model, we averaged the greatest incremental $R^2$ from the p-value thresholds over the five folds for comparison across all models.

The sex-specific, covariance-aware PGS model outperformed the additive model for 20/27 traits. The sex-specific, covariance-aware PGS model outperformed the sex-specific, covariance-naïve model for all traits but testosterone. Similarly, the additive model outperformed the sex-specific covariance-naïve for all traits but testosterone.

The increase of prediction accuracy in the additive model over the sex-specific covariance-naïve model may be due the additive model being estimated on nearly twice the sample size as the sex-specific models. We chose not to equalize the sample sizes to align the comparison to what would be used in practice. However, despite the disadvantage in decreased sample size, the covariance-aware PGS model outperformed the additive model in approximately 3/4 of the total traits.

Polygenic scores used in **Methods "Testosterone as an amplifier"** and **Fig. 2** were based on a 5-fold cross validation with 50K individuals in the test set (25K males and 25K females). The polygenic scores were gathered from the first fold for each trait.

##### Text S9: Data Filtering for Sexually-Antagonistic Selection Analysis.

We downloaded allele frequency data from the gnomAD dataset[14]. Specifically, we used gnomAD v3.1.2, which consists of 76,156 whole genome sequences mapped to the GRCh38 reference. In order to allow this assembly to conform to the GRCh37 build used in the UK Biobank GWAS, we used the UCSC in-browser tool LiftOver[15] (https://genome.ucsc.edu/cgi-bin/hgLiftOver) to identify and convert the GRCh38 sites of interest to their corresponding GRCh37 positions. Allele count summaries are available for the total sample, as well as stratified by sex chromosomes karyotype and genetic ancestry groupings: African/African American samples (abbreviated "afr" in the gnomAD files); Amish ("ami"); Latino/Admixed American ("amr"); Ashkenazi Jewish ("asj"); East Asian ("eas"); Finnish ("fin"); Non-Finnish European ("nfe"); Middle Eastern ("mid"); South Asian ("sas"); and samples not assigned to any population are designated Other ("oth"). Aneuploid individuals (e.g., X or XXY) are not included in the dataset. For the purposes of this study, we again refer to XX as female and XY as male. Total numbers of individuals sampled can be found on the gnomAD website's help page (https://gnomad.broadinstitute.org/help).[14]

We downloaded gnomAD VCF files from the gnomAD browser for all autosomes and used VCFTools[16] to parse the file. We filtered the data to exclude insertions or deletions, and only kept bi-allelic SNPs. We further removed missing data (3,698 sites). These filtering steps resulted in 2,285,169 remaining sites. In an effort to avoid confounding results that could arise from population substructure, we split the data into the different ancestry groups labeled by gnomAD and worked with the data in each subpopulation separately from this point forward. We removed sites with less than 1,000 alleles in each ancestry group independently. The number of sites we removed at this step depended on the sample size of each group (**Table S1**). This step resulted in the complete removal of the Amish and Middle Eastern subsamples because their low sample sizes.

Finally, we filtered out sites where difference in male and female allele counts may be partly or fully driven by the mismapping of autosomal reads to sex chromosomes or vice-versa[17,18]. We follow a similar approach to Kasimatis et al. to identify such sites[18]. In particular, for every

SNP, we extract the 301bp sequence surrounding the SNP, with the SNP's position at the center, from the GRCh37 genome assembly[19]. We further shorten the 301bp sequence into three 150bp-long subsequences with the SNP's position at the center, start or end of the sequence. We then use Mega-BLAST through NCBI's command-line BLAST tool[20] to search for regions of high sequence homology to either of the three subsequences. If any of the three was found to have a 90% or greater sequence identity to a sequence on a sex chromosome, we filtered out the site. This filtering was performed agnostic of ancestry.

We qualitatively compared our list of filtered SNPs to Kasimatis et al.[18]. We based our comparison on sites considered in both works. In particular, we limited the comparison to the genotype array sites, as they did not consider UKB imputed genotypes, and further only to sites we had not removed in a previous filtering step. In terms of parameters, our 90% homology threshold is the same as Kasimatis et al., but we diverge from their algorithm in testing three different sequences per SNP; and also in using shorter sequences (150bp) to model for the mapping of short reads rather than the hybridization of probes. In general, our approach tended to identify more sites than Kasimatis et al. as invalid for analysis (**Table S5**).

##### Text S10: Estimating Male-Female $F_{ST}$ and $V_{GxSex}$

We estimated Male-Female $F_{ST}$ for every site remaining in our dataset based on the new estimator we propose in **Eq. 51,** based on the sample allele frequencies. We removed sites for which there were no alternative allele calls for either males or females. This step resulted in a significant reduction in sample size by one or two orders of magnitude, depending on the ancestry (**Table S2**). We further removed sites that were not found in both the gnomAD dataset and the UKB GWAS dataset. The number of sites removed through this step varied greatly across ancestry groups (**Table S3**).

We used the point estimates and the standard errors of the sex-stratified GWAS (**Table S6)** for 27 physiological or physical traits. We further filtered sites by GWAS p-value. We used four different p-value thresholds at $10^{-3}$, $10^{-5}$, $10^{-8}$ and 1 (i.e., all SNPs; see **Table S4** for the number of sites remaining for each p-value threshold). In the main text, we focus on the $10^{-5}$ threshold, reasoning that it strikes a reasonable, albeit arbitrary, middle ground between sample size and noise. Results for other p-value thresholds are shown in **Fig. S20**.

Finally, we obtained an estimate of $V_{GxSex}$ using **Eq. 19**, where $\beta_m^2$ and $\beta_f^2$ were estimated using the squared GWAS effect estimates, and $p$ is the total alternate allele frequency as above. For detail on the estimation of the sampling error of $V_{GxSex}$, see “**Estimating the potential for sexually-antagonistic selection on standing variation ($A$)**” in the main text.

**Text S11: Competing models for sex differences in trait variance**

In the main text section “**Are polygenic and environmental effects jointly amplified**”, we show that under a model where amplification is pervasive and shared between environmental and genetic effects, we expect the male-female ratio of environmental variance to equal the male-female ratio of genetic variance. Here, we compare the expectation under the pervasive, joint amplification model with expectations under two other longstanding models.

As recently discussed by Zajitschek et al.[21] , the ”estrus-mediated variability” hypothesis predicts that females will display higher trait variability in traits affected by the estrous cycle. If we interpret estrus-mediated effects as environmental, this hypothesis suggests equal genetic variance but larger environmental variance in females (**Fig. S19A**; orange line in **Fig. S19B**).

The ”greater male variability” hypothesis predicts that males will display higher trait variability. There are multiple rationales offered for this hypothesis, such as stronger sexual selection on males that leads to higher variability via group selection[22–24]. A more widely applicable rationale for this prediction is that mammalian males, as the heterogametic sex, will experience more variable X chromosome effects, whereas these effects will be “averaged out” through heterogeneous X inactivation in females[23]. This hypothesis therefore predicts greater genetic variance in males and equal environmental variance between males and females (**Fig. S19A**; green line in **Fig. S19B**.

### Supplementary Figures

**Figure S1. Sex-specific Miami plots, related to Figure 3**

“Miami plots”, contrasting statistical significance of marginal effects in a female-specific GWAS (top) and a male-specific GWAS (bottom), for each of the 27 traits analyzed. Data points are thinned out by random selection over multiple levels of p-value thresholds. For larger p-values, more points are not shown.

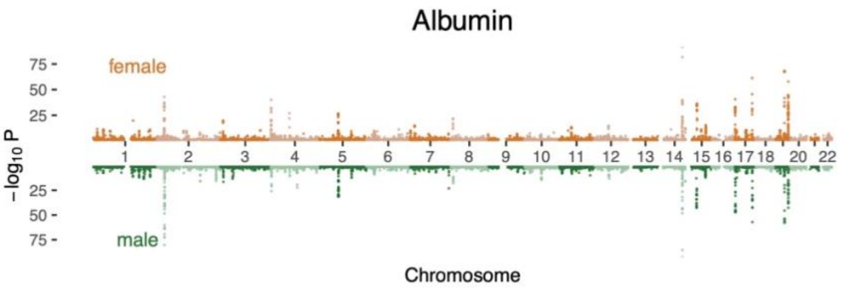

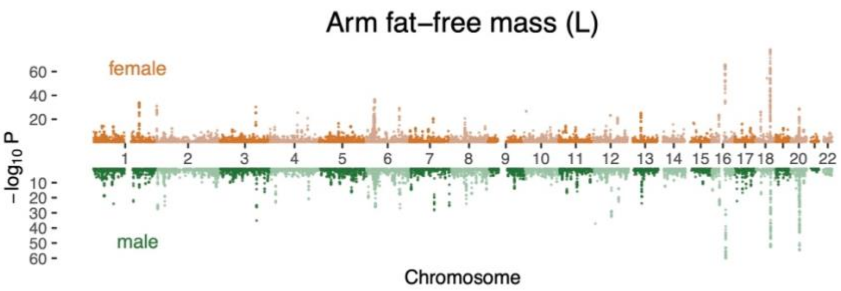

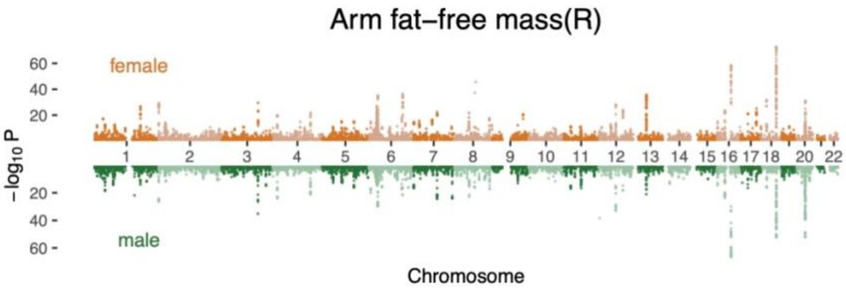

BMI

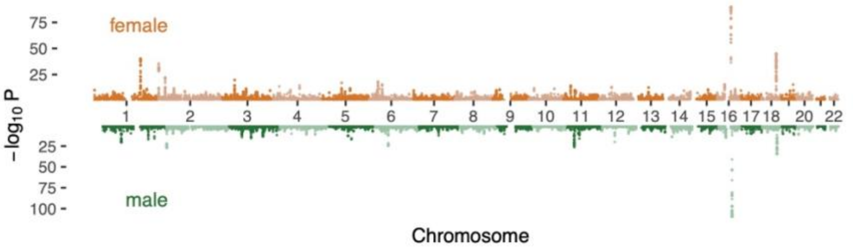

Calcium

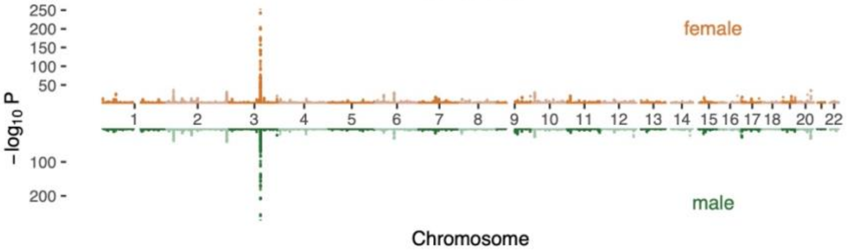

Creatinine

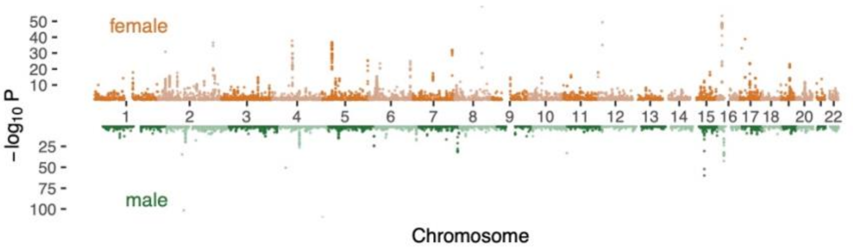

Diastolic BP

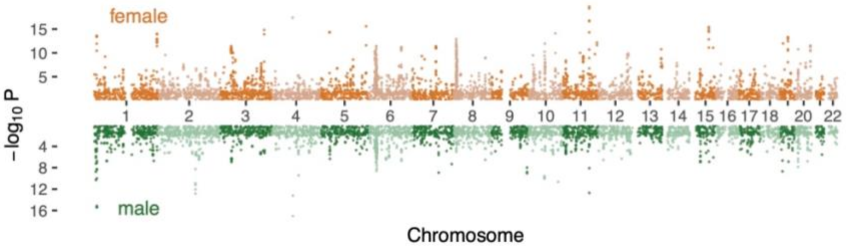

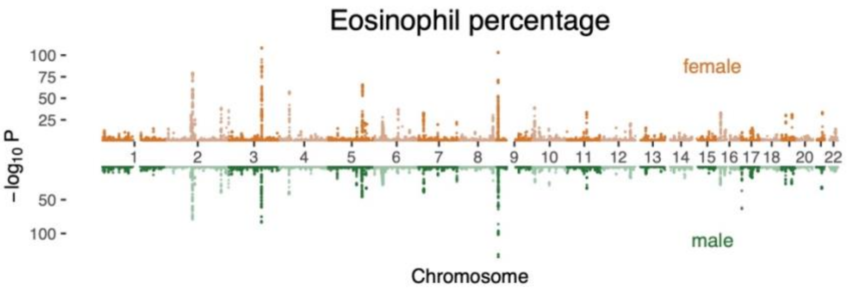
Eosinophil percentage
female
male
−log10 P
Chromosome

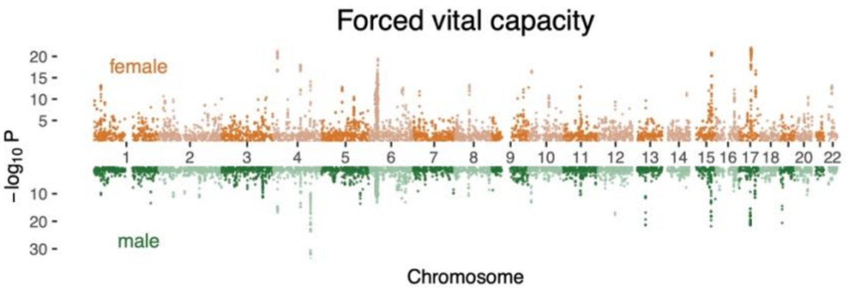
Forced vital capacity
female
male
−log10 P
Chromosome

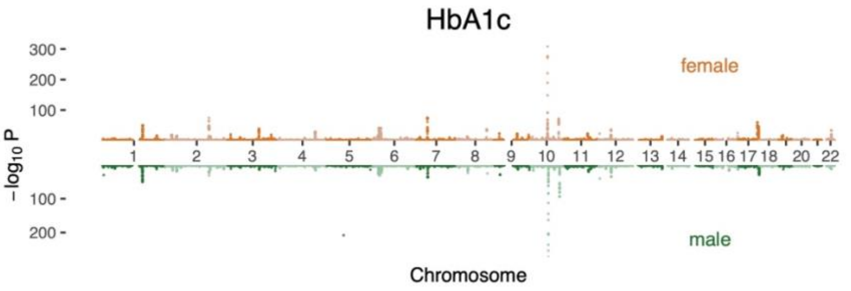
HbA1c
female
male
−log10 P
Chromosome

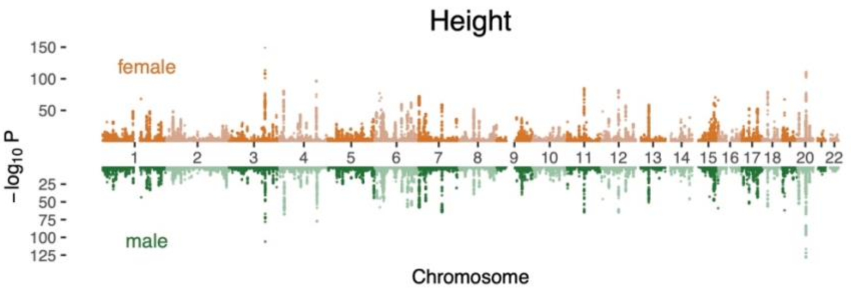
Height
female
male
−log10 P
Chromosome

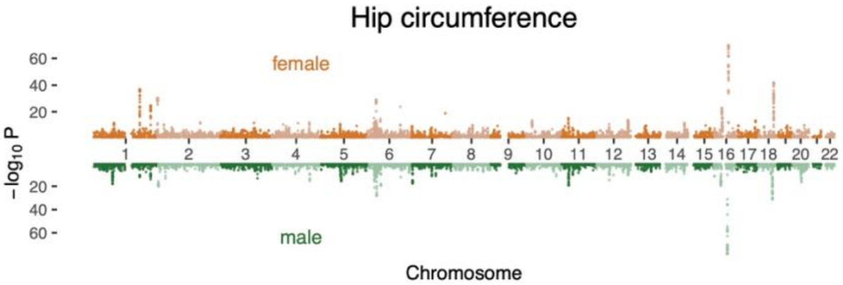
Hip circumference
female
male
−log10 P
60
40
20
20
40
60
1 2 3 4 5 6 7 8 9 10 11 12 13 14 15 16 17 18 20 22
Chromosome

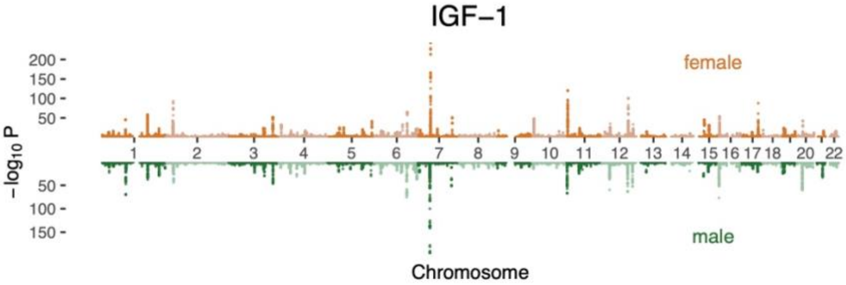
IGF−1
female
male
−log10 P
200
150
100
50
50
100
150
1 2 3 4 5 6 7 8 9 10 11 12 13 14 15 16 17 18 20 22
Chromosome

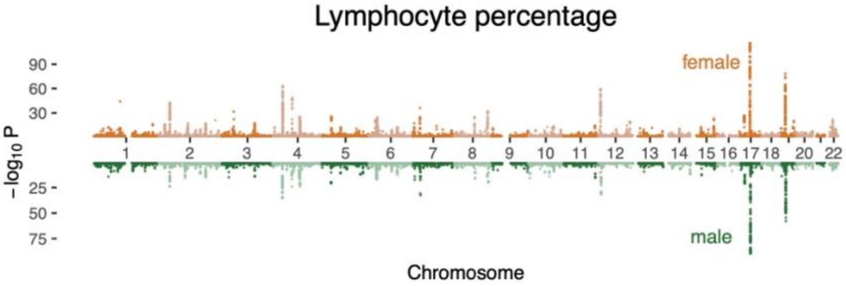
Lymphocyte percentage
female
male
−log10 P
90
60
30
25
50
75
1 2 3 4 5 6 7 8 9 10 11 12 13 14 15 16 17 18 20 22
Chromosome

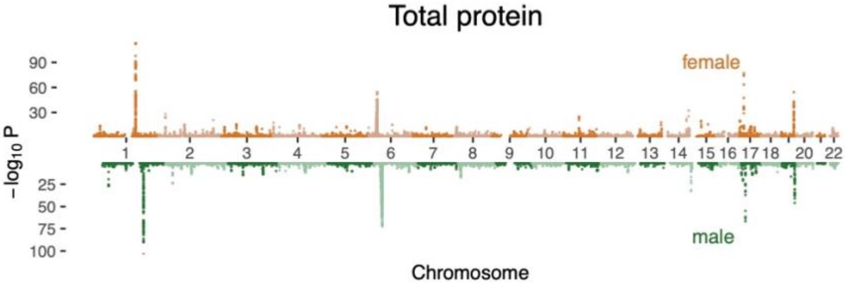
Total protein
female
male
−log10 P
90
60
30
25
50
75
100
1 2 3 4 5 6 7 8 9 10 11 12 13 14 15 16 17 18 20 22
Chromosome

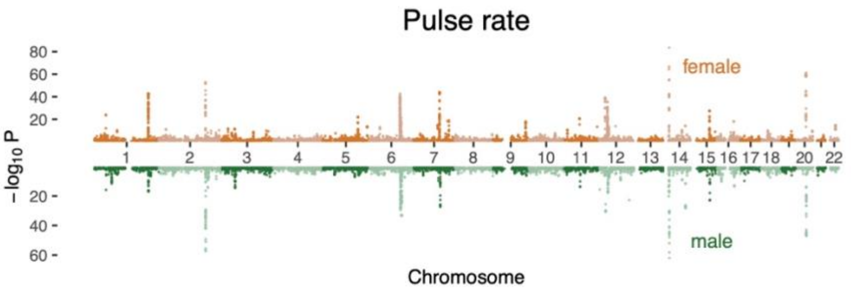
Pulse rate
80
60
40
20
−log10 P
20
40
60
1 2 3 4 5 6 7 8 9 10 11 12 13 14 15 16 17 18 20 22
female
male
Chromosome

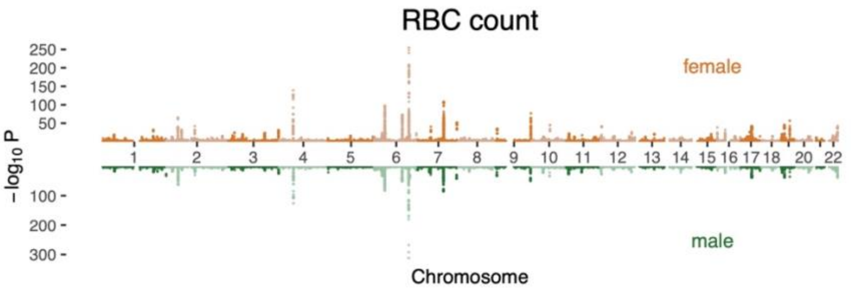
RBC count
250
200
150
100
50
−log10 P
100
200
300
1 2 3 4 5 6 7 8 9 10 11 12 13 14 15 16 17 18 20 22
female
male
Chromosome

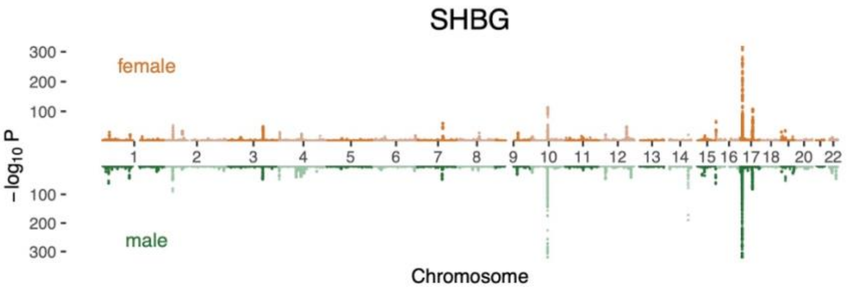
SHBG
300
200
100
−log10 P
100
200
300
1 2 3 4 5 6 7 8 9 10 11 12 13 14 15 16 17 18 20 22
female
male
Chromosome

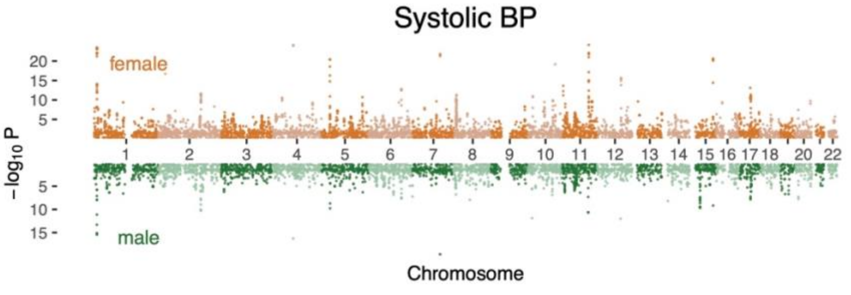
Systolic BP
20
15
10
5
−log10 P
5
10
15
1 2 3 4 5 6 7 8 9 10 11 12 13 14 15 16 17 18 20 22
female
male
Chromosome

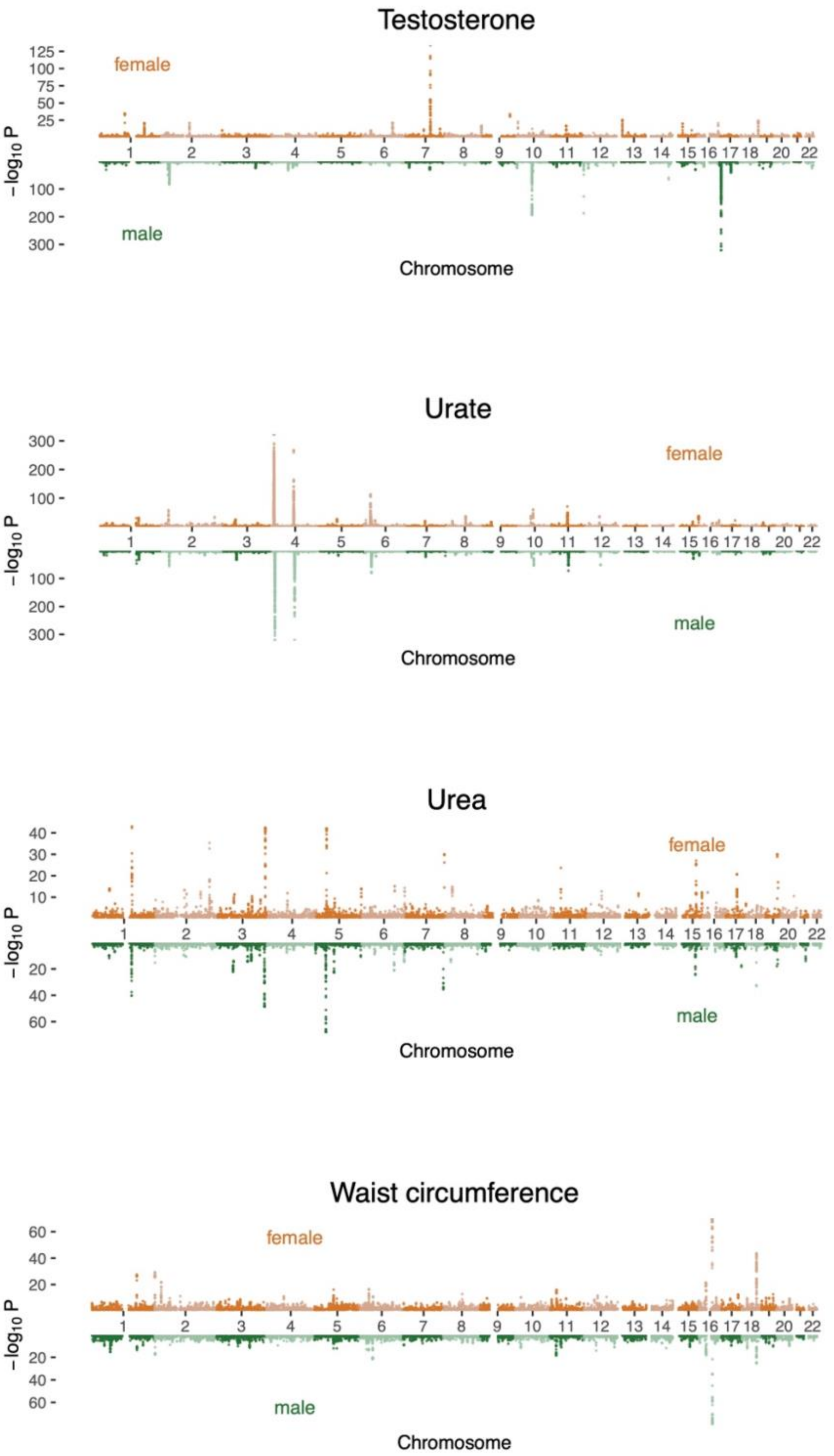
Testosterone
female
male
−log10 P
Chromosome
Urate
Urea
Waist circumference

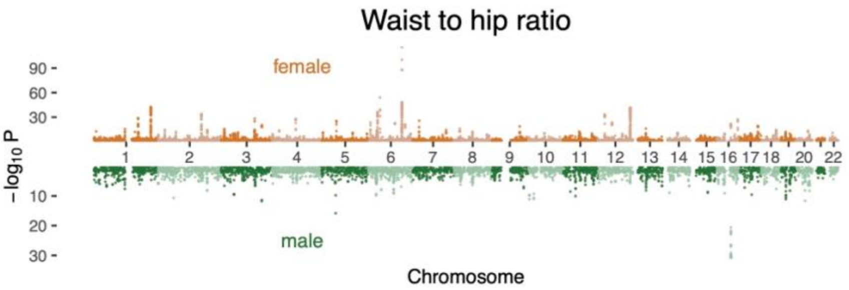
Waist to hip ratio
female
male
−log10 P
90
60
30
10
20
30
1 2 3 4 5 6 7 8 9 10 11 12 13 14 15 16 17 18 20 22
Chromosome

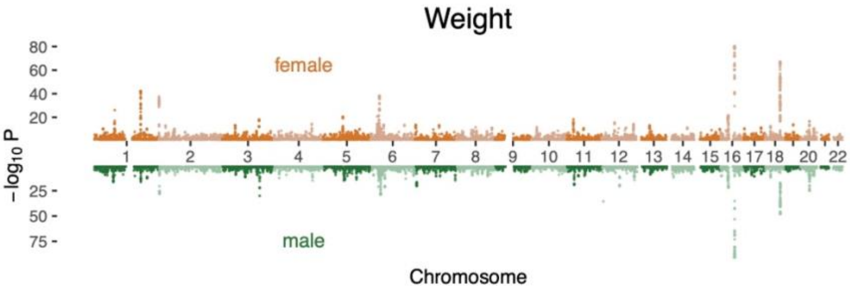
Weight
female
male
−log10 P
80
60
40
20
25
50
75
1 2 3 4 5 6 7 8 9 10 11 12 13 14 15 16 17 18 20 22
Chromosome

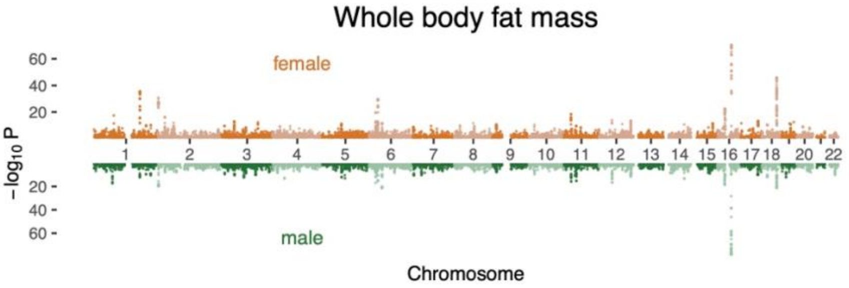
Whole body fat mass
female
male
−log10 P
60
40
20
20
40
60
1 2 3 4 5 6 7 8 9 10 11 12 13 14 15 16 17 18 20 22
Chromosome

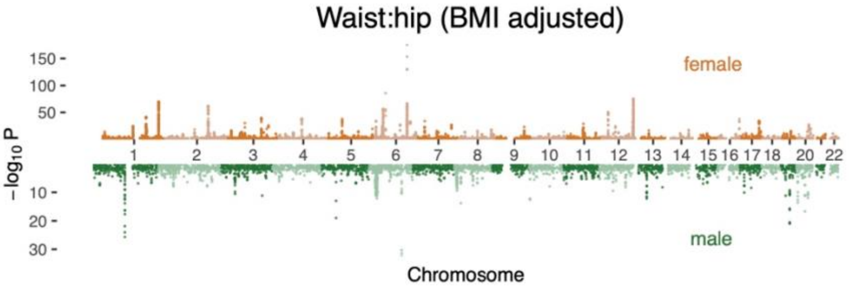
Waist:hip (BMI adjusted)
female
male
−log10 P
150
100
50
10
20
30
1 2 3 4 5 6 7 8 9 10 11 12 13 14 15 16 17 18 20 22
Chromosome

**Figure S2. All hypothesis covariance matrices, related to Figure 3**

All hypothesis matrices we inputted into *mash* are shown here organized by correlation and magnitude of effect size. The placement of the matrices parallels in the weights in **Fig. S5**.

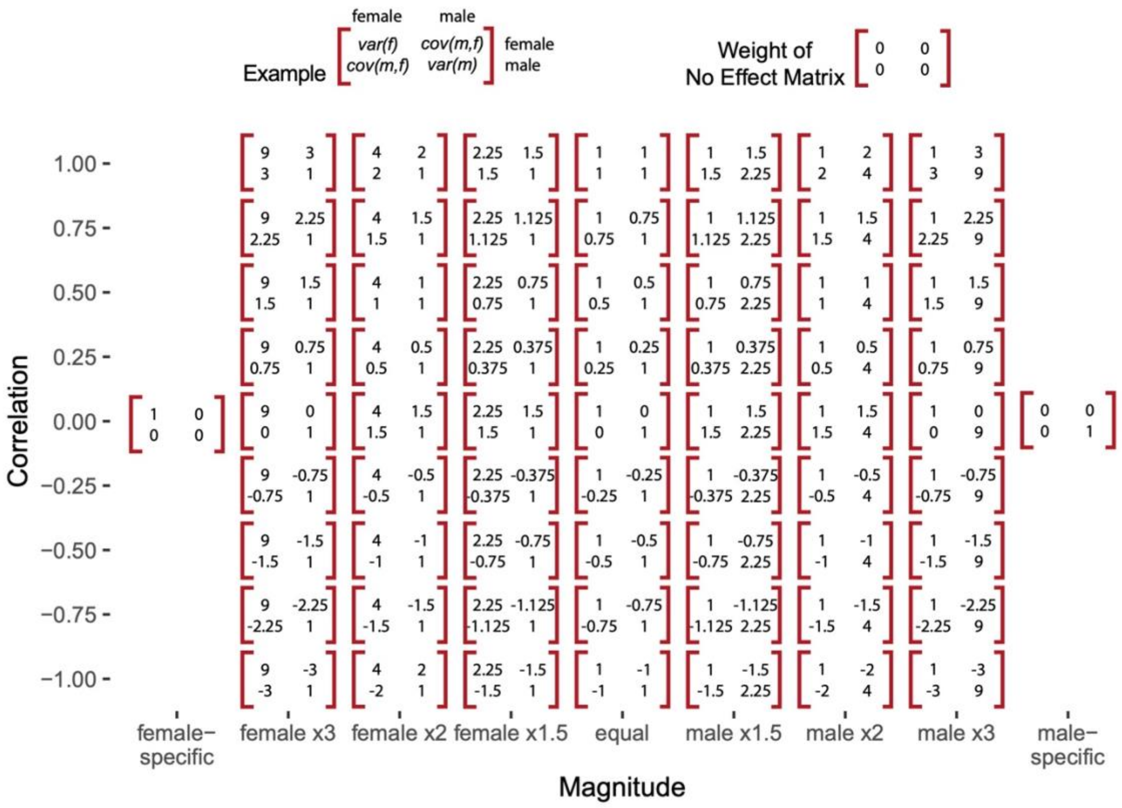

**Figure S3 Inferring polygenic covariance structure, related to Figure 3**

**(A)** Our analysis of the polygenic covariance between males and females is based on sex stratified GWAS. Shown for illustration, is a "Miami plot" for testosterone. Highly associated SNPs that are less than 5 kbp away from a transcription start site were annotated with the gene corresponding to the closest one. **(B)** We modelled the sex-stratified GWAS estimates as sampled with error from true effects arising from a mixture of pre-specified hypothesis covariance relationships between female and male genetic effects; see examples in red frames. Each box specified a mixture weight (±SE) that we infer for one hypothesis matrix. The weight, also indicated by the shade, corresponds to the relative frequency that the specified hypothesis matrix is represented by the variants. The axes state the relative magnitude (amplification) and correlation between males and females, which jointly make up the covariance relationship. **(C)** The x and y axes are a condensed version of the x and y axes from (B) for testosterone. The weights are a proportion of the non-null weights, i.e., the weight divided by sum of all weights except for the weight on the no effect matrix, corresponding to no effects in either sex. For example, the square in purple sums over all 12 weights for matrices corresponding to larger effects on testosterone in males that are negatively correlated with effects in females, 5.1%, divided by the total weight on matrices with nonzero effects, 32%.

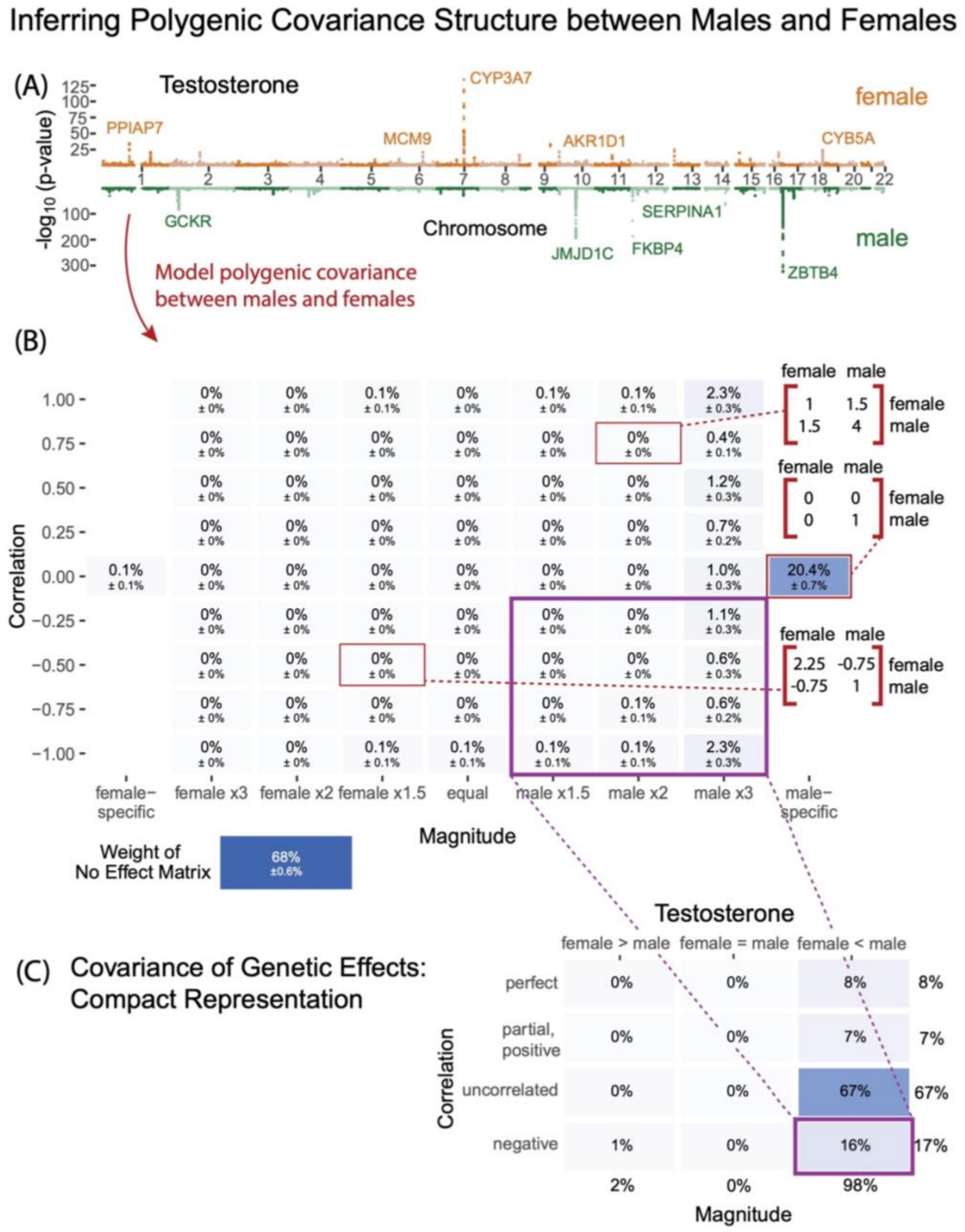

**Figure S4 Testing p-value thresholds for input into *mash*, related to Figure 3**

We tested the effect of inputting random subsets taken from various p-value thresholds [1, 5e-2, 1e-5, 5e-8] in the *mash* fitting step to estimate mixture proportions. The random subsets were drawn by sampling from the 1703 LD blocks (**Methods "Mixture weights for covariance structure between male and female effects"**). **(A)** Both depict the weight on the no-effect matrix per p-value threshold for four different traits. The left plot shows results from sampling once from each available LD block as long as there were still p-values below the specified threshold. For the right plot, we keep sampling from LD blocks until we reach a subset of 1703 SNPs, ensuring equal sample sizes across the thresholds. **(B)** The percentage of weight by correlation of effects are depicted across the thresholds. Plots in the box show results from equal sample sizes. **(C)** The percentage of weight on different types of magnitude of effects are shown for four different traits across the threshold. Plots in boxes also show results from equal sample sizes.

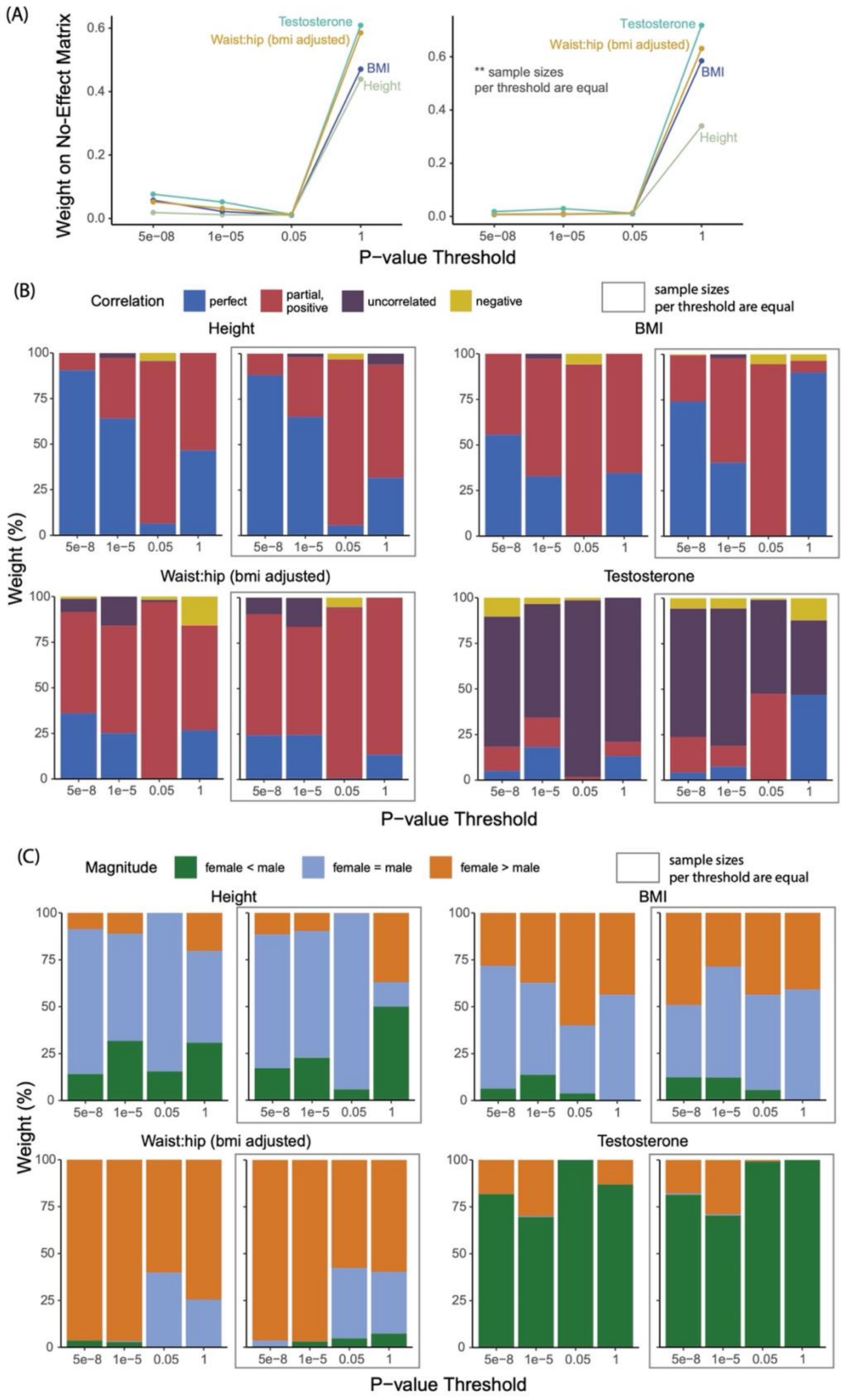
(A)
Weight on No-Effect Matrix
Testosterone
Waist:hip (bmi adjusted)
BMI
Height
** sample sizes
per threshold are equal
P−value Threshold
(B)
Correlation
perfect
partial, positive
uncorrelated
negative
sample sizes
per threshold are equal
Height
BMI
Waist:hip (bmi adjusted)
Testosterone
Weight (%)
P−value Threshold
(C)
Magnitude
female < male
female = male
female > male
sample sizes
per threshold are equal
Height
BMI
Waist:hip (bmi adjusted)
Testosterone
Weight (%)
P−value Threshold

**Figure S5 Mixture weights on hypothesis covariance matrices, related to Figure 3**

Mixture weights estimated from a sample of SNP effects using *mash* are shown with the corresponding hypothesis covariance matrix represented by the axes. Covariance matrices span all hypothesis matrices shown in **Fig. S2**. The “Weight of No Effect Matrix” box represents the weight on a matrix signifying no effects in both sexes. The bottom plot for each traits represents a consolidated version of the top plot. Weights are shown as percentages of non-null weights.

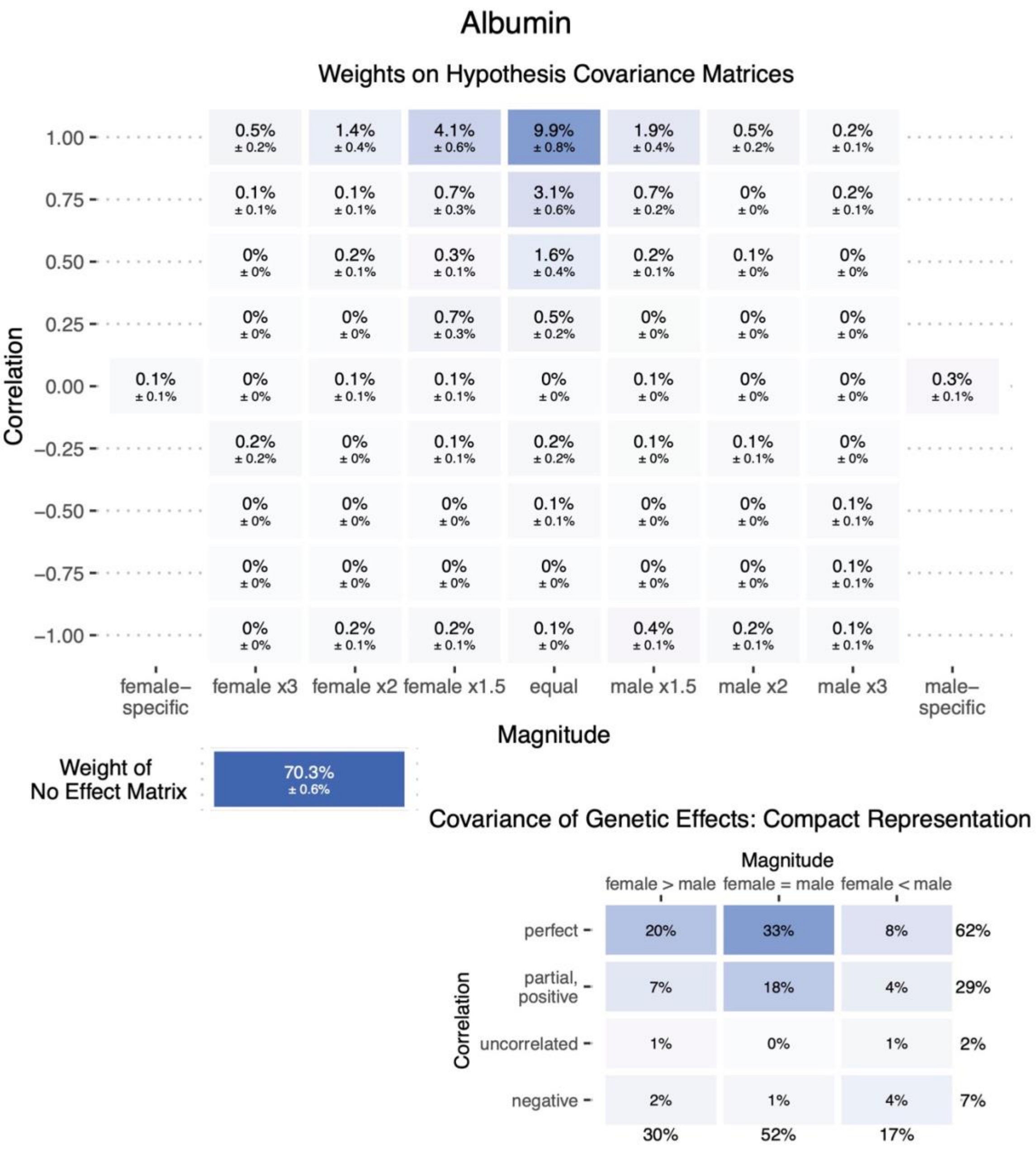

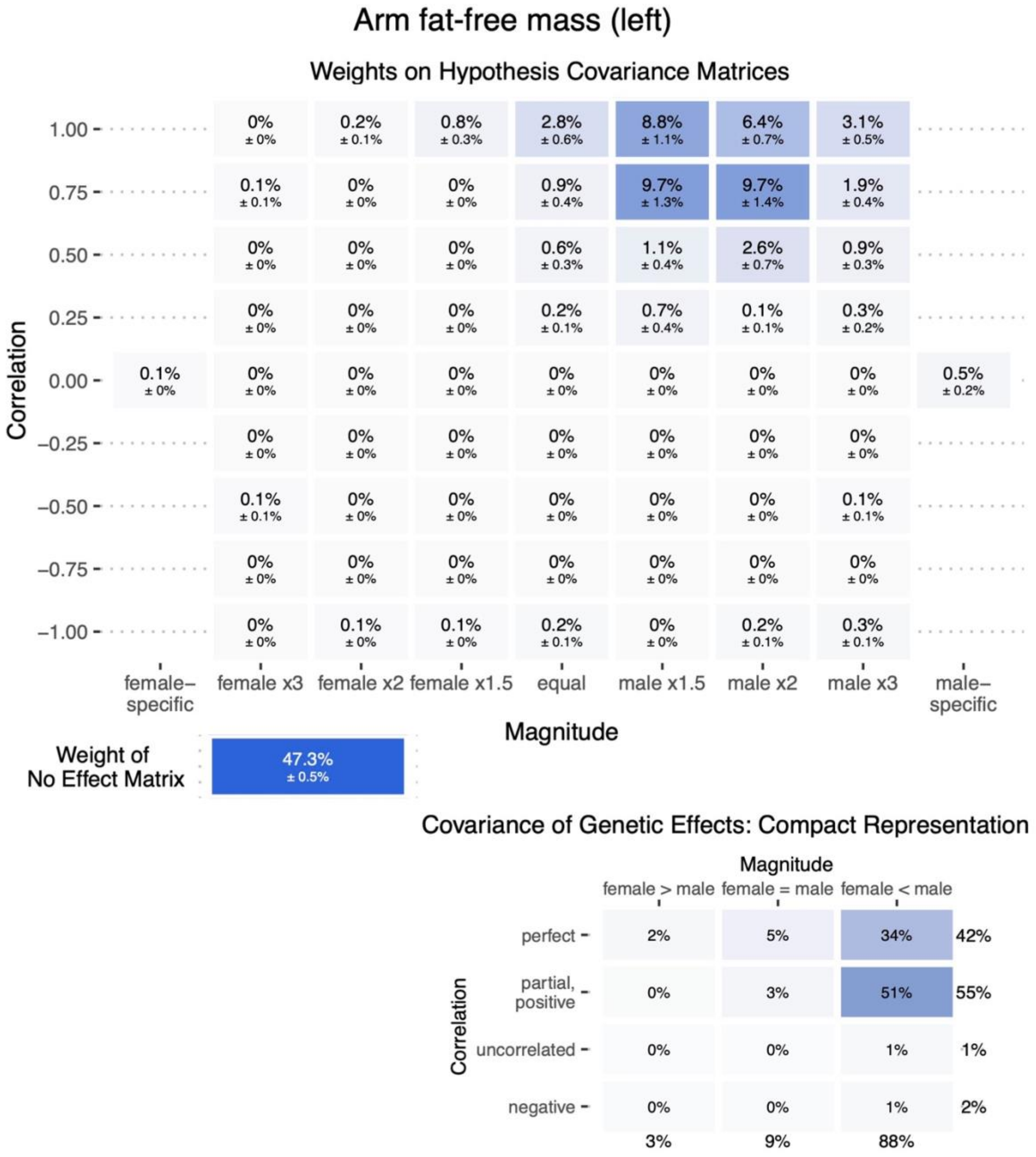
Arm fat-free mass (left)
Weights on Hypothesis Covariance Matrices
Correlation
Magnitude
female–specific
female x3
female x2
female x1.5
equal
male x1.5
male x2
male x3
male–specific
Weight of No Effect Matrix
47.3% ± 0.5%
Covariance of Genetic Effects: Compact Representation
female > male
female = male
female < male
perfect
partial, positive
uncorrelated
negative

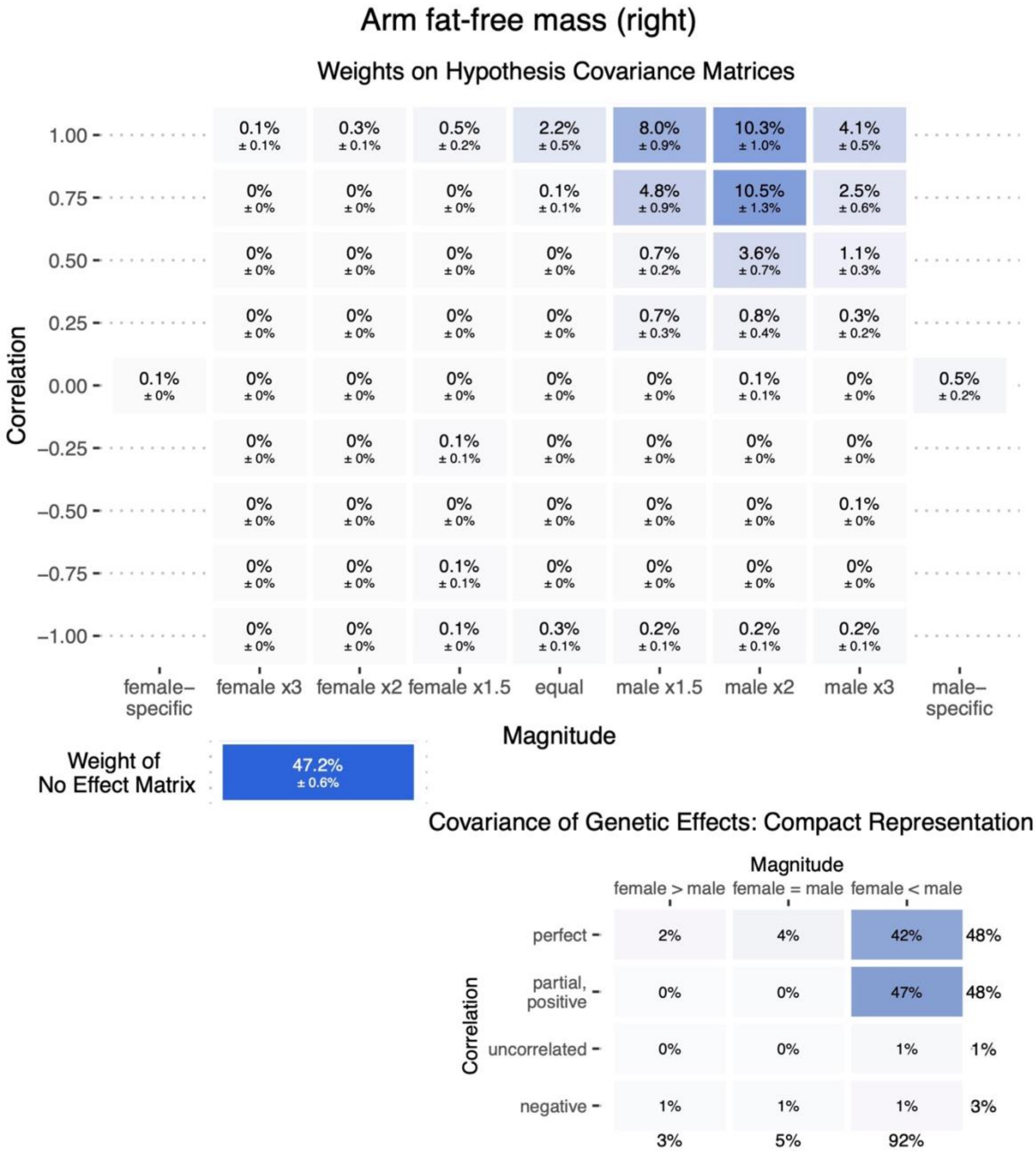
Arm fat-free mass (right)
Weights on Hypothesis Covariance Matrices
Correlation
1.00
0.75
0.50
0.25
0.00
−0.25
−0.50
−0.75
−1.00
1.00: 0.1% ± 0.1% | 0.3% ± 0.1% | 0.5% ± 0.2% | 2.2% ± 0.5% | 8.0% ± 0.9% | 10.3% ± 1.0% | 4.1% ± 0.5%
0.75: 0% ± 0% | 0% ± 0% | 0% ± 0% | 0.1% ± 0.1% | 4.8% ± 0.9% | 10.5% ± 1.3% | 2.5% ± 0.6%
0.50: 0% ± 0% | 0% ± 0% | 0% ± 0% | 0% ± 0% | 0.7% ± 0.2% | 3.6% ± 0.7% | 1.1% ± 0.3%
0.25: 0% ± 0% | 0% ± 0% | 0% ± 0% | 0% ± 0% | 0.7% ± 0.3% | 0.8% ± 0.4% | 0.3% ± 0.2%
0.00: 0.1% ± 0% | 0% ± 0% | 0% ± 0% | 0% ± 0% | 0% ± 0% | 0% ± 0% | 0.1% ± 0.1% | 0% ± 0% | 0.5% ± 0.2%
−0.25: 0% ± 0% | 0% ± 0% | 0.1% ± 0.1% | 0% ± 0% | 0% ± 0% | 0% ± 0% | 0% ± 0%
−0.50: 0% ± 0% | 0% ± 0% | 0% ± 0% | 0% ± 0% | 0% ± 0% | 0% ± 0% | 0.1% ± 0%
−0.75: 0% ± 0% | 0% ± 0% | 0.1% ± 0.1% | 0% ± 0% | 0% ± 0% | 0% ± 0% | 0% ± 0%
−1.00: 0% ± 0% | 0% ± 0% | 0.1% ± 0% | 0.3% ± 0.1% | 0.2% ± 0.1% | 0.2% ± 0.1% | 0.2% ± 0.1%
female–specific
female x3
female x2
female x1.5
equal
male x1.5
male x2
male x3
male–specific
Magnitude
Weight of No Effect Matrix
47.2% ± 0.6%
Covariance of Genetic Effects: Compact Representation
Magnitude
female > male
female = male
female < male
Correlation
perfect: 2% | 4% | 42% | 48%
partial, positive: 0% | 0% | 47% | 48%
uncorrelated: 0% | 0% | 1% | 1%
negative: 1% | 1% | 1% | 3%
3% | 5% | 92%

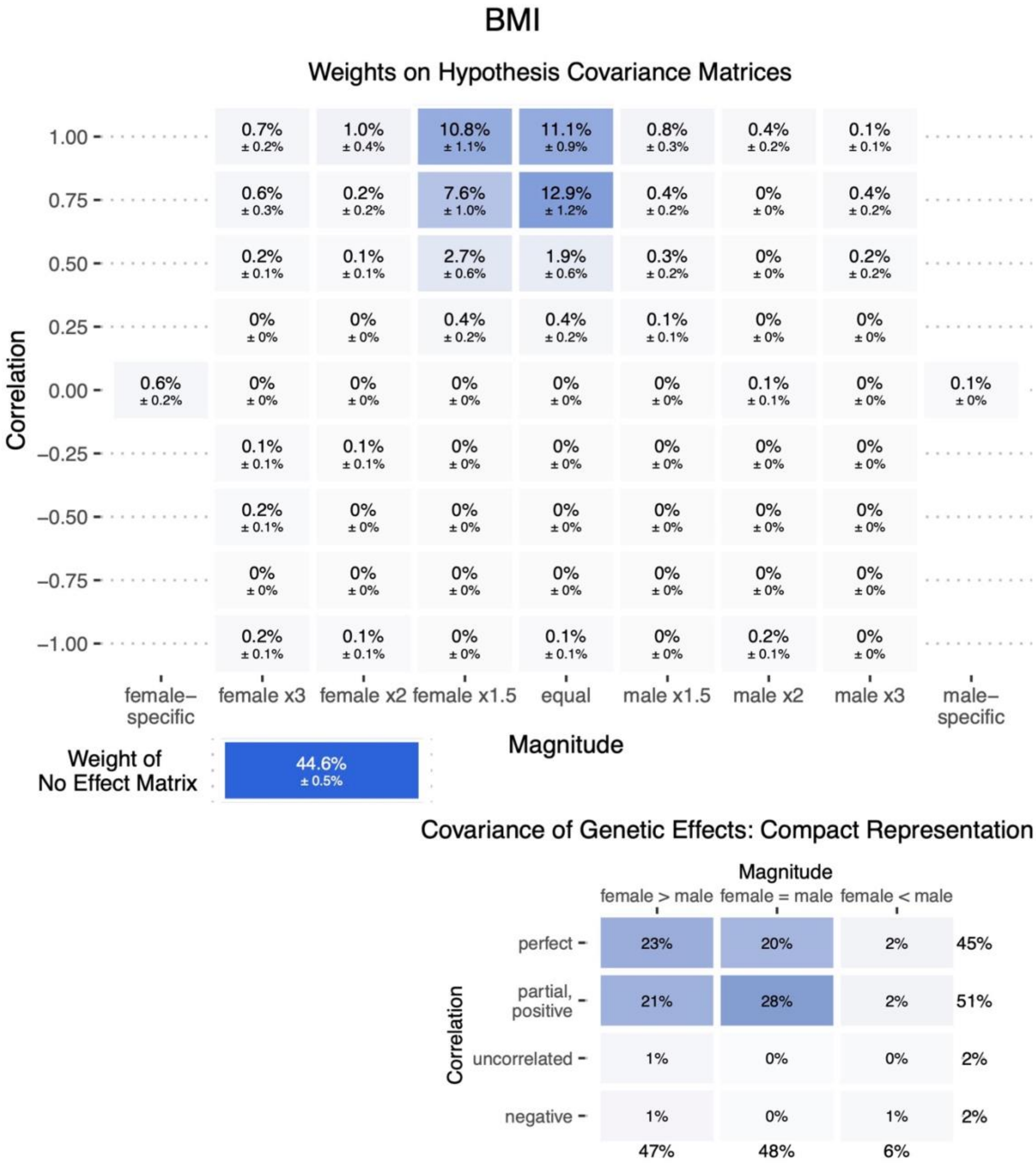
BMI
Weights on Hypothesis Covariance Matrices
Correlation
Magnitude
female–specific
female x3
female x2
female x1.5
equal
male x1.5
male x2
male x3
male–specific
Weight of No Effect Matrix
44.6% ± 0.5%
Covariance of Genetic Effects: Compact Representation
female > male
female = male
female < male
perfect
partial, positive
uncorrelated
negative

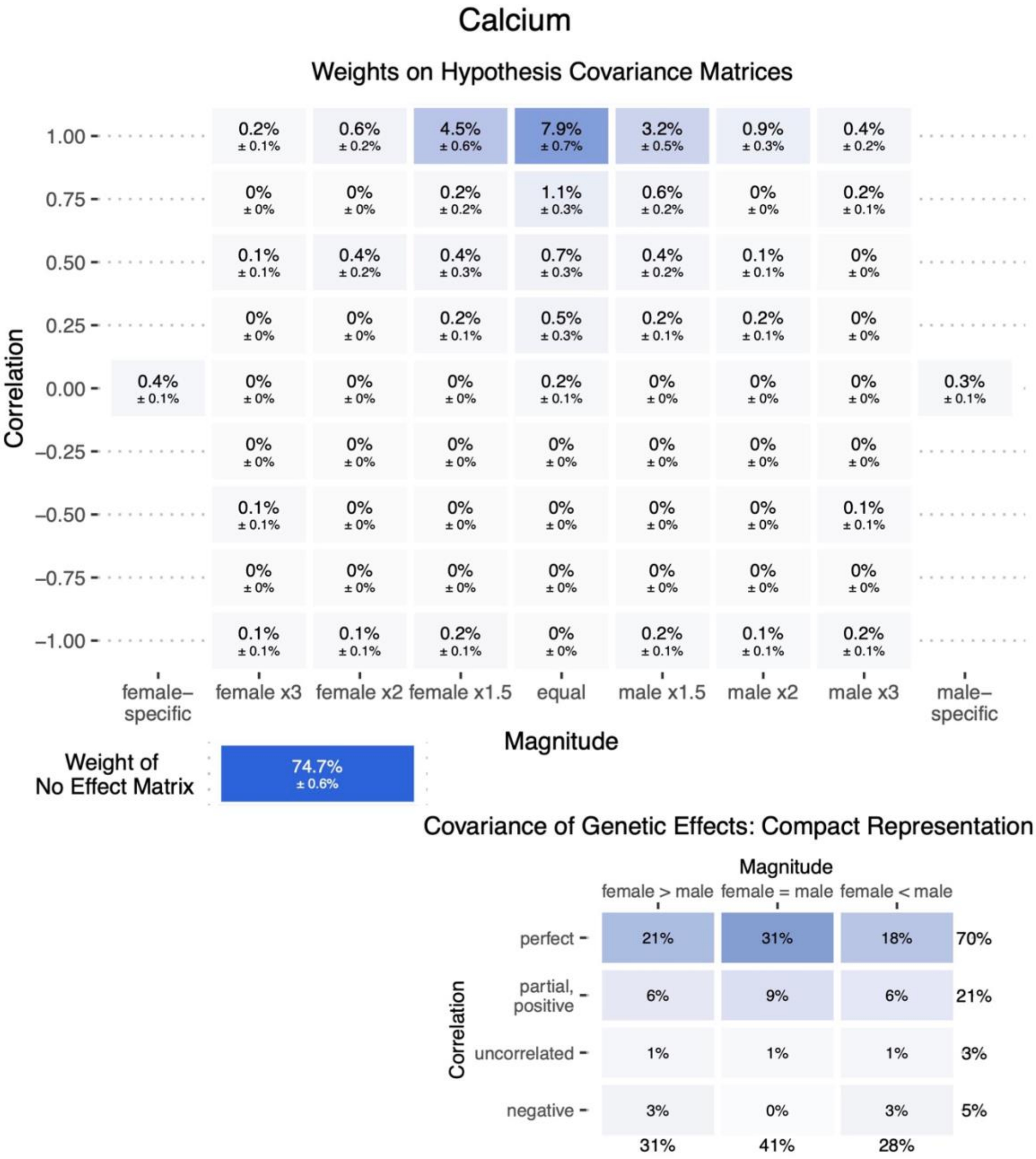
Calcium
Weights on Hypothesis Covariance Matrices
Correlation
1.00
0.75
0.50
0.25
0.00
−0.25
−0.50
−0.75
−1.00
0.2% ± 0.1% | 0.6% ± 0.2% | 4.5% ± 0.6% | 7.9% ± 0.7% | 3.2% ± 0.5% | 0.9% ± 0.3% | 0.4% ± 0.2%
0% ± 0% | 0% ± 0% | 0.2% ± 0.2% | 1.1% ± 0.3% | 0.6% ± 0.2% | 0% ± 0% | 0.2% ± 0.1%
0.1% ± 0.1% | 0.4% ± 0.2% | 0.4% ± 0.3% | 0.7% ± 0.3% | 0.4% ± 0.2% | 0.1% ± 0.1% | 0% ± 0%
0% ± 0% | 0% ± 0% | 0.2% ± 0.1% | 0.5% ± 0.3% | 0.2% ± 0.1% | 0.2% ± 0.1% | 0% ± 0%
0.4% ± 0.1% | 0% ± 0% | 0% ± 0% | 0% ± 0% | 0.2% ± 0.1% | 0% ± 0% | 0% ± 0% | 0% ± 0% | 0.3% ± 0.1%
0% ± 0% | 0% ± 0% | 0% ± 0% | 0% ± 0% | 0% ± 0% | 0% ± 0% | 0% ± 0%
0.1% ± 0.1% | 0% ± 0% | 0% ± 0% | 0% ± 0% | 0% ± 0% | 0% ± 0% | 0.1% ± 0.1%
0% ± 0% | 0% ± 0% | 0% ± 0% | 0% ± 0% | 0% ± 0% | 0% ± 0% | 0% ± 0%
0.1% ± 0.1% | 0.1% ± 0.1% | 0.2% ± 0.1% | 0% ± 0% | 0.2% ± 0.1% | 0.1% ± 0.1% | 0.2% ± 0.1%
female–specific
female x3
female x2
female x1.5
equal
male x1.5
male x2
male x3
male–specific
Magnitude
Weight of No Effect Matrix
74.7% ± 0.6%
Covariance of Genetic Effects: Compact Representation
Magnitude
female > male
female = male
female < male
Correlation
perfect
partial, positive
uncorrelated
negative
21% | 31% | 18% | 70%
6% | 9% | 6% | 21%
1% | 1% | 1% | 3%
3% | 0% | 3% | 5%
31% | 41% | 28%

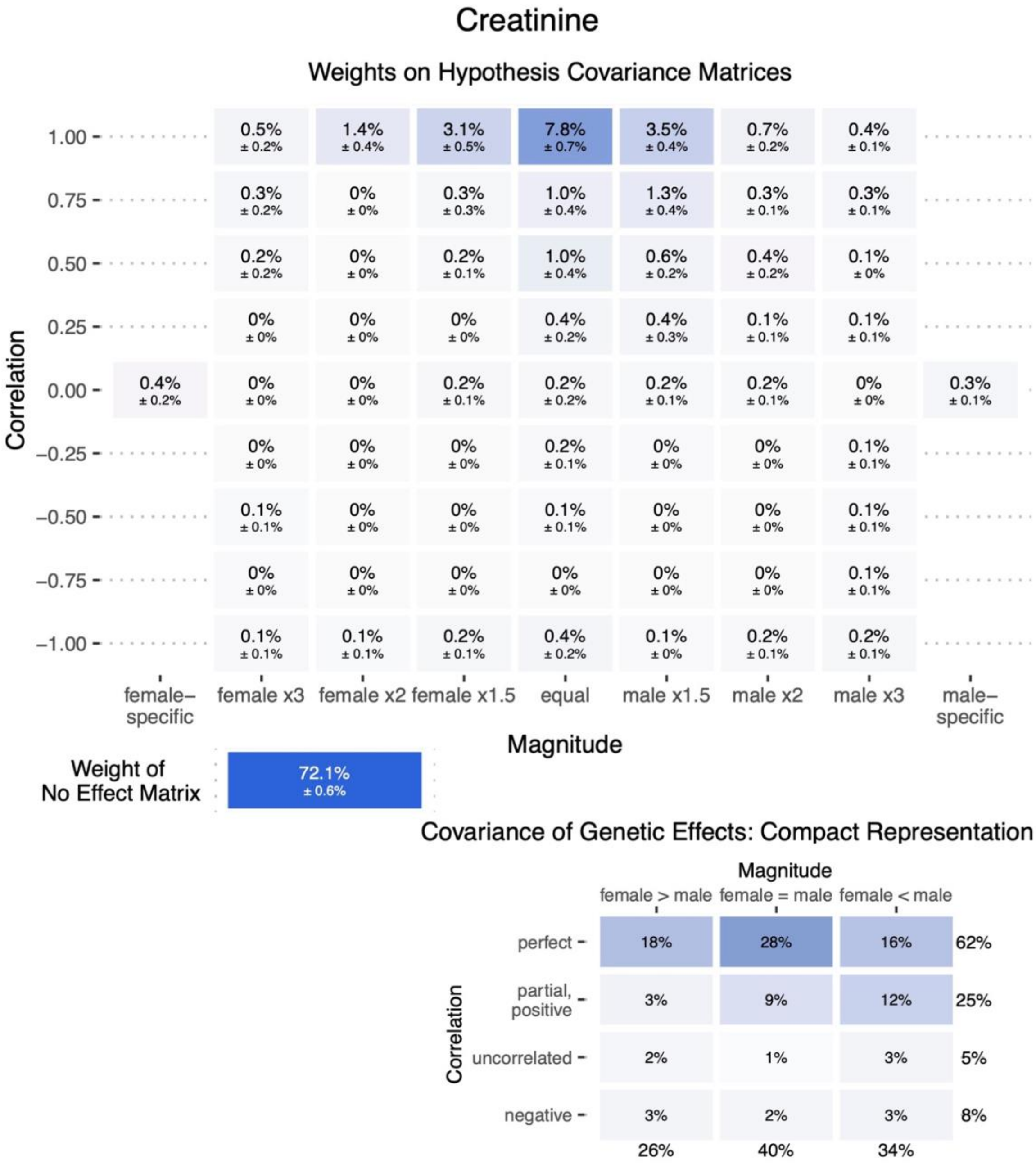
Creatinine
Weights on Hypothesis Covariance Matrices
Correlation
1.00
0.75
0.50
0.25
0.00
−0.25
−0.50
−0.75
−1.00
1.00: 0.5% ± 0.2% | 1.4% ± 0.4% | 3.1% ± 0.5% | 7.8% ± 0.7% | 3.5% ± 0.4% | 0.7% ± 0.2% | 0.4% ± 0.1%
0.75: 0.3% ± 0.2% | 0% ± 0% | 0.3% ± 0.3% | 1.0% ± 0.4% | 1.3% ± 0.4% | 0.3% ± 0.1% | 0.3% ± 0.1%
0.50: 0.2% ± 0.2% | 0% ± 0% | 0.2% ± 0.1% | 1.0% ± 0.4% | 0.6% ± 0.2% | 0.4% ± 0.2% | 0.1% ± 0%
0.25: 0% ± 0% | 0% ± 0% | 0% ± 0% | 0.4% ± 0.2% | 0.4% ± 0.3% | 0.1% ± 0.1% | 0.1% ± 0.1%
0.00: 0.4% ± 0.2% | 0% ± 0% | 0% ± 0% | 0.2% ± 0.1% | 0.2% ± 0.2% | 0.2% ± 0.1% | 0.2% ± 0.1% | 0% ± 0% | 0.3% ± 0.1%
−0.25: 0% ± 0% | 0% ± 0% | 0% ± 0% | 0.2% ± 0.1% | 0% ± 0% | 0% ± 0% | 0.1% ± 0.1%
−0.50: 0.1% ± 0.1% | 0% ± 0% | 0% ± 0% | 0.1% ± 0.1% | 0% ± 0% | 0% ± 0% | 0.1% ± 0.1%
−0.75: 0% ± 0% | 0% ± 0% | 0% ± 0% | 0% ± 0% | 0% ± 0% | 0% ± 0% | 0.1% ± 0.1%
−1.00: 0.1% ± 0.1% | 0.1% ± 0.1% | 0.2% ± 0.1% | 0.4% ± 0.2% | 0.1% ± 0% | 0.2% ± 0.1% | 0.2% ± 0.1%
female–specific
female x3
female x2
female x1.5
equal
male x1.5
male x2
male x3
male–specific
Magnitude
Weight of No Effect Matrix
72.1% ± 0.6%
Covariance of Genetic Effects: Compact Representation
Magnitude
female > male
female = male
female < male
Correlation
perfect: 18% | 28% | 16% | 62%
partial, positive: 3% | 9% | 12% | 25%
uncorrelated: 2% | 1% | 3% | 5%
negative: 3% | 2% | 3% | 8%
26% | 40% | 34%

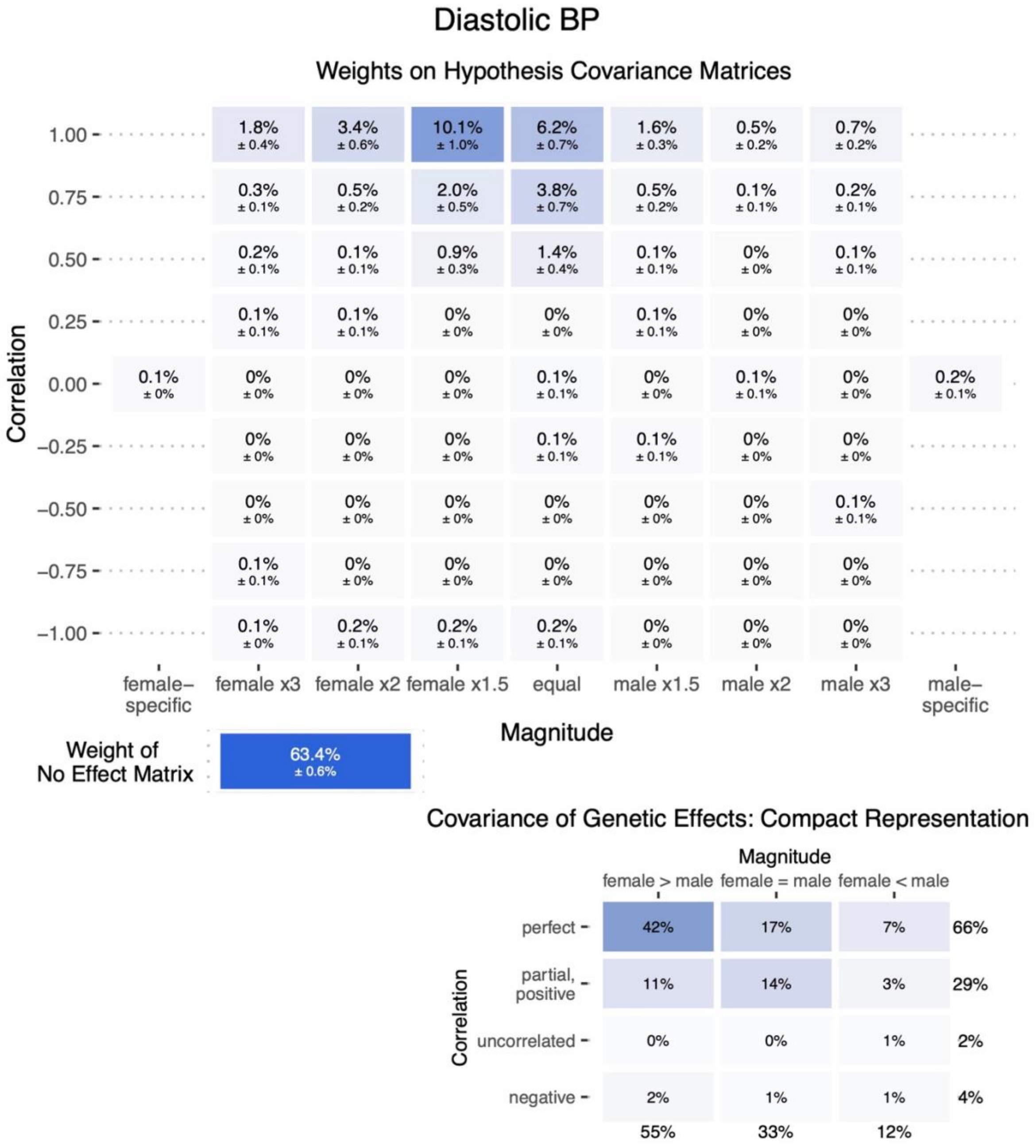
Diastolic BP
Weights on Hypothesis Covariance Matrices
Correlation
Magnitude
female–specific
female x3
female x2
female x1.5
equal
male x1.5
male x2
male x3
male–specific
Weight of No Effect Matrix
63.4% ± 0.6%
Covariance of Genetic Effects: Compact Representation
Magnitude
female > male
female = male
female < male
perfect
partial, positive
uncorrelated
negative
42%
17%
7%
66%
11%
14%
3%
29%
0%
0%
1%
2%
2%
1%
1%
4%
55%
33%
12%

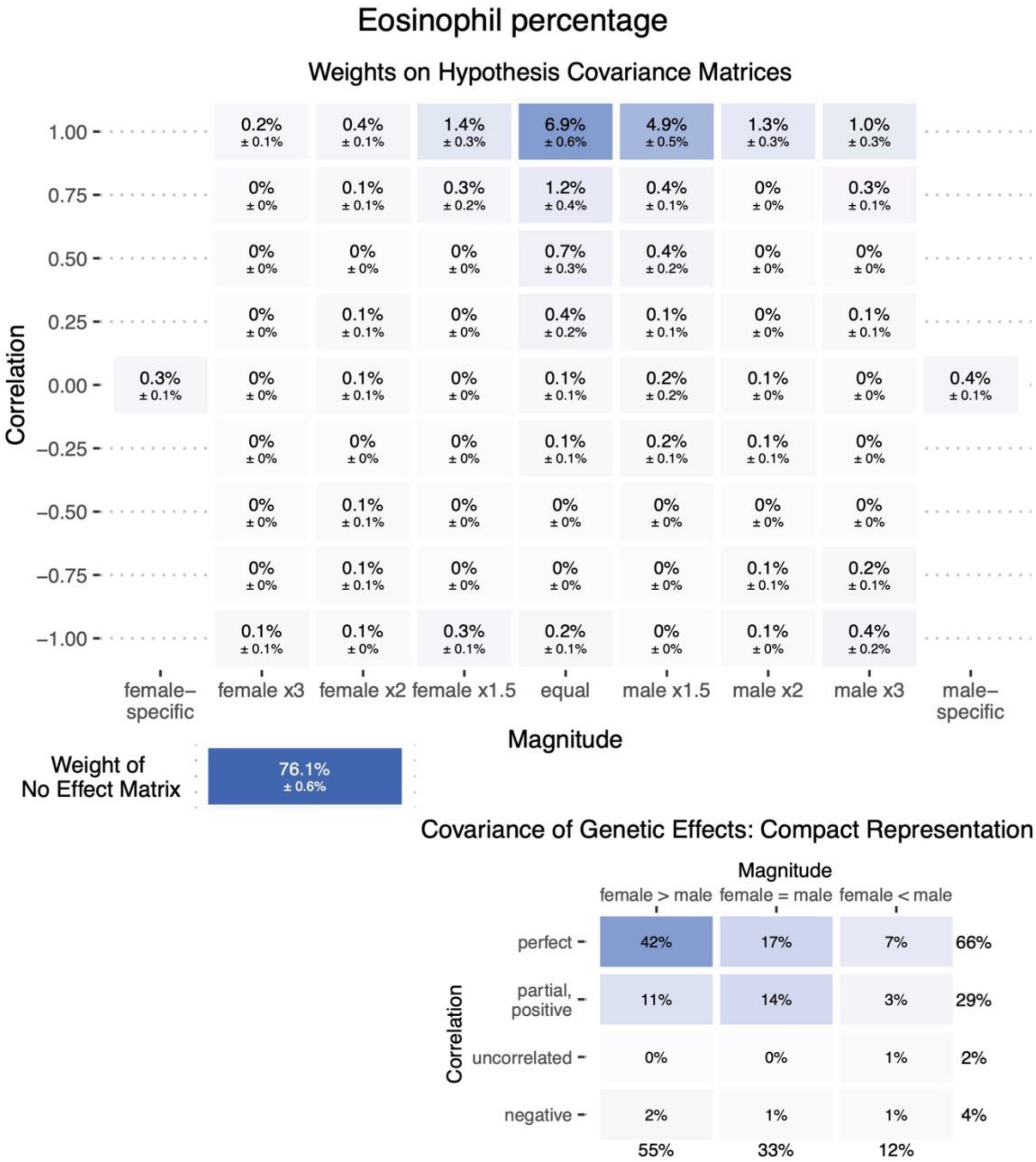
Eosinophil percentage
Weights on Hypothesis Covariance Matrices
Correlation
1.00
0.75
0.50
0.25
0.00
−0.25
−0.50
−0.75
−1.00
0.2% ± 0.1%
0.4% ± 0.1%
1.4% ± 0.3%
6.9% ± 0.6%
4.9% ± 0.5%
1.3% ± 0.3%
1.0% ± 0.3%
0% ± 0%
0.1% ± 0.1%
0.3% ± 0.2%
1.2% ± 0.4%
0.4% ± 0.1%
0% ± 0%
0.3% ± 0.1%
0% ± 0%
0% ± 0%
0% ± 0%
0.7% ± 0.3%
0.4% ± 0.2%
0% ± 0%
0% ± 0%
0% ± 0%
0.1% ± 0.1%
0% ± 0%
0.4% ± 0.2%
0.1% ± 0.1%
0% ± 0%
0.1% ± 0.1%
0.3% ± 0.1%
0% ± 0%
0.1% ± 0.1%
0% ± 0%
0.1% ± 0.1%
0.2% ± 0.2%
0.1% ± 0%
0% ± 0%
0.4% ± 0.1%
0% ± 0%
0% ± 0%
0% ± 0%
0.1% ± 0.1%
0.2% ± 0.1%
0.1% ± 0.1%
0% ± 0%
0% ± 0%
0.1% ± 0.1%
0% ± 0%
0% ± 0%
0% ± 0%
0% ± 0%
0% ± 0%
0% ± 0%
0.1% ± 0.1%
0% ± 0%
0% ± 0%
0% ± 0%
0.1% ± 0.1%
0.2% ± 0.1%
0.1% ± 0.1%
0.1% ± 0%
0.3% ± 0.1%
0.2% ± 0.1%
0% ± 0%
0.1% ± 0%
0.4% ± 0.2%
female–specific
female x3
female x2
female x1.5
equal
male x1.5
male x2
male x3
male–specific
Magnitude
Weight of No Effect Matrix
76.1% ± 0.6%
Covariance of Genetic Effects: Compact Representation
Magnitude
female > male
female = male
female < male
Correlation
perfect
partial, positive
uncorrelated
negative
42%
17%
7%
66%
11%
14%
3%
29%
0%
0%
1%
2%
2%
1%
1%
4%
55%
33%
12%

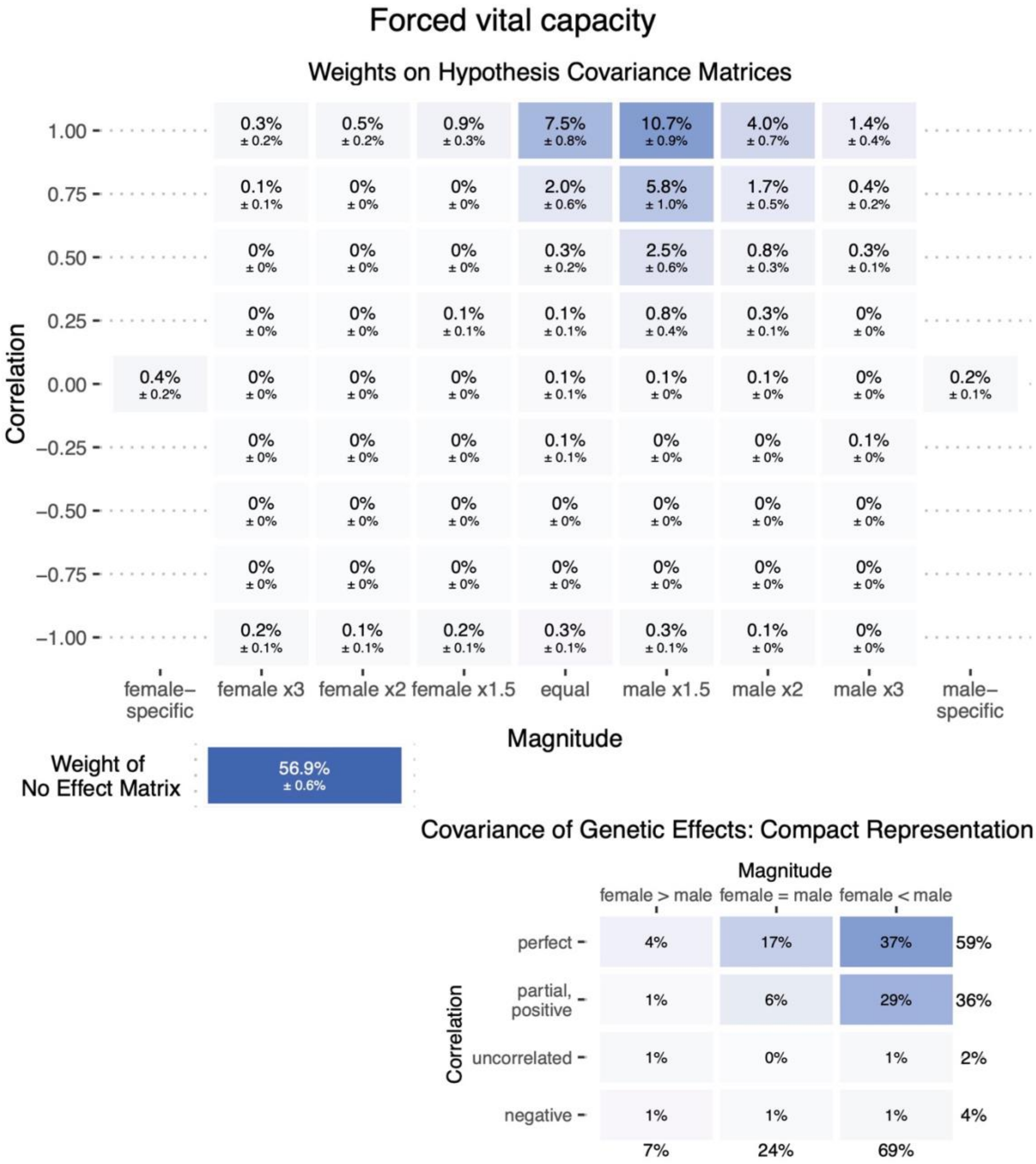

Forced vital capacity
Weights on Hypothesis Covariance Matrices
Correlation
1.00
0.75
0.50
0.25
0.00
−0.25
−0.50
−0.75
−1.00
0.3% ± 0.2%
0.5% ± 0.2%
0.9% ± 0.3%
7.5% ± 0.8%
10.7% ± 0.9%
4.0% ± 0.7%
1.4% ± 0.4%
0.1% ± 0.1%
2.0% ± 0.6%
5.8% ± 1.0%
1.7% ± 0.5%
0.4% ± 0.2%
0.3% ± 0.2%
2.5% ± 0.6%
0.8% ± 0.3%
0.3% ± 0.1%
0.8% ± 0.4%
0.4% ± 0.2%
0.2% ± 0.1%
56.9% ± 0.6%
female−specific
female x3
female x2
female x1.5
equal
male x1.5
male x2
male x3
male−specific
Magnitude
Weight of No Effect Matrix
Covariance of Genetic Effects: Compact Representation
Magnitude
female > male
female = male
female < male
Correlation
perfect
partial, positive
uncorrelated
negative
4%
17%
37%
59%
1%
6%
29%
36%
1%
0%
1%
2%
1%
1%
1%
4%
7%
24%
69%

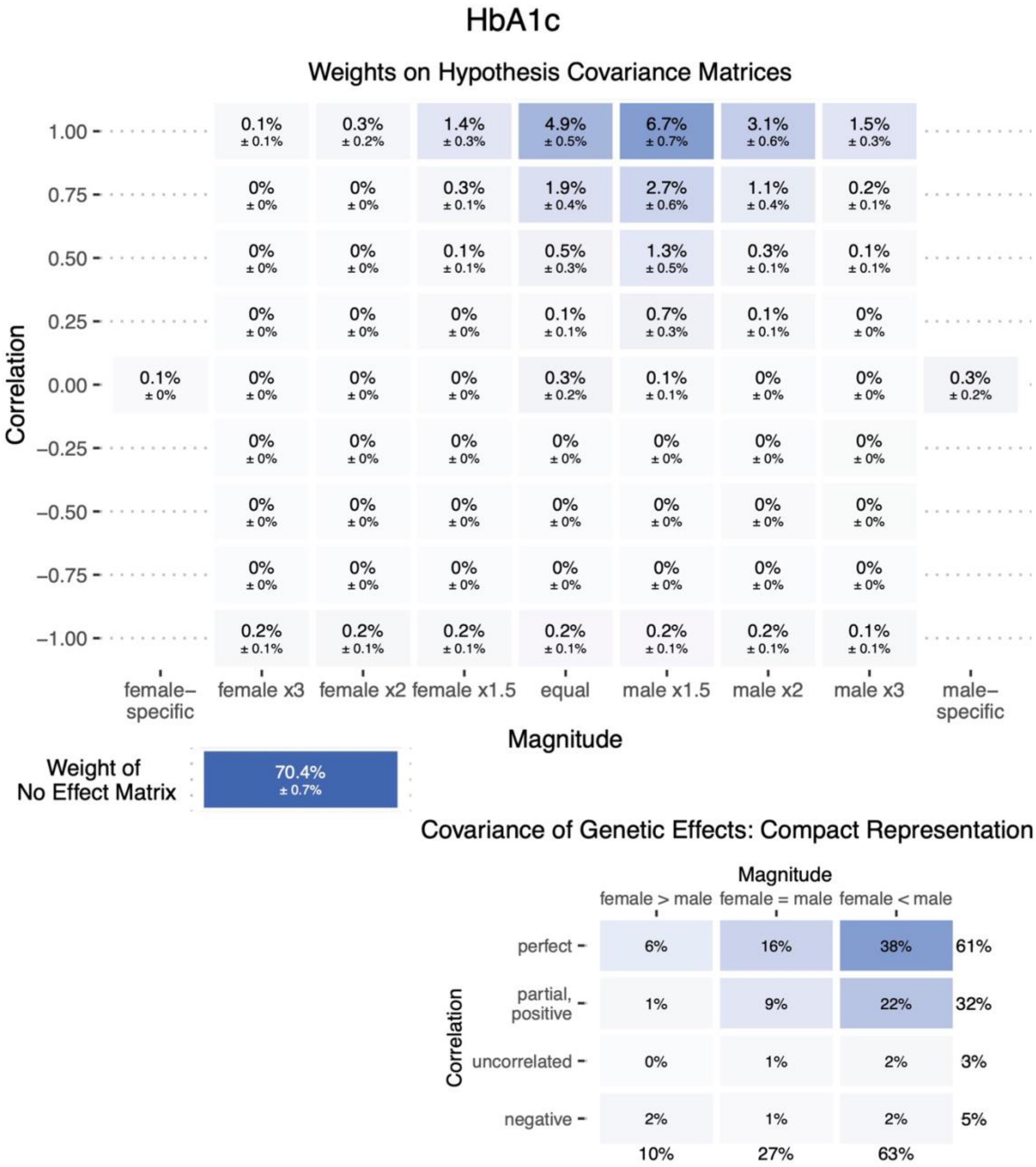
HbA1c
Weights on Hypothesis Covariance Matrices
Correlation
1.00
0.75
0.50
0.25
0.00
−0.25
−0.50
−0.75
−1.00
0.1% ± 0.1%
0.3% ± 0.2%
1.4% ± 0.3%
4.9% ± 0.5%
6.7% ± 0.7%
3.1% ± 0.6%
1.5% ± 0.3%
0% ± 0%
0% ± 0%
0.3% ± 0.1%
1.9% ± 0.4%
2.7% ± 0.6%
1.1% ± 0.4%
0.2% ± 0.1%
0% ± 0%
0% ± 0%
0.1% ± 0.1%
0.5% ± 0.3%
1.3% ± 0.5%
0.3% ± 0.1%
0.1% ± 0.1%
0% ± 0%
0% ± 0%
0% ± 0%
0.1% ± 0.1%
0.7% ± 0.3%
0.1% ± 0.1%
0% ± 0%
0.1% ± 0%
0% ± 0%
0% ± 0%
0% ± 0%
0.3% ± 0.2%
0.1% ± 0.1%
0% ± 0%
0% ± 0%
0.3% ± 0.2%
0% ± 0%
0.2% ± 0.1%
0.2% ± 0.1%
0.2% ± 0.1%
0.2% ± 0.1%
0.2% ± 0.1%
0.2% ± 0.1%
0.1% ± 0.1%
female− specific
female x3
female x2
female x1.5
equal
male x1.5
male x2
male x3
male− specific
Magnitude
Weight of No Effect Matrix
70.4% ± 0.7%
Covariance of Genetic Effects: Compact Representation
Magnitude
female > male
female = male
female < male
Correlation
perfect
partial, positive
uncorrelated
negative
6%
16%
38%
61%
1%
9%
22%
32%
0%
1%
2%
3%
2%
1%
2%
5%
10%
27%
63%

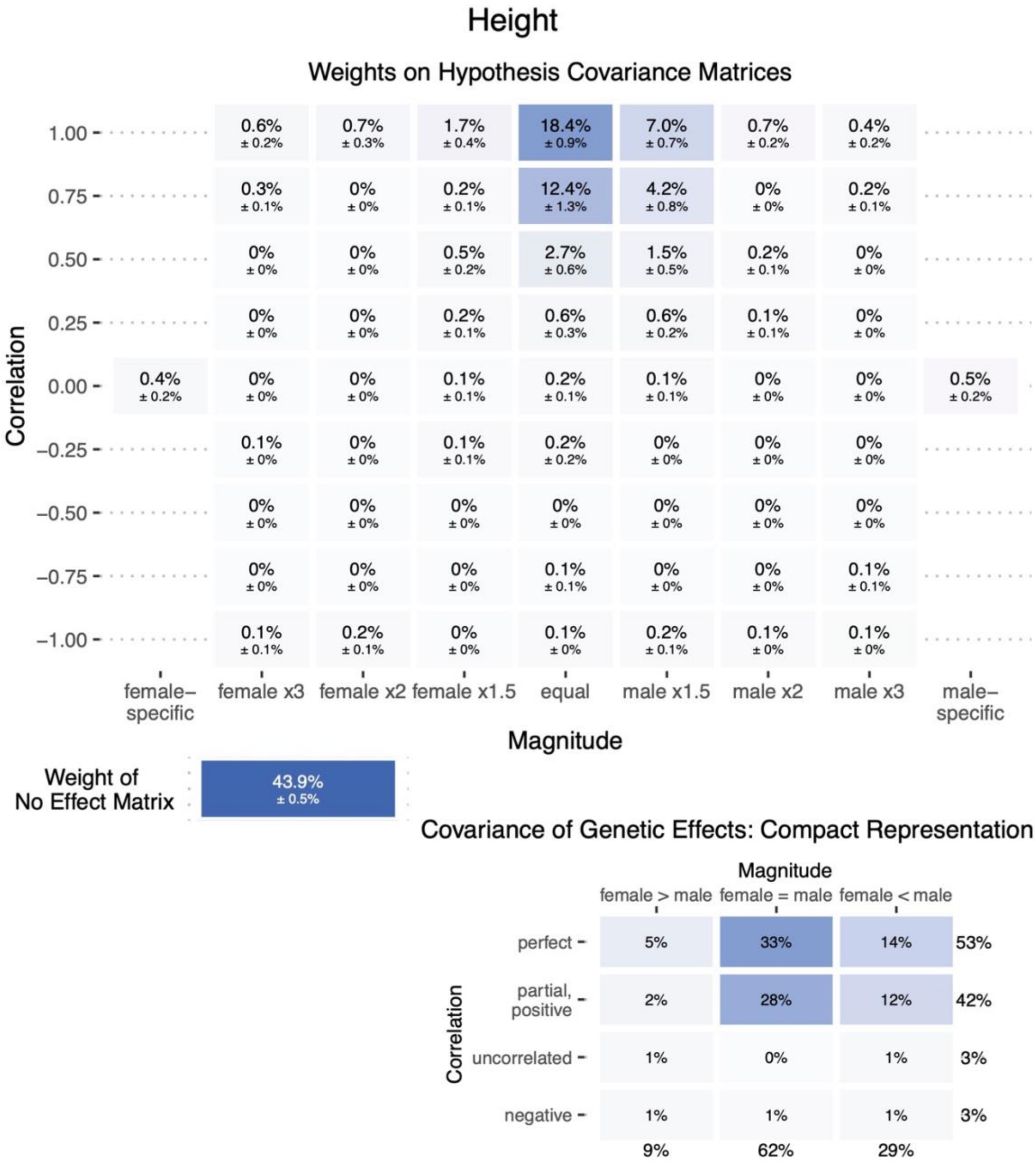
Height
Weights on Hypothesis Covariance Matrices
Correlation
1.00
0.75
0.50
0.25
0.00
−0.25
−0.50
−0.75
−1.00
0.6% ± 0.2%
0.7% ± 0.3%
1.7% ± 0.4%
18.4% ± 0.9%
7.0% ± 0.7%
0.7% ± 0.2%
0.4% ± 0.2%
0.3% ± 0.1%
0% ± 0%
0.2% ± 0.1%
12.4% ± 1.3%
4.2% ± 0.8%
0% ± 0%
0.2% ± 0.1%
0% ± 0%
0% ± 0%
0.5% ± 0.2%
2.7% ± 0.6%
1.5% ± 0.5%
0.2% ± 0.1%
0% ± 0%
0% ± 0%
0% ± 0%
0.2% ± 0.1%
0.6% ± 0.3%
0.6% ± 0.2%
0.1% ± 0.1%
0% ± 0%
0.4% ± 0.2%
0% ± 0%
0% ± 0%
0.1% ± 0.1%
0.2% ± 0.1%
0.1% ± 0.1%
0% ± 0%
0% ± 0%
0.5% ± 0.2%
0.1% ± 0%
0% ± 0%
0.1% ± 0.1%
0.2% ± 0.2%
0% ± 0%
0% ± 0%
0% ± 0%
0% ± 0%
0% ± 0%
0% ± 0%
0% ± 0%
0% ± 0%
0% ± 0%
0% ± 0%
0% ± 0%
0% ± 0%
0% ± 0%
0.1% ± 0.1%
0% ± 0%
0% ± 0%
0.1% ± 0.1%
0.1% ± 0.1%
0.2% ± 0.1%
0% ± 0%
0.1% ± 0%
0.2% ± 0.1%
0.1% ± 0%
0.1% ± 0%
female–specific
female x3
female x2
female x1.5
equal
male x1.5
male x2
male x3
male–specific
Magnitude
Weight of No Effect Matrix
43.9% ± 0.5%
Covariance of Genetic Effects: Compact Representation
Magnitude
female > male
female = male
female < male
Correlation
perfect
partial, positive
uncorrelated
negative
5%
33%
14%
53%
2%
28%
12%
42%
1%
0%
1%
3%
1%
1%
1%
3%
9%
62%
29%

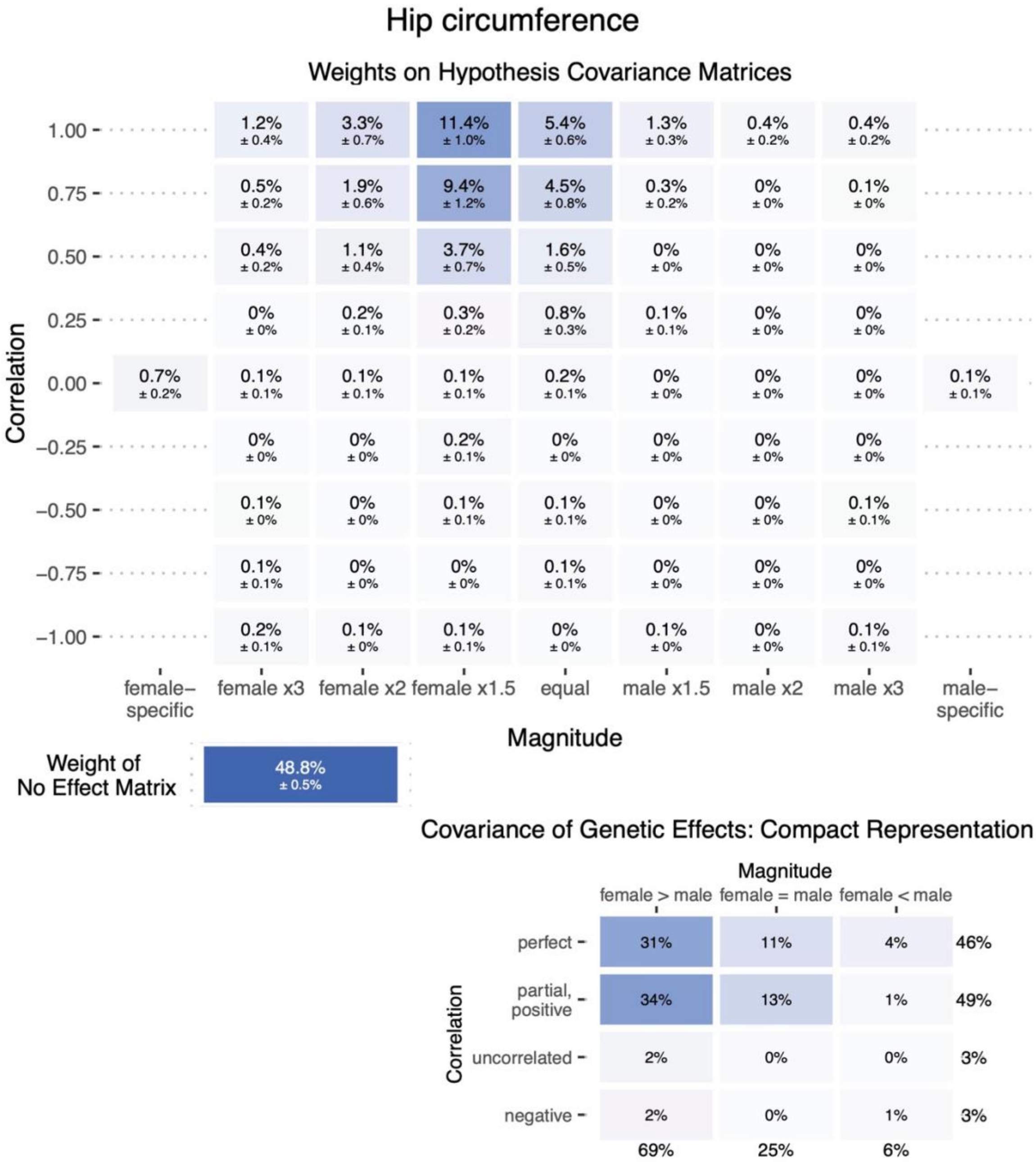

Hip circumference
Weights on Hypothesis Covariance Matrices
Correlation
Magnitude
female–specific
female x3
female x2
female x1.5
equal
male x1.5
male x2
male x3
male–specific
Weight of No Effect Matrix
48.8% ± 0.5%
Covariance of Genetic Effects: Compact Representation
female > male
female = male
female < male
perfect
partial, positive
uncorrelated
negative

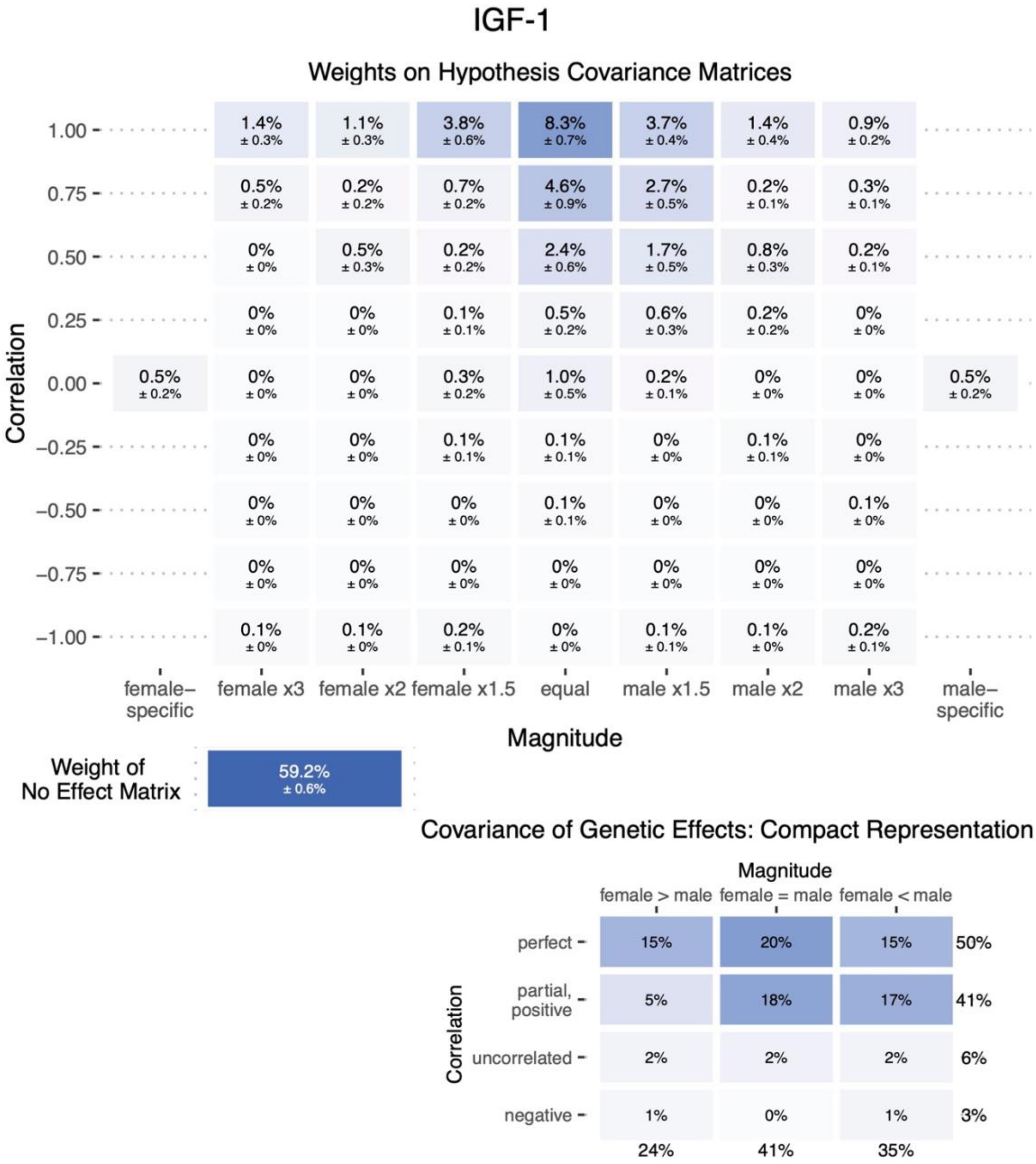
IGF-1
Weights on Hypothesis Covariance Matrices
Correlation
1.00 - 1.4% ± 0.3% | 1.1% ± 0.3% | 3.8% ± 0.6% | 8.3% ± 0.7% | 3.7% ± 0.4% | 1.4% ± 0.4% | 0.9% ± 0.2%
0.75 - 0.5% ± 0.2% | 0.2% ± 0.2% | 0.7% ± 0.2% | 4.6% ± 0.9% | 2.7% ± 0.5% | 0.2% ± 0.1% | 0.3% ± 0.1%
0.50 - 0% ± 0% | 0.5% ± 0.3% | 0.2% ± 0.2% | 2.4% ± 0.6% | 1.7% ± 0.5% | 0.8% ± 0.3% | 0.2% ± 0.1%
0.25 - 0% ± 0% | 0% ± 0% | 0.1% ± 0.1% | 0.5% ± 0.2% | 0.6% ± 0.3% | 0.2% ± 0.2% | 0% ± 0%
0.00 - 0.5% ± 0.2% | 0% ± 0% | 0% ± 0% | 0.3% ± 0.2% | 1.0% ± 0.5% | 0.2% ± 0.1% | 0% ± 0% | 0% ± 0% | 0.5% ± 0.2%
−0.25 - 0% ± 0% | 0% ± 0% | 0.1% ± 0.1% | 0.1% ± 0.1% | 0% ± 0% | 0.1% ± 0.1% | 0% ± 0%
−0.50 - 0% ± 0% | 0% ± 0% | 0% ± 0% | 0.1% ± 0.1% | 0% ± 0% | 0% ± 0% | 0.1% ± 0%
−0.75 - 0% ± 0% | 0% ± 0% | 0% ± 0% | 0% ± 0% | 0% ± 0% | 0% ± 0% | 0% ± 0%
−1.00 - 0.1% ± 0% | 0.1% ± 0% | 0.2% ± 0.1% | 0% ± 0% | 0.1% ± 0.1% | 0.1% ± 0% | 0.2% ± 0.1%
female– specific
female x3
female x2
female x1.5
equal
male x1.5
male x2
male x3
male– specific
Magnitude
Weight of No Effect Matrix
59.2% ± 0.6%
Covariance of Genetic Effects: Compact Representation
Magnitude
female > male
female = male
female < male
Correlation
perfect - 15% | 20% | 15% | 50%
partial, positive - 5% | 18% | 17% | 41%
uncorrelated - 2% | 2% | 2% | 6%
negative - 1% | 0% | 1% | 3%
24% | 41% | 35%

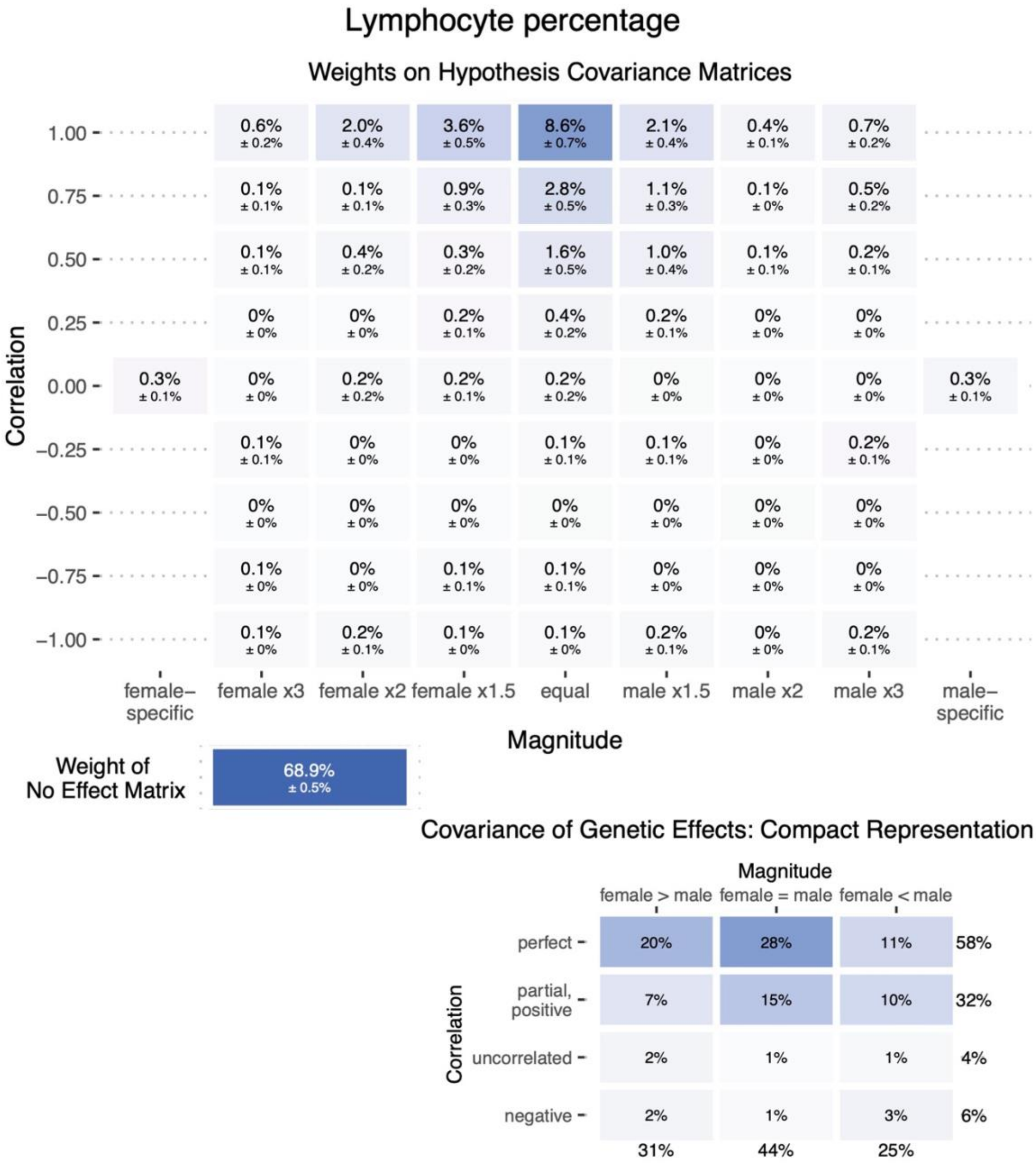
Lymphocyte percentage
Weights on Hypothesis Covariance Matrices
Correlation
1.00
0.75
0.50
0.25
0.00
−0.25
−0.50
−0.75
−1.00
0.6% ± 0.2% | 2.0% ± 0.4% | 3.6% ± 0.5% | 8.6% ± 0.7% | 2.1% ± 0.4% | 0.4% ± 0.1% | 0.7% ± 0.2%
0.1% ± 0.1% | 0.1% ± 0.1% | 0.9% ± 0.3% | 2.8% ± 0.5% | 1.1% ± 0.3% | 0.1% ± 0% | 0.5% ± 0.2%
0.1% ± 0.1% | 0.4% ± 0.2% | 0.3% ± 0.2% | 1.6% ± 0.5% | 1.0% ± 0.4% | 0.1% ± 0.1% | 0.2% ± 0.1%
0% ± 0% | 0% ± 0% | 0.2% ± 0.1% | 0.4% ± 0.2% | 0.2% ± 0.1% | 0% ± 0% | 0% ± 0%
0.3% ± 0.1% | 0% ± 0% | 0.2% ± 0.2% | 0.2% ± 0.1% | 0.2% ± 0.2% | 0% ± 0% | 0% ± 0% | 0% ± 0% | 0.3% ± 0.1%
0.1% ± 0.1% | 0% ± 0% | 0% ± 0% | 0.1% ± 0.1% | 0.1% ± 0.1% | 0% ± 0% | 0.2% ± 0.1%
0% ± 0% | 0% ± 0% | 0% ± 0% | 0% ± 0% | 0% ± 0% | 0% ± 0% | 0% ± 0%
0.1% ± 0% | 0% ± 0% | 0.1% ± 0.1% | 0.1% ± 0.1% | 0% ± 0% | 0% ± 0% | 0% ± 0%
0.1% ± 0% | 0.2% ± 0.1% | 0.1% ± 0% | 0.1% ± 0% | 0.2% ± 0.1% | 0% ± 0% | 0.2% ± 0.1%
female–specific
female x3
female x2
female x1.5
equal
male x1.5
male x2
male x3
male–specific
Magnitude
Weight of No Effect Matrix
68.9% ± 0.5%
Covariance of Genetic Effects: Compact Representation
Magnitude
female > male
female = male
female < male
Correlation
perfect
partial, positive
uncorrelated
negative
20% | 28% | 11% | 58%
7% | 15% | 10% | 32%
2% | 1% | 1% | 4%
2% | 1% | 3% | 6%
31% | 44% | 25%

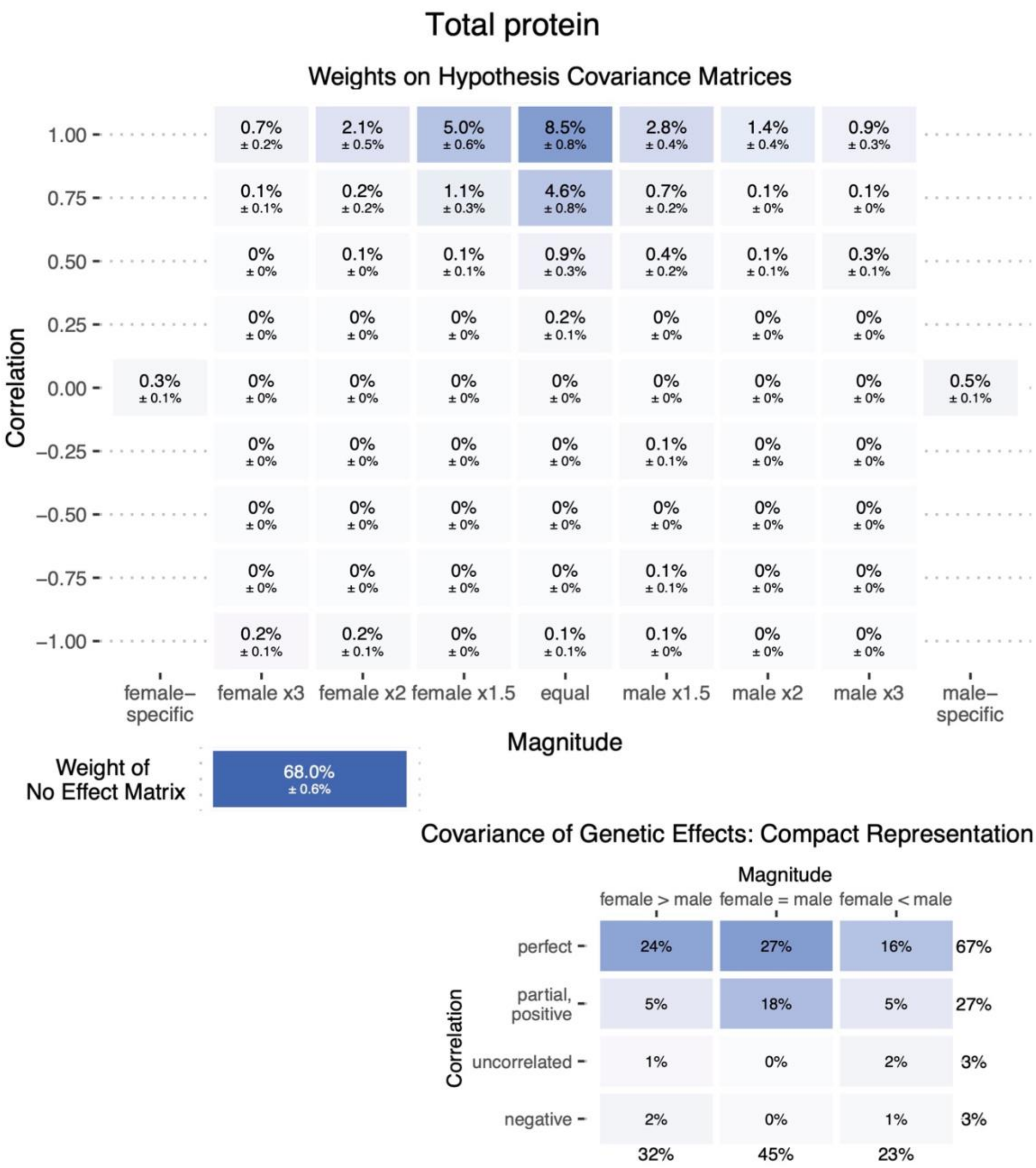
Total protein
Weights on Hypothesis Covariance Matrices
Correlation
1.00
0.75
0.50
0.25
0.00
−0.25
−0.50
−0.75
−1.00
0.7% ± 0.2%
2.1% ± 0.5%
5.0% ± 0.6%
8.5% ± 0.8%
2.8% ± 0.4%
1.4% ± 0.4%
0.9% ± 0.3%
0.1% ± 0.1%
0.2% ± 0.2%
1.1% ± 0.3%
4.6% ± 0.8%
0.7% ± 0.2%
0.1% ± 0%
0.1% ± 0%
0% ± 0%
0.1% ± 0%
0.1% ± 0.1%
0.9% ± 0.3%
0.4% ± 0.2%
0.1% ± 0.1%
0.3% ± 0.1%
0.2% ± 0.1%
0.3% ± 0.1%
0.5% ± 0.1%
0.1% ± 0.1%
0.1% ± 0.1%
0.2% ± 0.1%
0.2% ± 0.1%
0% ± 0%
0.1% ± 0.1%
0.1% ± 0%
female–specific
female x3
female x2
female x1.5
equal
male x1.5
male x2
male x3
male–specific
Magnitude
Weight of No Effect Matrix
68.0% ± 0.6%
Covariance of Genetic Effects: Compact Representation
Magnitude
female > male
female = male
female < male
Correlation
perfect
partial, positive
uncorrelated
negative
24%
27%
16%
67%
5%
18%
5%
27%
1%
0%
2%
3%
2%
0%
1%
3%
32%
45%
23%

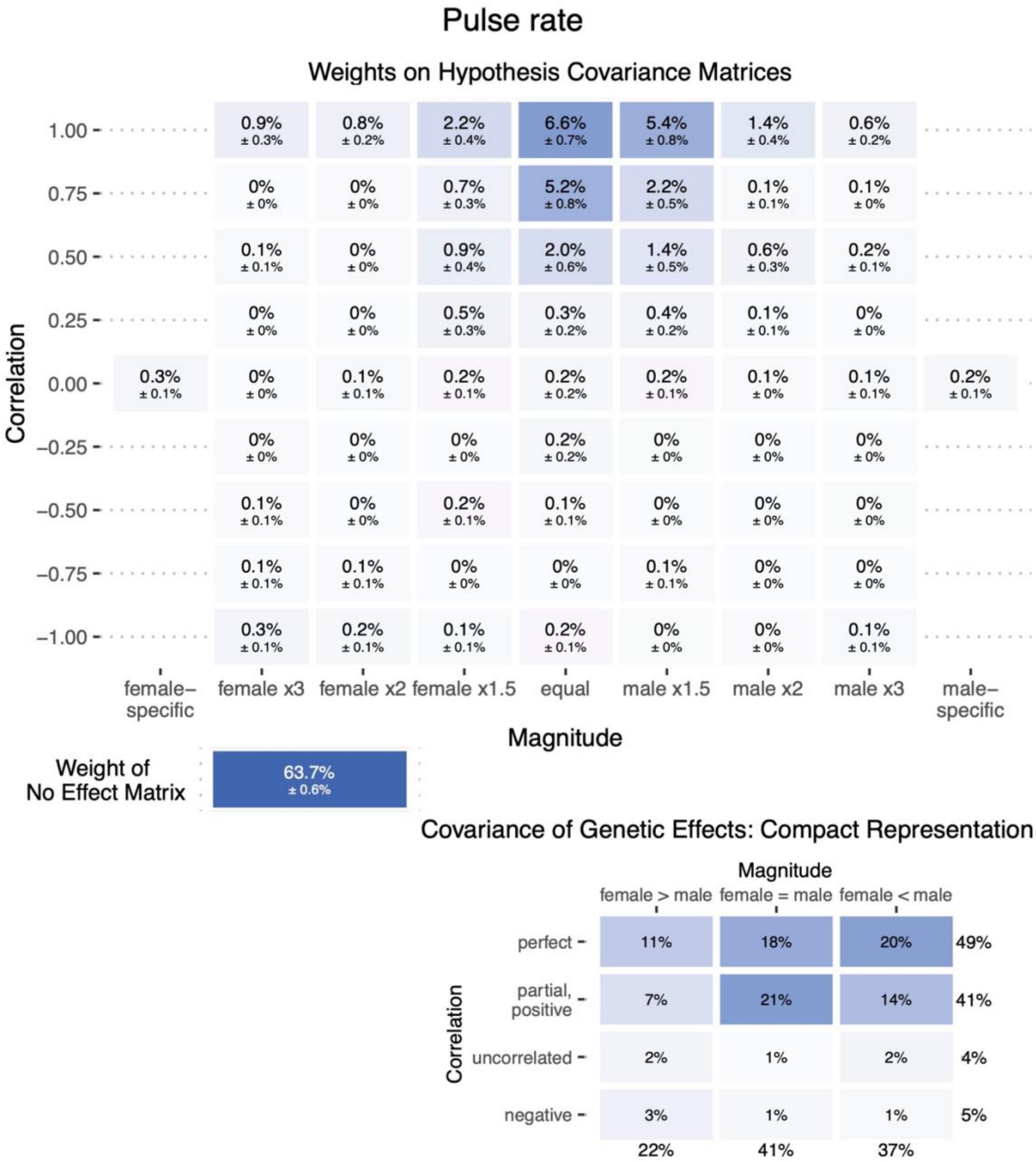

Pulse rate
Weights on Hypothesis Covariance Matrices
Correlation
1.00
0.75
0.50
0.25
0.00
−0.25
−0.50
−0.75
−1.00
1.00: 0.9% ± 0.3% | 0.8% ± 0.2% | 2.2% ± 0.4% | 6.6% ± 0.7% | 5.4% ± 0.8% | 1.4% ± 0.4% | 0.6% ± 0.2%
0.75: 0% ± 0% | 0% ± 0% | 0.7% ± 0.3% | 5.2% ± 0.8% | 2.2% ± 0.5% | 0.1% ± 0.1% | 0.1% ± 0%
0.50: 0.1% ± 0.1% | 0% ± 0% | 0.9% ± 0.4% | 2.0% ± 0.6% | 1.4% ± 0.5% | 0.6% ± 0.3% | 0.2% ± 0.1%
0.25: 0% ± 0% | 0% ± 0% | 0.5% ± 0.3% | 0.3% ± 0.2% | 0.4% ± 0.2% | 0.1% ± 0.1% | 0% ± 0%
0.00: 0.3% ± 0.1% | 0% ± 0% | 0.1% ± 0.1% | 0.2% ± 0.1% | 0.2% ± 0.2% | 0.2% ± 0.1% | 0.1% ± 0% | 0.1% ± 0.1% | 0.2% ± 0.1%
−0.25: 0% ± 0% | 0% ± 0% | 0% ± 0% | 0.2% ± 0.2% | 0% ± 0% | 0% ± 0% | 0% ± 0%
−0.50: 0.1% ± 0.1% | 0% ± 0% | 0.2% ± 0.1% | 0.1% ± 0.1% | 0% ± 0% | 0% ± 0% | 0% ± 0%
−0.75: 0.1% ± 0.1% | 0.1% ± 0.1% | 0% ± 0% | 0% ± 0% | 0.1% ± 0.1% | 0% ± 0% | 0% ± 0%
−1.00: 0.3% ± 0.1% | 0.2% ± 0.1% | 0.1% ± 0.1% | 0.2% ± 0.1% | 0% ± 0% | 0% ± 0% | 0.1% ± 0.1%
female– specific
female x3
female x2
female x1.5
equal
male x1.5
male x2
male x3
male– specific
Magnitude
Weight of No Effect Matrix
63.7% ± 0.6%
Covariance of Genetic Effects: Compact Representation
Magnitude
female > male
female = male
female < male
Correlation
perfect: 11% | 18% | 20% | 49%
partial, positive: 7% | 21% | 14% | 41%
uncorrelated: 2% | 1% | 2% | 4%
negative: 3% | 1% | 1% | 5%
22% | 41% | 37%

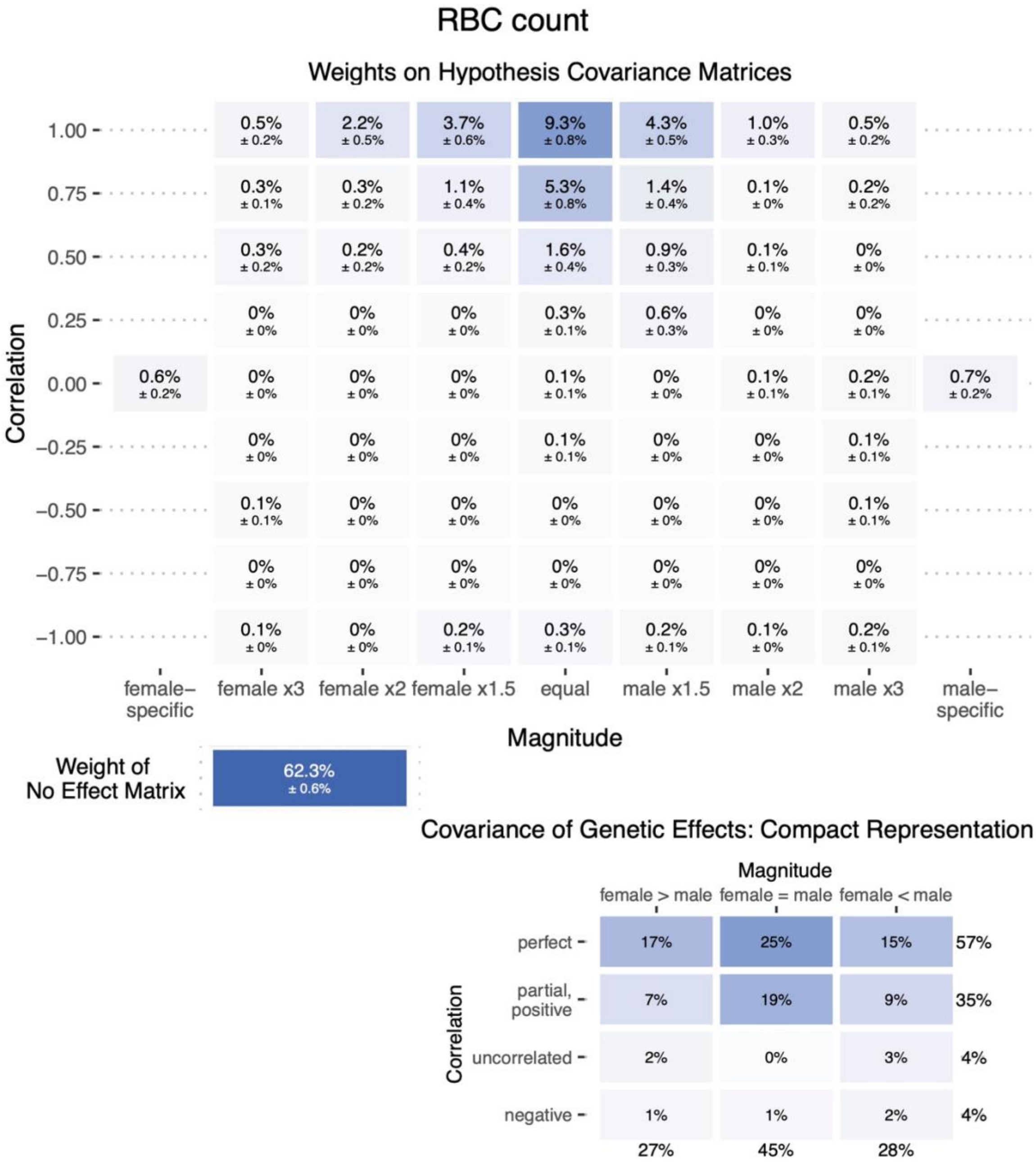
RBC count
Weights on Hypothesis Covariance Matrices
Correlation
1.00
0.75
0.50
0.25
0.00
−0.25
−0.50
−0.75
−1.00
0.5% ± 0.2% 2.2% ± 0.5% 3.7% ± 0.6% 9.3% ± 0.8% 4.3% ± 0.5% 1.0% ± 0.3% 0.5% ± 0.2%
0.3% ± 0.1% 0.3% ± 0.2% 1.1% ± 0.4% 5.3% ± 0.8% 1.4% ± 0.4% 0.1% ± 0% 0.2% ± 0.2%
0.3% ± 0.2% 0.2% ± 0.2% 0.4% ± 0.2% 1.6% ± 0.4% 0.9% ± 0.3% 0.1% ± 0.1% 0% ± 0%
0% ± 0% 0% ± 0% 0% ± 0% 0.3% ± 0.1% 0.6% ± 0.3% 0% ± 0% 0% ± 0%
0.6% ± 0.2% 0% ± 0% 0% ± 0% 0% ± 0% 0.1% ± 0.1% 0% ± 0% 0.1% ± 0.1% 0.2% ± 0.1% 0.7% ± 0.2%
0% ± 0% 0% ± 0% 0% ± 0% 0.1% ± 0.1% 0% ± 0% 0% ± 0% 0.1% ± 0.1%
0.1% ± 0.1% 0% ± 0% 0% ± 0% 0% ± 0% 0% ± 0% 0% ± 0% 0.1% ± 0.1%
0% ± 0% 0% ± 0% 0% ± 0% 0% ± 0% 0% ± 0% 0% ± 0% 0% ± 0%
0.1% ± 0% 0% ± 0% 0.2% ± 0.1% 0.3% ± 0.1% 0.2% ± 0.1% 0.1% ± 0% 0.2% ± 0.1%
female–specific
female x3
female x2
female x1.5
equal
male x1.5
male x2
male x3
male–specific
Magnitude
Weight of No Effect Matrix
62.3% ± 0.6%
Covariance of Genetic Effects: Compact Representation
Magnitude
female > male
female = male
female < male
Correlation
perfect 17% 25% 15% 57%
partial, positive 7% 19% 9% 35%
uncorrelated 2% 0% 3% 4%
negative 1% 1% 2% 4%
27% 45% 28%

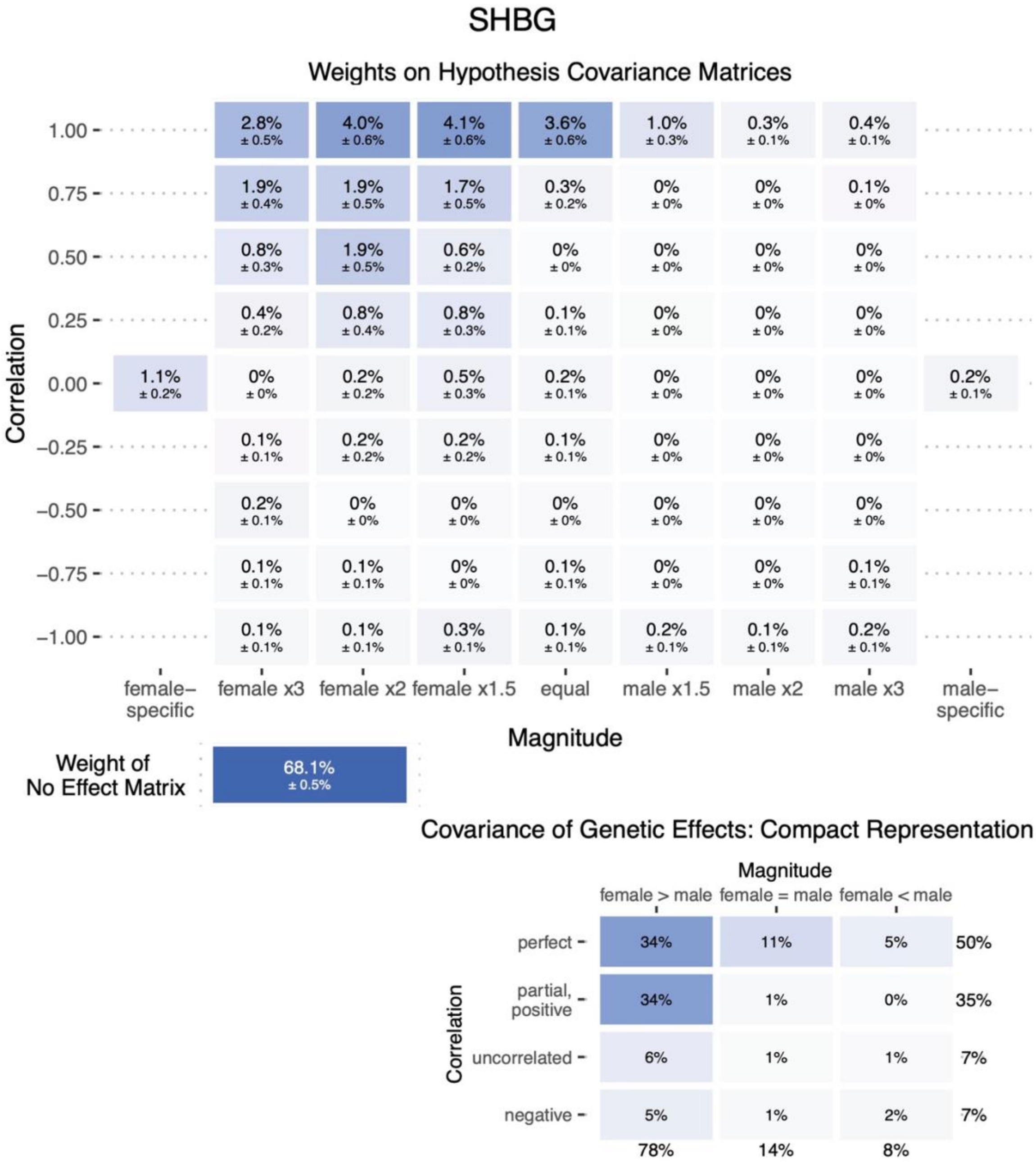

SHBG
Weights on Hypothesis Covariance Matrices
Correlation
Magnitude
female-specific
female x3
female x2
female x1.5
equal
male x1.5
male x2
male x3
male-specific
Weight of No Effect Matrix
68.1%
± 0.5%
Covariance of Genetic Effects: Compact Representation
Magnitude
female > male
female = male
female < male
perfect
partial, positive
uncorrelated
negative

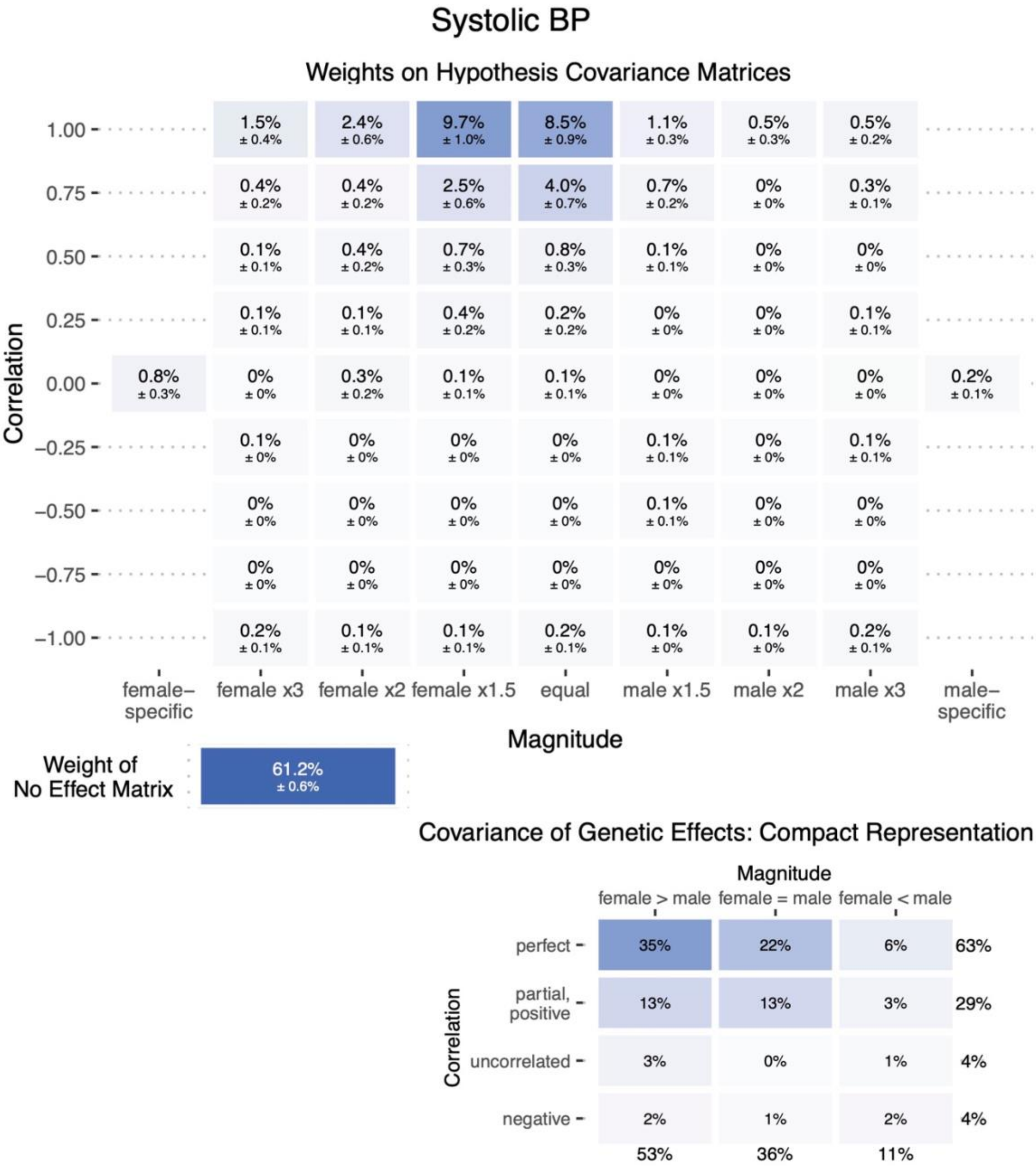
Systolic BP
Weights on Hypothesis Covariance Matrices
Correlation
1.00
0.75
0.50
0.25
0.00
−0.25
−0.50
−0.75
−1.00
1.00: 1.5% ± 0.4% | 2.4% ± 0.6% | 9.7% ± 1.0% | 8.5% ± 0.9% | 1.1% ± 0.3% | 0.5% ± 0.3% | 0.5% ± 0.2%
0.75: 0.4% ± 0.2% | 0.4% ± 0.2% | 2.5% ± 0.6% | 4.0% ± 0.7% | 0.7% ± 0.2% | 0% ± 0% | 0.3% ± 0.1%
0.50: 0.1% ± 0.1% | 0.4% ± 0.2% | 0.7% ± 0.3% | 0.8% ± 0.3% | 0.1% ± 0.1% | 0% ± 0% | 0% ± 0%
0.25: 0.1% ± 0.1% | 0.1% ± 0.1% | 0.4% ± 0.2% | 0.2% ± 0.2% | 0% ± 0% | 0% ± 0% | 0.1% ± 0.1%
0.00: 0.8% ± 0.3% | 0% ± 0% | 0.3% ± 0.2% | 0.1% ± 0.1% | 0.1% ± 0.1% | 0% ± 0% | 0% ± 0% | 0% ± 0% | 0.2% ± 0.1%
−0.25: 0.1% ± 0% | 0% ± 0% | 0% ± 0% | 0% ± 0% | 0.1% ± 0.1% | 0% ± 0% | 0.1% ± 0.1%
−0.50: 0% ± 0% | 0% ± 0% | 0% ± 0% | 0% ± 0% | 0.1% ± 0.1% | 0% ± 0% | 0% ± 0%
−0.75: 0% ± 0% | 0% ± 0% | 0% ± 0% | 0% ± 0% | 0% ± 0% | 0% ± 0% | 0% ± 0%
−1.00: 0.2% ± 0.1% | 0.1% ± 0.1% | 0.1% ± 0.1% | 0.2% ± 0.1% | 0.1% ± 0% | 0.1% ± 0% | 0.2% ± 0.1%
female–specific
female x3
female x2
female x1.5
equal
male x1.5
male x2
male x3
male–specific
Magnitude
Weight of No Effect Matrix
61.2% ± 0.6%
Covariance of Genetic Effects: Compact Representation
Magnitude
female > male
female = male
female < male
Correlation
perfect: 35% | 22% | 6% | 63%
partial, positive: 13% | 13% | 3% | 29%
uncorrelated: 3% | 0% | 1% | 4%
negative: 2% | 1% | 2% | 4%
53% | 36% | 11%

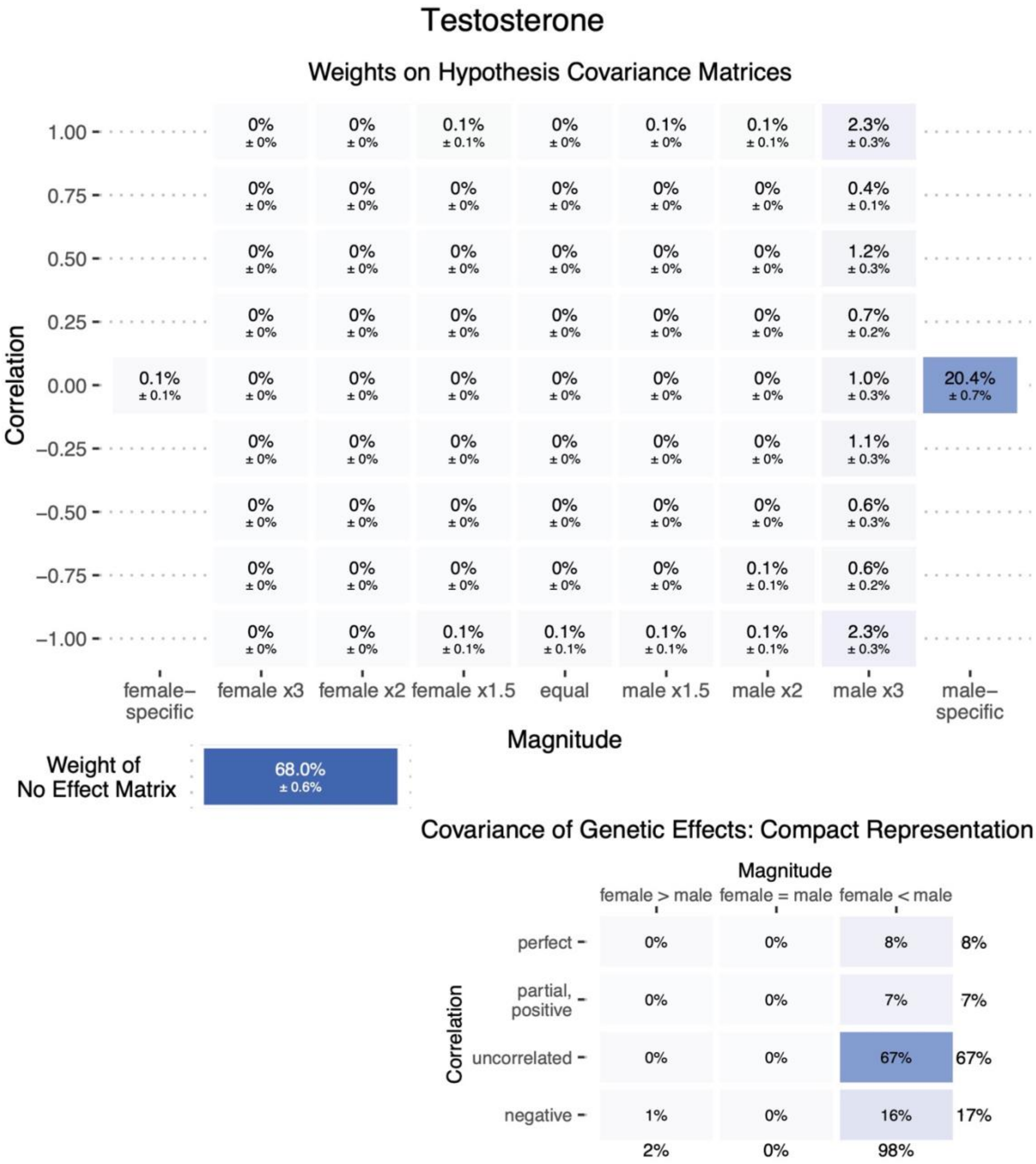

Testosterone
Weights on Hypothesis Covariance Matrices
Correlation
1.00
0.75
0.50
0.25
0.00
−0.25
−0.50
−0.75
−1.00
1.00: 0% ± 0% | 0% ± 0% | 0.1% ± 0.1% | 0% ± 0% | 0.1% ± 0% | 0.1% ± 0.1% | 2.3% ± 0.3%
0.75: 0% ± 0% | 0% ± 0% | 0% ± 0% | 0% ± 0% | 0% ± 0% | 0% ± 0% | 0.4% ± 0.1%
0.50: 0% ± 0% | 0% ± 0% | 0% ± 0% | 0% ± 0% | 0% ± 0% | 0% ± 0% | 1.2% ± 0.3%
0.25: 0% ± 0% | 0% ± 0% | 0% ± 0% | 0% ± 0% | 0% ± 0% | 0% ± 0% | 0.7% ± 0.2%
0.00: 0.1% ± 0.1% | 0% ± 0% | 0% ± 0% | 0% ± 0% | 0% ± 0% | 0% ± 0% | 0% ± 0% | 1.0% ± 0.3% | 20.4% ± 0.7%
−0.25: 0% ± 0% | 0% ± 0% | 0% ± 0% | 0% ± 0% | 0% ± 0% | 0% ± 0% | 1.1% ± 0.3%
−0.50: 0% ± 0% | 0% ± 0% | 0% ± 0% | 0% ± 0% | 0% ± 0% | 0% ± 0% | 0.6% ± 0.3%
−0.75: 0% ± 0% | 0% ± 0% | 0% ± 0% | 0% ± 0% | 0% ± 0% | 0.1% ± 0.1% | 0.6% ± 0.2%
−1.00: 0% ± 0% | 0% ± 0% | 0.1% ± 0.1% | 0.1% ± 0.1% | 0.1% ± 0.1% | 0.1% ± 0.1% | 2.3% ± 0.3%
female–specific
female x3
female x2
female x1.5
equal
male x1.5
male x2
male x3
male–specific
Magnitude
Weight of No Effect Matrix
68.0% ± 0.6%
Covariance of Genetic Effects: Compact Representation
Magnitude
female > male
female = male
female < male
Correlation
perfect: 0% | 0% | 8% | 8%
partial, positive: 0% | 0% | 7% | 7%
uncorrelated: 0% | 0% | 67% | 67%
negative: 1% | 0% | 16% | 17%
2% | 0% | 98%

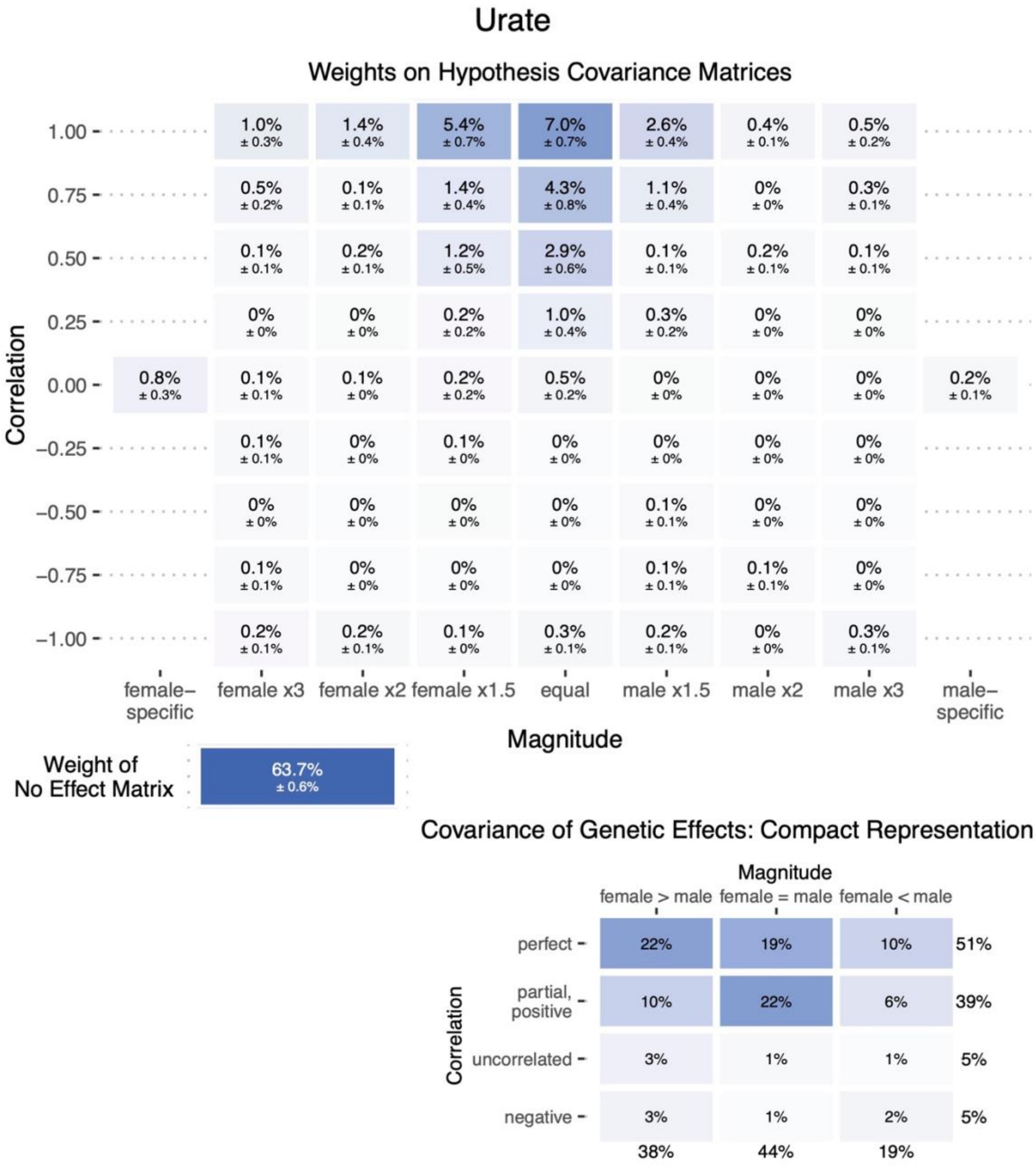

Urate
Weights on Hypothesis Covariance Matrices
Correlation
1.00
0.75
0.50
0.25
0.00
−0.25
−0.50
−0.75
−1.00
1.0% ± 0.3%  1.4% ± 0.4%  5.4% ± 0.7%  7.0% ± 0.7%  2.6% ± 0.4%  0.4% ± 0.1%  0.5% ± 0.2%
0.5% ± 0.2%  0.1% ± 0.1%  1.4% ± 0.4%  4.3% ± 0.8%  1.1% ± 0.4%  0% ± 0%  0.3% ± 0.1%
0.1% ± 0.1%  0.2% ± 0.1%  1.2% ± 0.5%  2.9% ± 0.6%  0.1% ± 0.1%  0.2% ± 0.1%  0.1% ± 0.1%
0% ± 0%  0% ± 0%  0.2% ± 0.2%  1.0% ± 0.4%  0.3% ± 0.2%  0% ± 0%  0% ± 0%
0.8% ± 0.3%  0.1% ± 0.1%  0.1% ± 0%  0.2% ± 0.2%  0.5% ± 0.2%  0% ± 0%  0% ± 0%  0% ± 0%  0.2% ± 0.1%
0.1% ± 0.1%  0% ± 0%  0.1% ± 0%  0% ± 0%  0% ± 0%  0% ± 0%  0% ± 0%
0% ± 0%  0% ± 0%  0% ± 0%  0% ± 0%  0.1% ± 0.1%  0% ± 0%  0% ± 0%
0.1% ± 0.1%  0% ± 0%  0% ± 0%  0% ± 0%  0.1% ± 0.1%  0.1% ± 0.1%  0% ± 0%
0.2% ± 0.1%  0.2% ± 0.1%  0.1% ± 0%  0.3% ± 0.1%  0.2% ± 0.1%  0% ± 0%  0.3% ± 0.1%
female–specific
female x3
female x2
female x1.5
equal
male x1.5
male x2
male x3
male–specific
Magnitude
Weight of No Effect Matrix
63.7% ± 0.6%
Covariance of Genetic Effects: Compact Representation
Magnitude
female > male
female = male
female < male
Correlation
perfect  22%  19%  10%  51%
partial, positive  10%  22%  6%  39%
uncorrelated  3%  1%  1%  5%
negative  3%  1%  2%  5%
38%  44%  19%

#### Urea

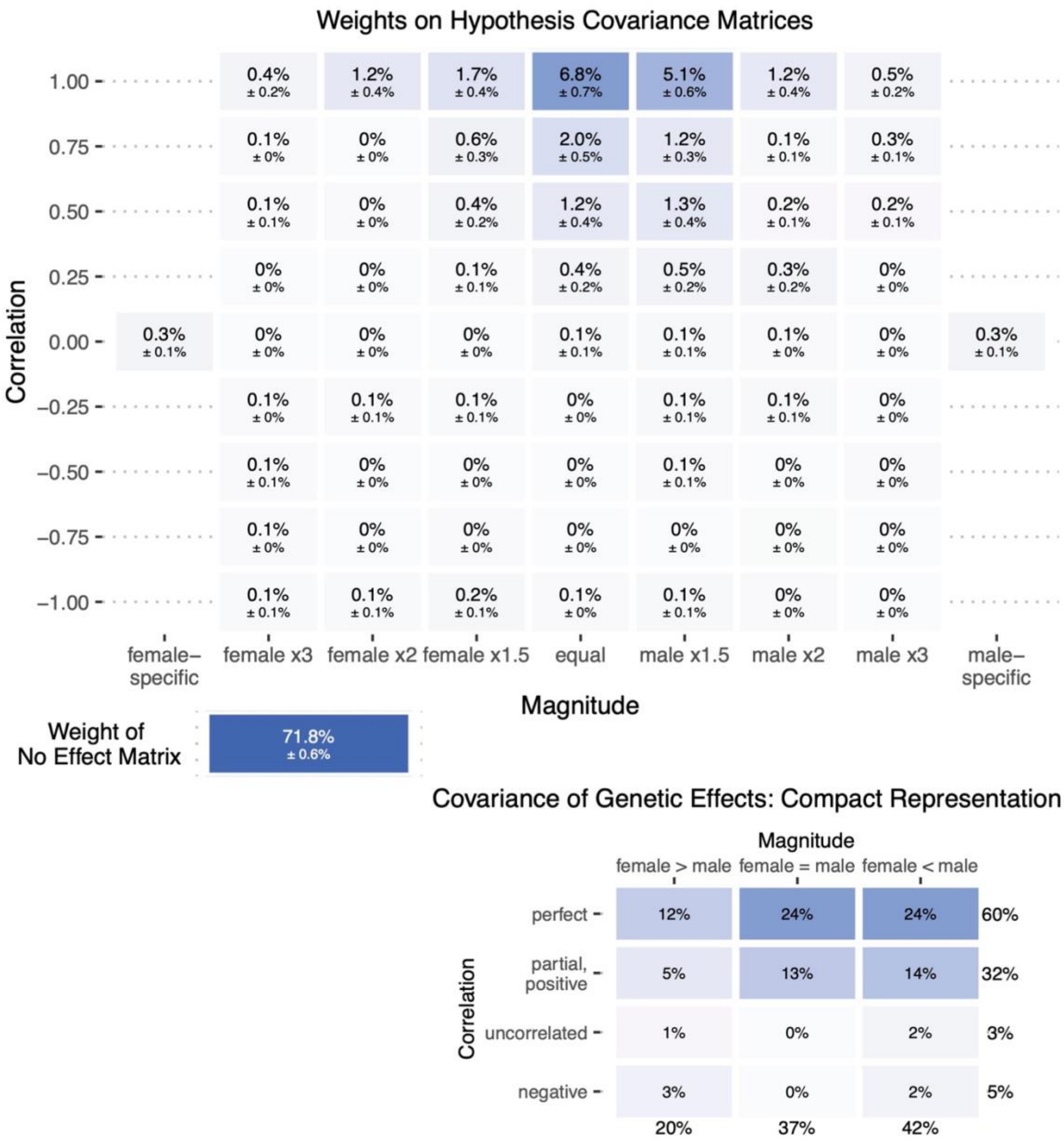
Weights on Hypothesis Covariance Matrices
Correlation
1.00
0.75
0.50
0.25
0.00
−0.25
−0.50
−0.75
−1.00
0.4% ± 0.2% | 1.2% ± 0.4% | 1.7% ± 0.4% | 6.8% ± 0.7% | 5.1% ± 0.6% | 1.2% ± 0.4% | 0.5% ± 0.2%
0.1% ± 0% | 0% ± 0% | 0.6% ± 0.3% | 2.0% ± 0.5% | 1.2% ± 0.3% | 0.1% ± 0.1% | 0.3% ± 0.1%
0.1% ± 0.1% | 0% ± 0% | 0.4% ± 0.2% | 1.2% ± 0.4% | 1.3% ± 0.4% | 0.2% ± 0.1% | 0.2% ± 0.1%
0% ± 0% | 0% ± 0% | 0.1% ± 0.1% | 0.4% ± 0.2% | 0.5% ± 0.2% | 0.3% ± 0.2% | 0% ± 0%
0.3% ± 0.1% | 0% ± 0% | 0% ± 0% | 0% ± 0% | 0.1% ± 0.1% | 0.1% ± 0.1% | 0.1% ± 0% | 0% ± 0% | 0.3% ± 0.1%
0.1% ± 0% | 0.1% ± 0.1% | 0.1% ± 0.1% | 0% ± 0% | 0.1% ± 0.1% | 0.1% ± 0.1% | 0% ± 0%
0.1% ± 0.1% | 0% ± 0% | 0% ± 0% | 0% ± 0% | 0.1% ± 0.1% | 0% ± 0% | 0% ± 0%
0.1% ± 0% | 0% ± 0% | 0% ± 0% | 0% ± 0% | 0% ± 0% | 0% ± 0% | 0% ± 0%
0.1% ± 0.1% | 0.1% ± 0.1% | 0.2% ± 0.1% | 0.1% ± 0% | 0.1% ± 0.1% | 0% ± 0% | 0% ± 0%
female–specific
female x3
female x2
female x1.5
equal
male x1.5
male x2
male x3
male–specific
Magnitude
Weight of No Effect Matrix
71.8% ± 0.6%
Covariance of Genetic Effects: Compact Representation
Magnitude
female > male
female = male
female < male
Correlation
perfect | 12% | 24% | 24% | 60%
partial, positive | 5% | 13% | 14% | 32%
uncorrelated | 1% | 0% | 2% | 3%
negative | 3% | 0% | 2% | 5%
20% | 37% | 42%

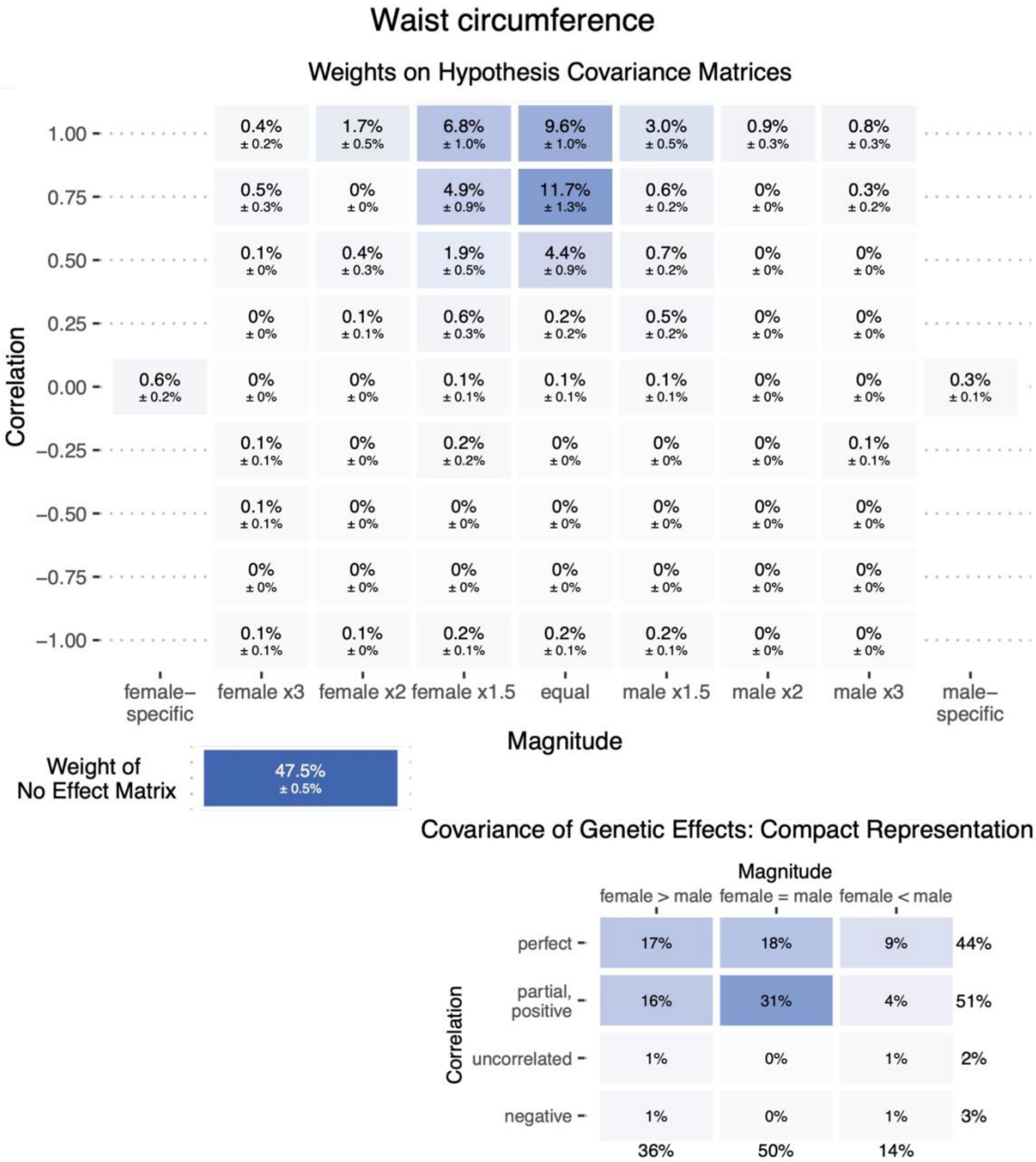

Waist circumference
Weights on Hypothesis Covariance Matrices
Correlation
1.00
0.4% ± 0.2%
1.7% ± 0.5%
6.8% ± 1.0%
9.6% ± 1.0%
3.0% ± 0.5%
0.9% ± 0.3%
0.8% ± 0.3%
0.75
0.5% ± 0.3%
0% ± 0%
4.9% ± 0.9%
11.7% ± 1.3%
0.6% ± 0.2%
0% ± 0%
0.3% ± 0.2%
0.50
0.1% ± 0%
0.4% ± 0.3%
1.9% ± 0.5%
4.4% ± 0.9%
0.7% ± 0.2%
0% ± 0%
0% ± 0%
0.25
0% ± 0%
0.1% ± 0.1%
0.6% ± 0.3%
0.2% ± 0.2%
0.5% ± 0.2%
0% ± 0%
0% ± 0%
0.00
0.6% ± 0.2%
0% ± 0%
0% ± 0%
0.1% ± 0.1%
0.1% ± 0.1%
0.1% ± 0.1%
0% ± 0%
0% ± 0%
0.3% ± 0.1%
−0.25
0.1% ± 0.1%
0% ± 0%
0.2% ± 0.2%
0% ± 0%
0% ± 0%
0% ± 0%
0.1% ± 0.1%
−0.50
0.1% ± 0.1%
0% ± 0%
0% ± 0%
0% ± 0%
0% ± 0%
0% ± 0%
0% ± 0%
−0.75
0% ± 0%
0% ± 0%
0% ± 0%
0% ± 0%
0% ± 0%
0% ± 0%
0% ± 0%
−1.00
0.1% ± 0.1%
0.1% ± 0%
0.2% ± 0.1%
0.2% ± 0.1%
0.2% ± 0.1%
0% ± 0%
0% ± 0%
female–specific
female x3
female x2
female x1.5
equal
male x1.5
male x2
male x3
male–specific
Magnitude
Weight of No Effect Matrix
47.5% ± 0.5%
Covariance of Genetic Effects: Compact Representation
Magnitude
female > male
female = male
female < male
Correlation
perfect
17%
18%
9%
44%
partial, positive
16%
31%
4%
51%
uncorrelated
1%
0%
1%
2%
negative
1%
0%
1%
3%
36%
50%
14%

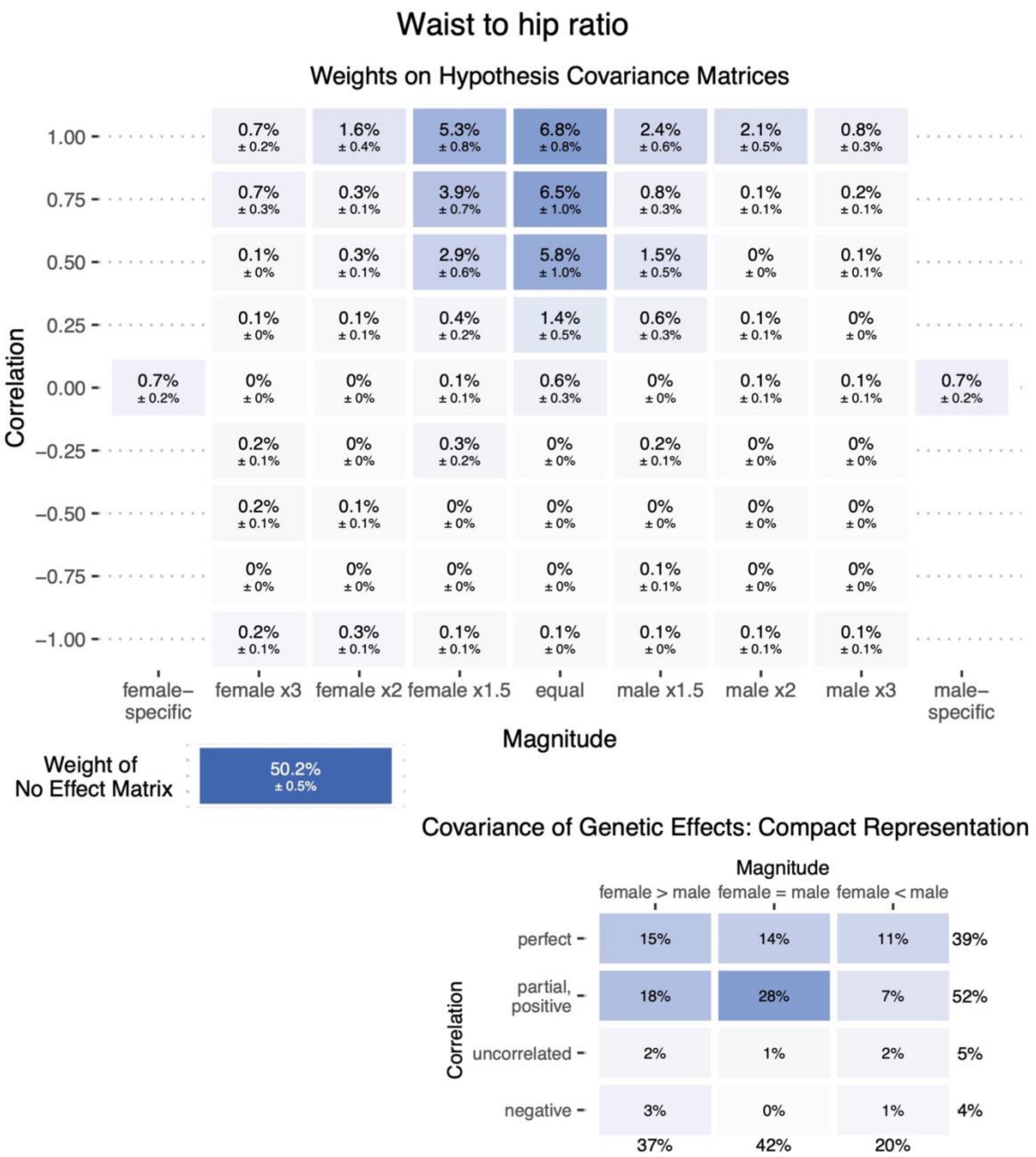
Waist to hip ratio
Weights on Hypothesis Covariance Matrices
Correlation
Magnitude
female– specific
female x3
female x2
female x1.5
equal
male x1.5
male x2
male x3
male– specific
Weight of No Effect Matrix
50.2% ± 0.5%
Covariance of Genetic Effects: Compact Representation
female > male
female = male
female < male
perfect
partial, positive
uncorrelated
negative

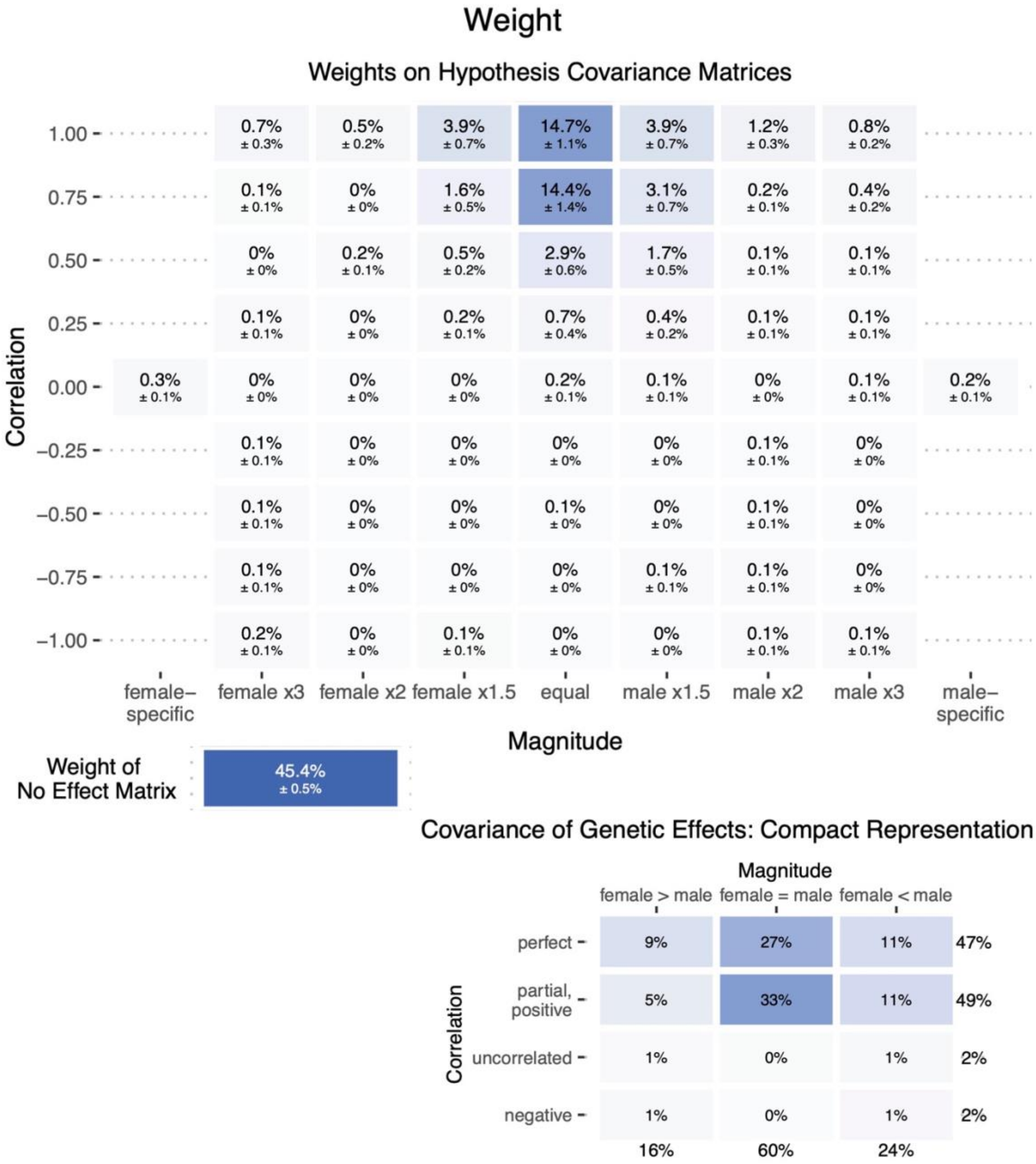
Weight
Weights on Hypothesis Covariance Matrices
Correlation
1.00
0.75
0.50
0.25
0.00
−0.25
−0.50
−0.75
−1.00
0.7% ± 0.3%
0.5% ± 0.2%
3.9% ± 0.7%
14.7% ± 1.1%
3.9% ± 0.7%
1.2% ± 0.3%
0.8% ± 0.2%
0.1% ± 0.1%
0% ± 0%
1.6% ± 0.5%
14.4% ± 1.4%
3.1% ± 0.7%
0.2% ± 0.1%
0.4% ± 0.2%
0% ± 0%
0.2% ± 0.1%
0.5% ± 0.2%
2.9% ± 0.6%
1.7% ± 0.5%
0.1% ± 0.1%
0.1% ± 0.1%
0.1% ± 0.1%
0% ± 0%
0.2% ± 0.1%
0.7% ± 0.4%
0.4% ± 0.2%
0.1% ± 0.1%
0.1% ± 0.1%
0.3% ± 0.1%
0% ± 0%
0% ± 0%
0% ± 0%
0.2% ± 0.1%
0.1% ± 0.1%
0% ± 0%
0.1% ± 0.1%
0.2% ± 0.1%
0.1% ± 0.1%
0% ± 0%
0% ± 0%
0% ± 0%
0% ± 0%
0.1% ± 0.1%
0% ± 0%
0.1% ± 0.1%
0% ± 0%
0% ± 0%
0.1% ± 0%
0% ± 0%
0.1% ± 0.1%
0% ± 0%
0.1% ± 0.1%
0% ± 0%
0% ± 0%
0% ± 0%
0.1% ± 0.1%
0.1% ± 0.1%
0% ± 0%
0.2% ± 0.1%
0% ± 0%
0.1% ± 0.1%
0% ± 0%
0% ± 0%
0.1% ± 0.1%
0.1% ± 0.1%
female–specific
female x3
female x2
female x1.5
equal
male x1.5
male x2
male x3
male–specific
Magnitude
Weight of No Effect Matrix
45.4% ± 0.5%
Covariance of Genetic Effects: Compact Representation
Magnitude
female > male
female = male
female < male
Correlation
perfect
partial, positive
uncorrelated
negative
9%
27%
11%
47%
5%
33%
11%
49%
1%
0%
1%
2%
1%
0%
1%
2%
16%
60%
24%

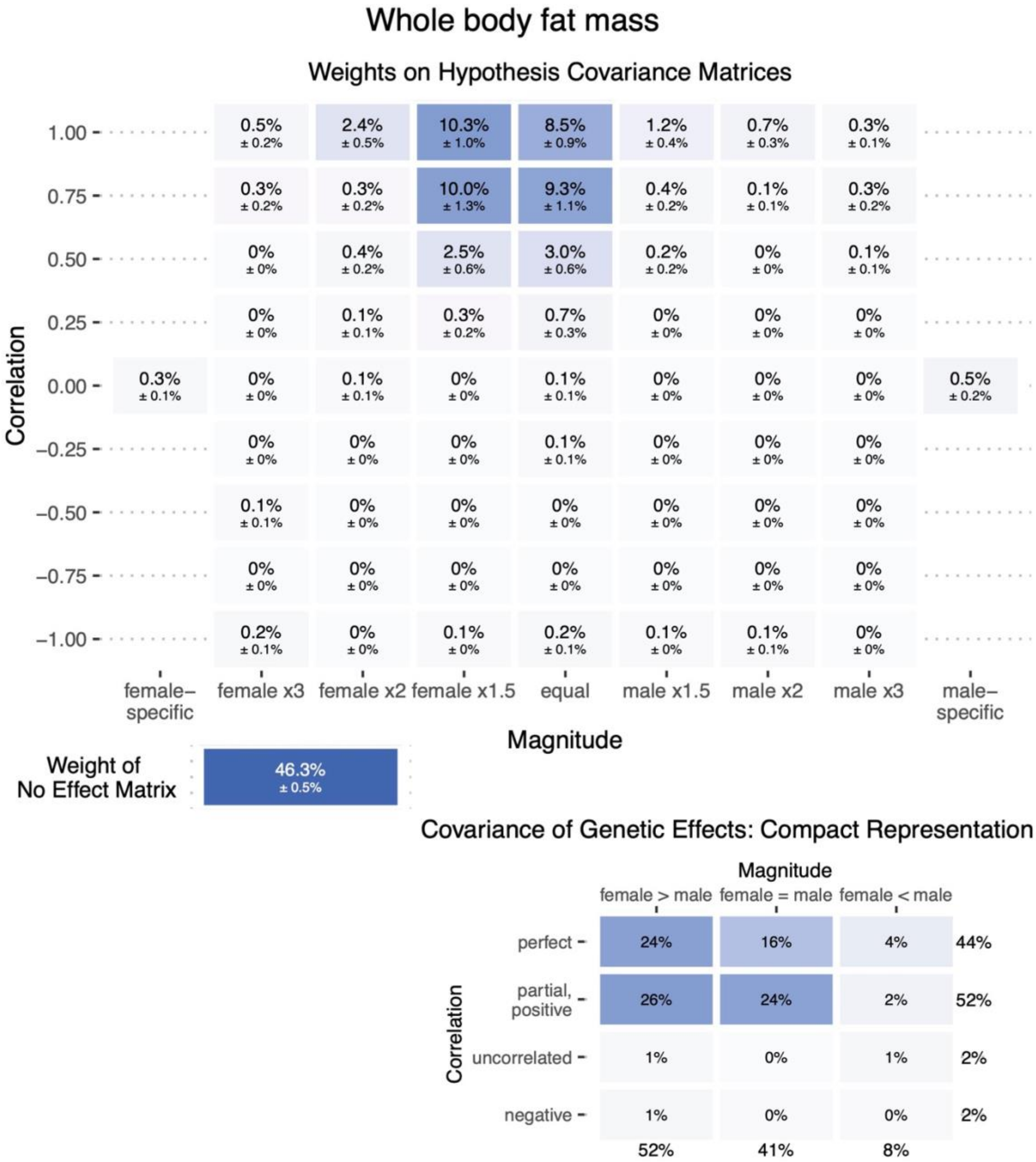

Whole body fat mass
Weights on Hypothesis Covariance Matrices
Correlation
1.00
0.75
0.50
0.25
0.00
−0.25
−0.50
−0.75
−1.00
0.5% ± 0.2%
2.4% ± 0.5%
10.3% ± 1.0%
8.5% ± 0.9%
1.2% ± 0.4%
0.7% ± 0.3%
0.3% ± 0.1%
0.3% ± 0.2%
0.3% ± 0.2%
10.0% ± 1.3%
9.3% ± 1.1%
0.4% ± 0.2%
0.1% ± 0.1%
0.3% ± 0.2%
0% ± 0%
0.4% ± 0.2%
2.5% ± 0.6%
3.0% ± 0.6%
0.2% ± 0.2%
0% ± 0%
0.1% ± 0.1%
0% ± 0%
0.1% ± 0.1%
0.3% ± 0.2%
0.7% ± 0.3%
0% ± 0%
0% ± 0%
0% ± 0%
0.3% ± 0.1%
0% ± 0%
0.1% ± 0.1%
0% ± 0%
0.1% ± 0.1%
0% ± 0%
0% ± 0%
0% ± 0%
0.5% ± 0.2%
0% ± 0%
0% ± 0%
0% ± 0%
0.1% ± 0.1%
0% ± 0%
0% ± 0%
0% ± 0%
0.1% ± 0.1%
0% ± 0%
0% ± 0%
0% ± 0%
0% ± 0%
0% ± 0%
0% ± 0%
0% ± 0%
0% ± 0%
0% ± 0%
0% ± 0%
0% ± 0%
0% ± 0%
0% ± 0%
0% ± 0%
0.2% ± 0.1%
0% ± 0%
0.1% ± 0%
0.2% ± 0.1%
0.1% ± 0%
0.1% ± 0.1%
0% ± 0%
female–specific
female x3
female x2
female x1.5
equal
male x1.5
male x2
male x3
male–specific
Magnitude
Weight of No Effect Matrix
46.3% ± 0.5%
Covariance of Genetic Effects: Compact Representation
Magnitude
female > male
female = male
female < male
Correlation
perfect
partial, positive
uncorrelated
negative
24%
16%
4%
44%
26%
24%
2%
52%
1%
0%
1%
2%
1%
0%
0%
2%
52%
41%
8%

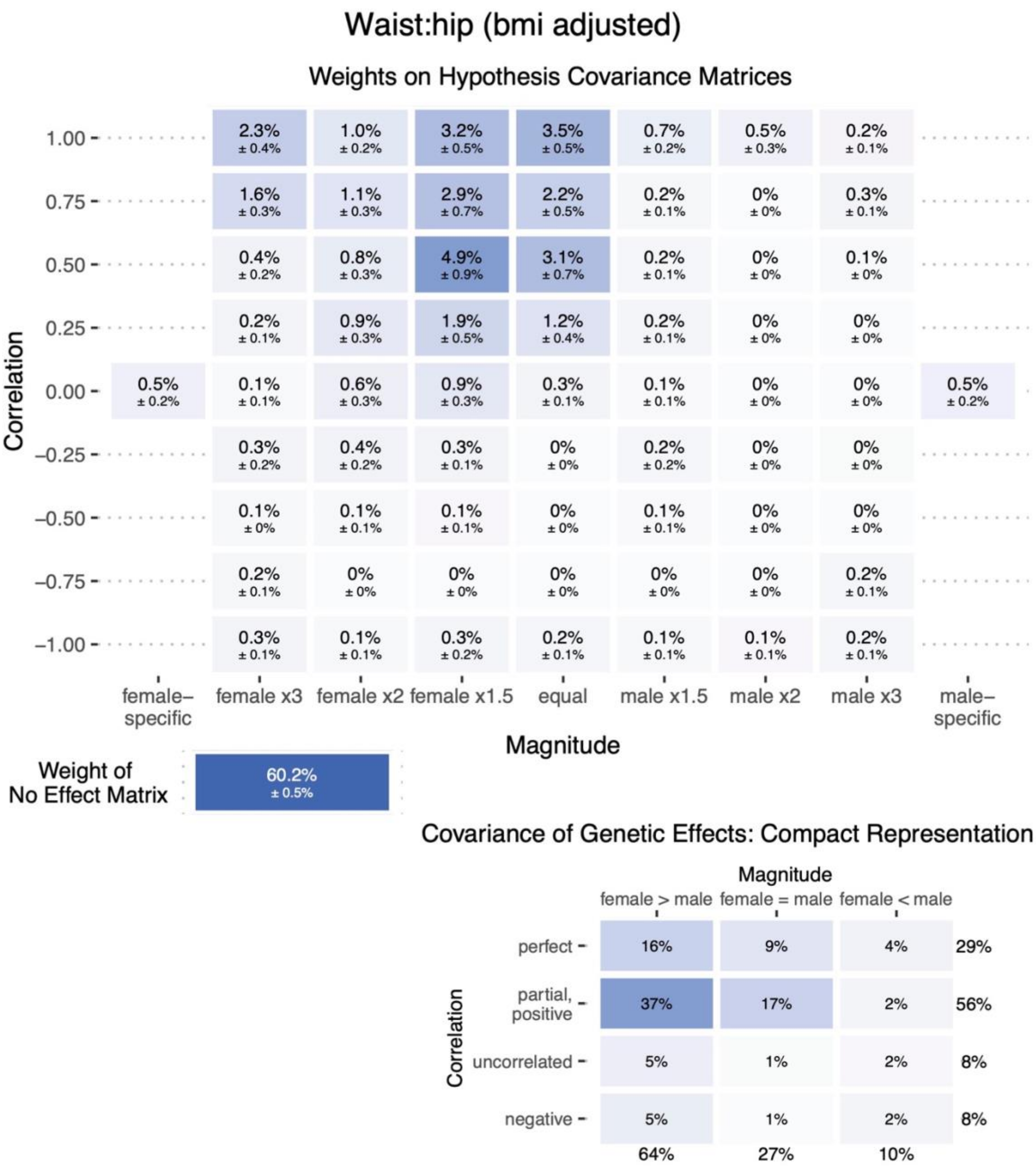
Waist:hip (bmi adjusted)
Weights on Hypothesis Covariance Matrices
Correlation
1.00
0.75
0.50
0.25
0.00
−0.25
−0.50
−0.75
−1.00
2.3% ± 0.4%
1.0% ± 0.2%
3.2% ± 0.5%
3.5% ± 0.5%
0.7% ± 0.2%
0.5% ± 0.3%
0.2% ± 0.1%
1.6% ± 0.3%
1.1% ± 0.3%
2.9% ± 0.7%
2.2% ± 0.5%
0.2% ± 0.1%
0% ± 0%
0.3% ± 0.1%
0.4% ± 0.2%
0.8% ± 0.3%
4.9% ± 0.9%
3.1% ± 0.7%
0.2% ± 0.1%
0% ± 0%
0.1% ± 0%
0.2% ± 0.1%
0.9% ± 0.3%
1.9% ± 0.5%
1.2% ± 0.4%
0.2% ± 0.1%
0% ± 0%
0% ± 0%
0.5% ± 0.2%
0.1% ± 0.1%
0.6% ± 0.3%
0.9% ± 0.3%
0.3% ± 0.1%
0.1% ± 0.1%
0% ± 0%
0% ± 0%
0.5% ± 0.2%
0.3% ± 0.2%
0.4% ± 0.2%
0.3% ± 0.1%
0% ± 0%
0.2% ± 0.2%
0% ± 0%
0% ± 0%
0.1% ± 0%
0.1% ± 0.1%
0.1% ± 0.1%
0% ± 0%
0.1% ± 0.1%
0% ± 0%
0% ± 0%
0.2% ± 0.1%
0% ± 0%
0% ± 0%
0% ± 0%
0% ± 0%
0% ± 0%
0.2% ± 0.1%
0.3% ± 0.1%
0.1% ± 0.1%
0.3% ± 0.2%
0.2% ± 0.1%
0.1% ± 0.1%
0.1% ± 0.1%
0.2% ± 0.1%
female–specific
female x3
female x2
female x1.5
equal
male x1.5
male x2
male x3
male–specific
Magnitude
Weight of No Effect Matrix
60.2% ± 0.5%
Covariance of Genetic Effects: Compact Representation
Magnitude
female > male
female = male
female < male
Correlation
perfect
partial, positive
uncorrelated
negative
16%
9%
4%
29%
37%
17%
2%
56%
5%
1%
2%
8%
5%
1%
2%
8%
64%
27%
10%

**Figure S6: Simulation of sex-invariant genetic effects but with variable environmental variance, related to Figure 3**

Mixture weights based on various parameters for number of causal SNPs, male-specific heritability, and female to male environmental variance ratio are shown for all hypothesis covariance matrices represented by the axes. Results are generated from a simulation study based on **Eq. 2**, in which environmental variance differs between the sexes while genetic effects stay the same to test whether *mash* interprets the environmental variance difference as GxSex (**Methods "Environmental variance simulation for *mash*"**). We tested three parameters for female to male environmental variance ratio [1, 1.5, 5]. Parameters for the number of causal SNPs [100, 1K, 10K] and male heritability [0.01, 0.1, 0.5] are listed on the top right.

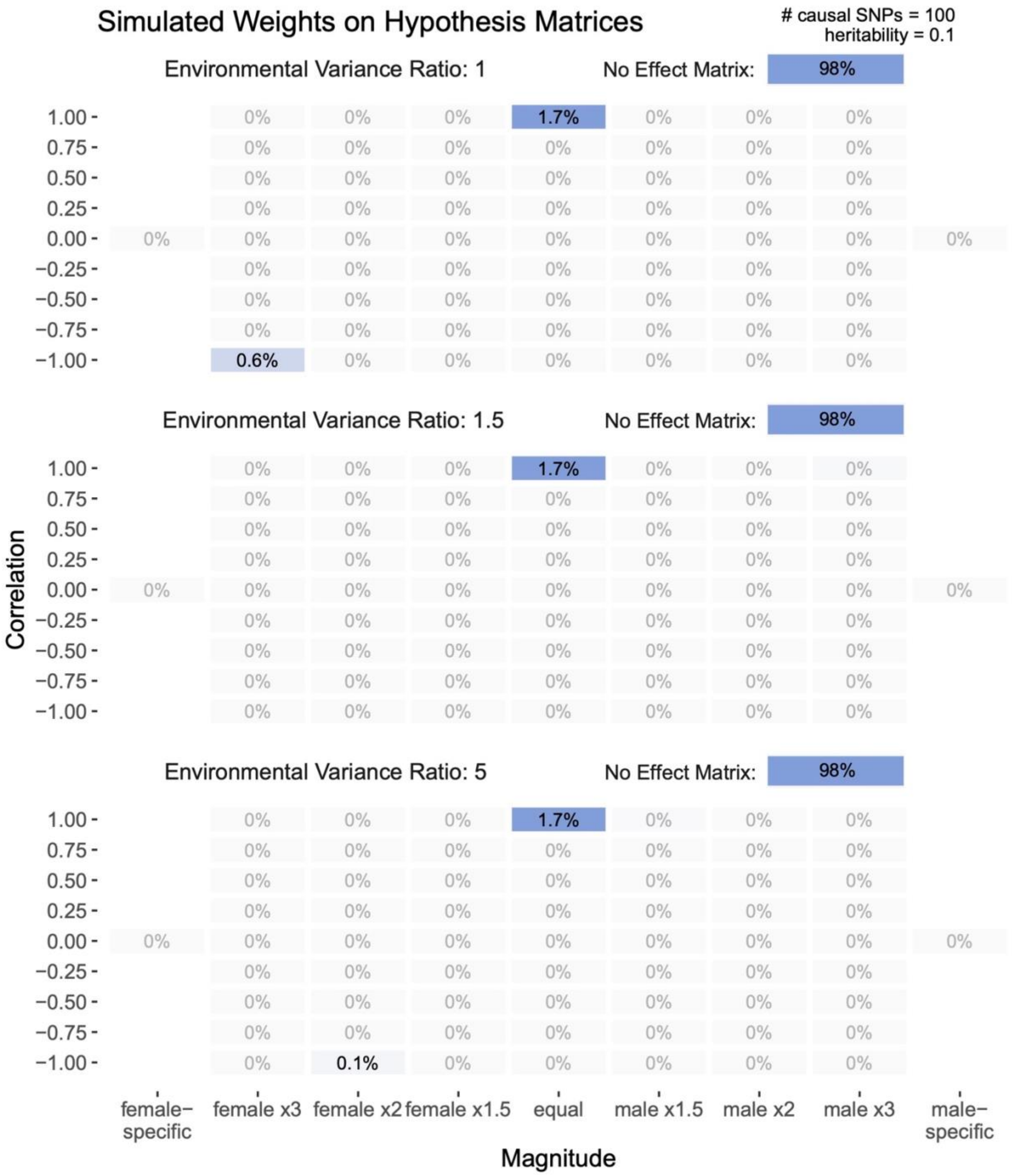
Simulated Weights on Hypothesis Matrices
### causal SNPs = 100
heritability = 0.1
Environmental Variance Ratio: 1
No Effect Matrix: 98%
Environmental Variance Ratio: 1.5
No Effect Matrix: 98%
Environmental Variance Ratio: 5
No Effect Matrix: 98%
Correlation
Magnitude
female−specific
female x3
female x2
female x1.5
equal
male x1.5
male x2
male x3
male−specific

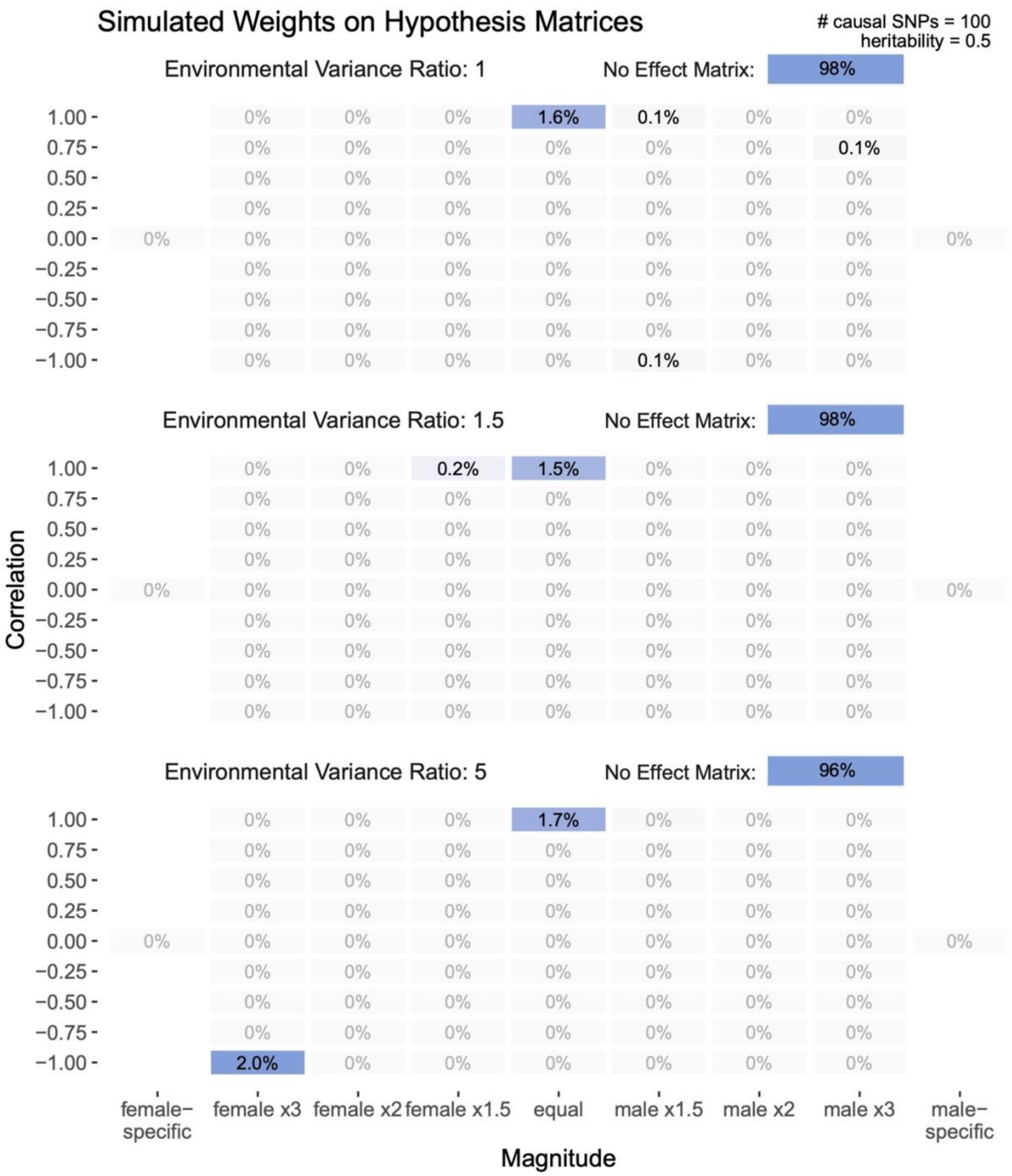
Simulated Weights on Hypothesis Matrices
### causal SNPs = 100
heritability = 0.5
Environmental Variance Ratio: 1
No Effect Matrix:
98%
1.6%
0.1%
0.1%
0.1%
Environmental Variance Ratio: 1.5
No Effect Matrix:
98%
0.2%
1.5%
Environmental Variance Ratio: 5
No Effect Matrix:
96%
1.7%
2.0%
0%
1.00
0.75
0.50
0.25
0.00
−0.25
−0.50
−0.75
−1.00
Correlation
female−specific
female x3
female x2
female x1.5
equal
male x1.5
male x2
male x3
male−specific
Magnitude

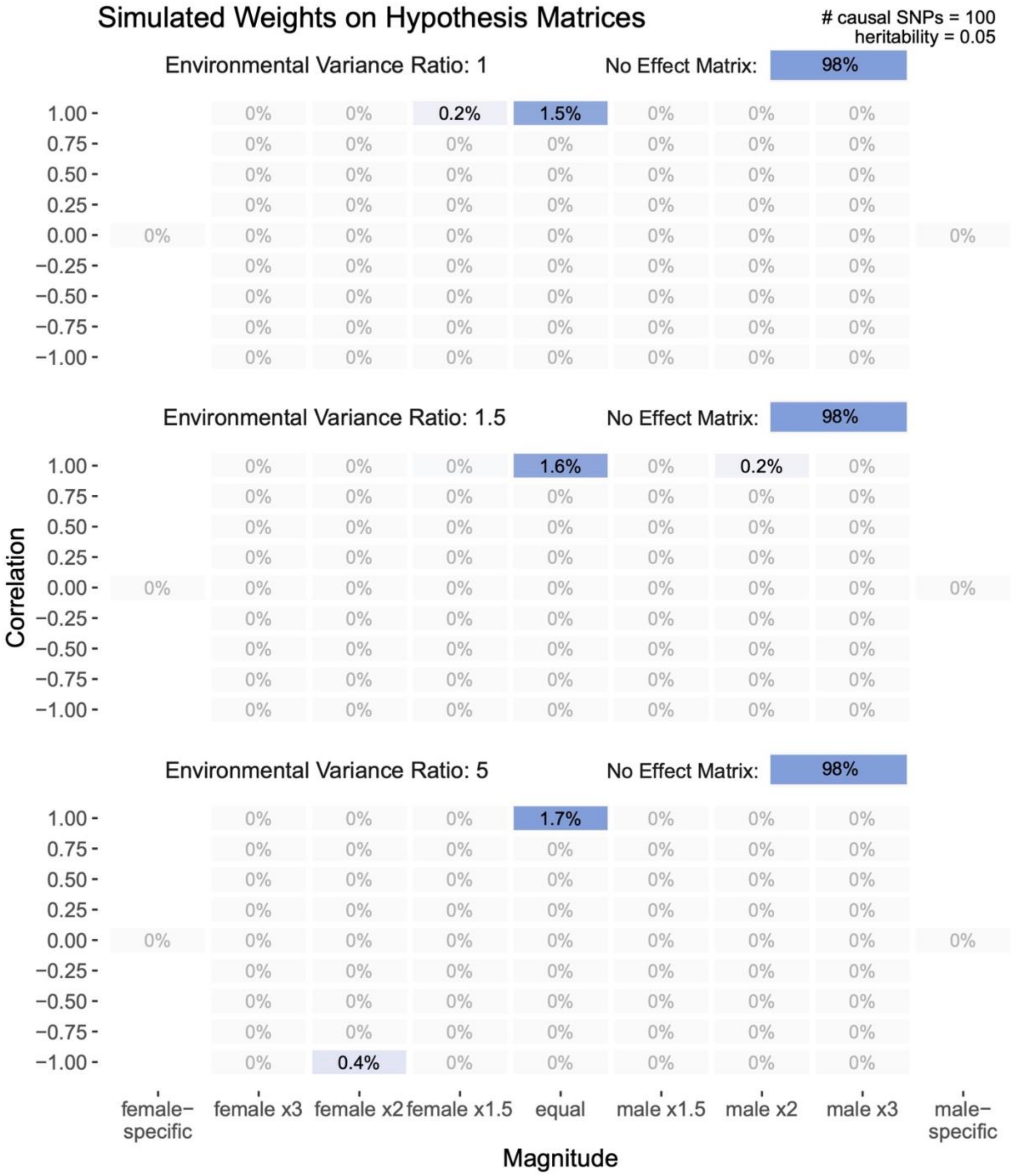
Simulated Weights on Hypothesis Matrices
### causal SNPs = 100
heritability = 0.05
Environmental Variance Ratio: 1
No Effect Matrix: 98%
Environmental Variance Ratio: 1.5
No Effect Matrix: 98%
Environmental Variance Ratio: 5
No Effect Matrix: 98%
Correlation
Magnitude
female−specific
female x3
female x2
female x1.5
equal
male x1.5
male x2
male x3
male−specific

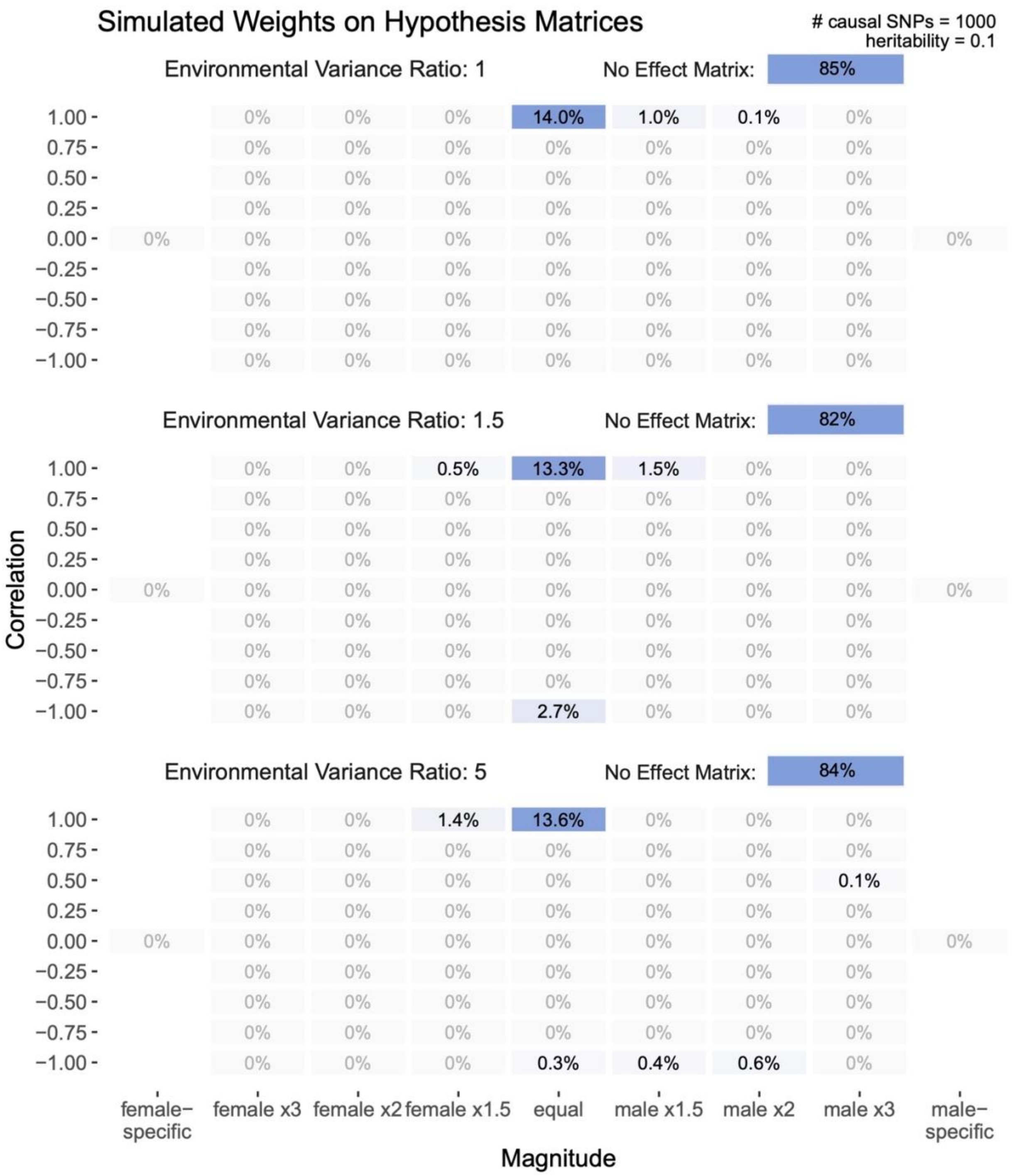
Simulated Weights on Hypothesis Matrices
### causal SNPs = 1000
heritability = 0.1
Environmental Variance Ratio: 1
No Effect Matrix: 85%
14.0%
1.0%
0.1%
Environmental Variance Ratio: 1.5
No Effect Matrix: 82%
0.5%
13.3%
1.5%
2.7%
Environmental Variance Ratio: 5
No Effect Matrix: 84%
1.4%
13.6%
0.1%
0.3%
0.4%
0.6%
Correlation
1.00
0.75
0.50
0.25
0.00
−0.25
−0.50
−0.75
−1.00
female−specific
female x3
female x2
female x1.5
equal
male x1.5
male x2
male x3
male−specific
Magnitude

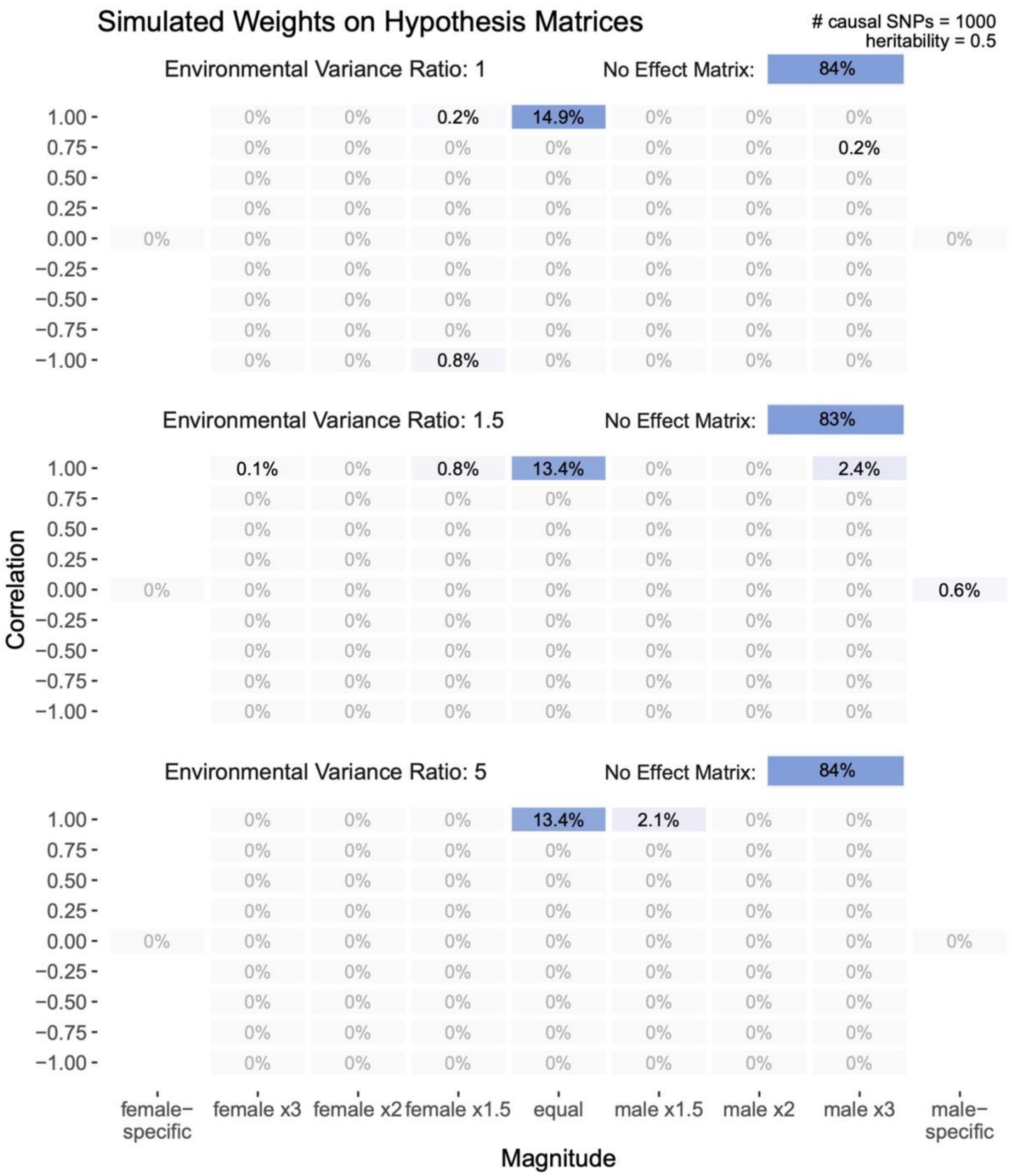
Simulated Weights on Hypothesis Matrices
### causal SNPs = 1000
heritability = 0.5
Environmental Variance Ratio: 1
No Effect Matrix: 84%
0.2%
14.9%
0.2%
0.8%
Environmental Variance Ratio: 1.5
No Effect Matrix: 83%
0.1%
0.8%
13.4%
2.4%
0.6%
Environmental Variance Ratio: 5
No Effect Matrix: 84%
13.4%
2.1%
0%
1.00
0.75
0.50
0.25
0.00
−0.25
−0.50
−0.75
−1.00
Correlation
female−specific
female x3
female x2
female x1.5
equal
male x1.5
male x2
male x3
male−specific
Magnitude

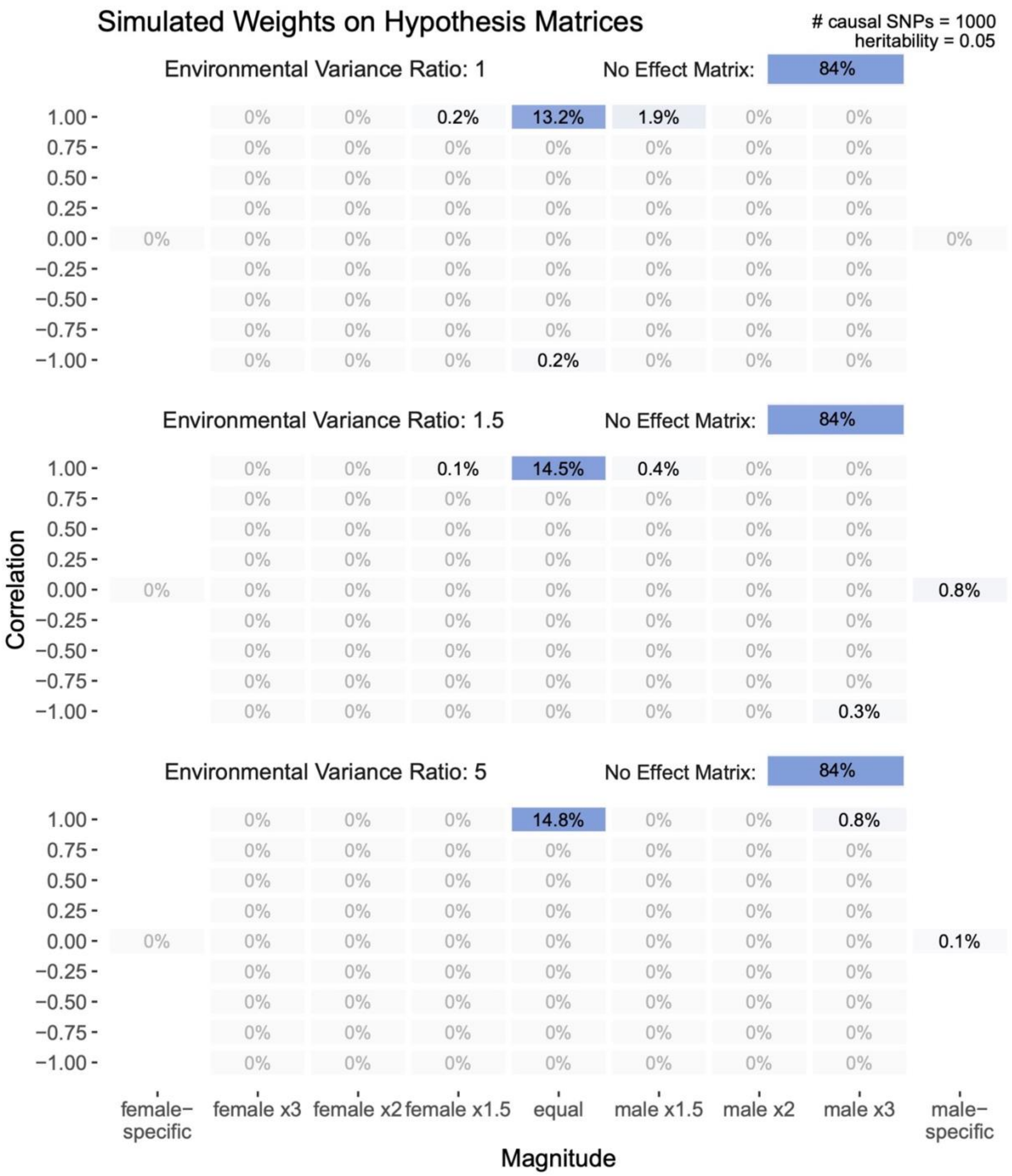

Simulated Weights on Hypothesis Matrices
### causal SNPs = 1000
heritability = 0.05
Environmental Variance Ratio: 1
No Effect Matrix: 84%
Environmental Variance Ratio: 1.5
No Effect Matrix: 84%
Environmental Variance Ratio: 5
No Effect Matrix: 84%
Correlation
Magnitude
female−specific
female x3
female x2
female x1.5
equal
male x1.5
male x2
male x3
male−specific

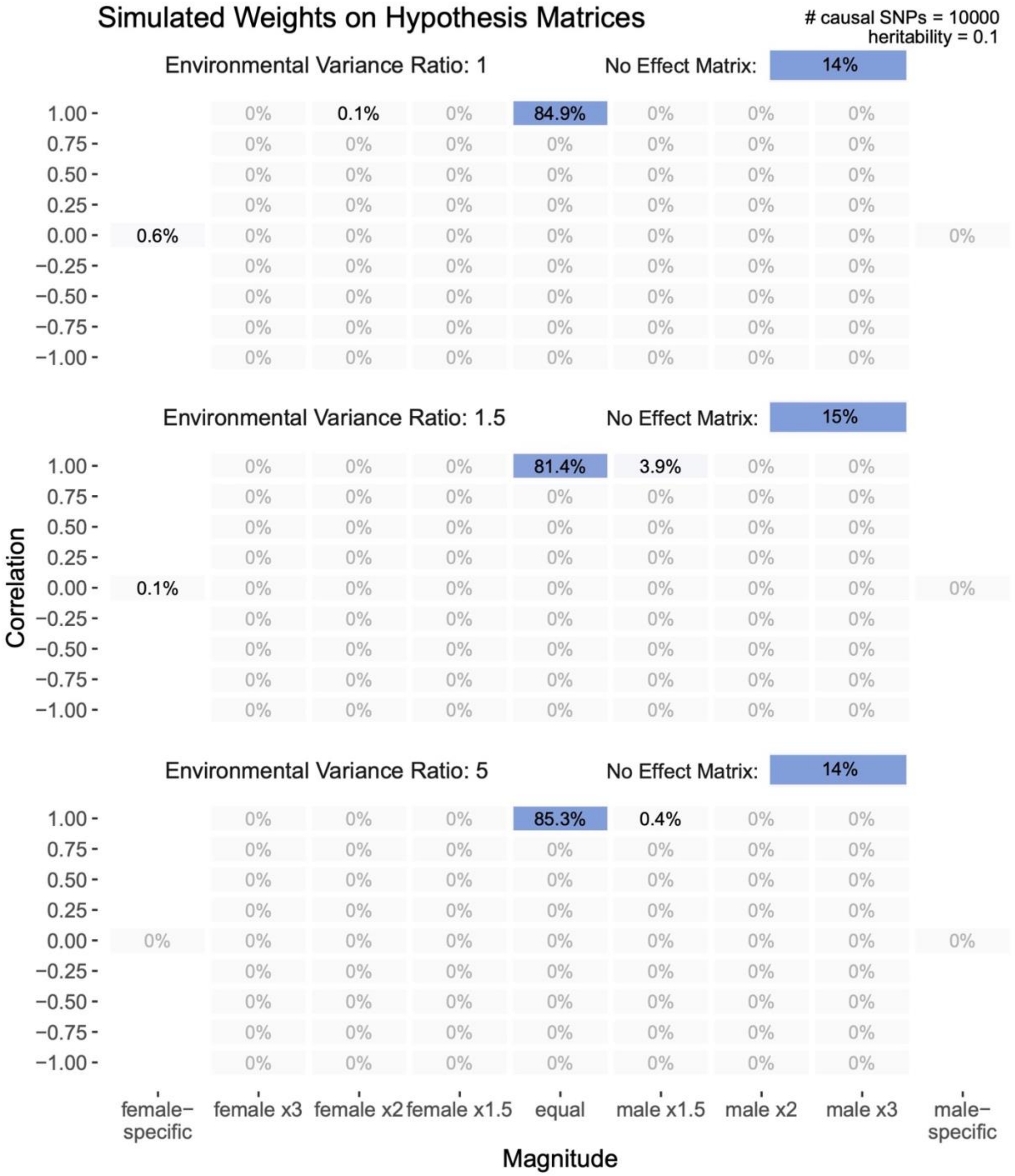

Simulated Weights on Hypothesis Matrices
### causal SNPs = 10000
heritability = 0.1
Environmental Variance Ratio: 1
No Effect Matrix: 14%
Environmental Variance Ratio: 1.5
No Effect Matrix: 15%
Environmental Variance Ratio: 5
No Effect Matrix: 14%
Correlation
Magnitude
female−specific
female x3
female x2
female x1.5
equal
male x1.5
male x2
male x3
male−specific

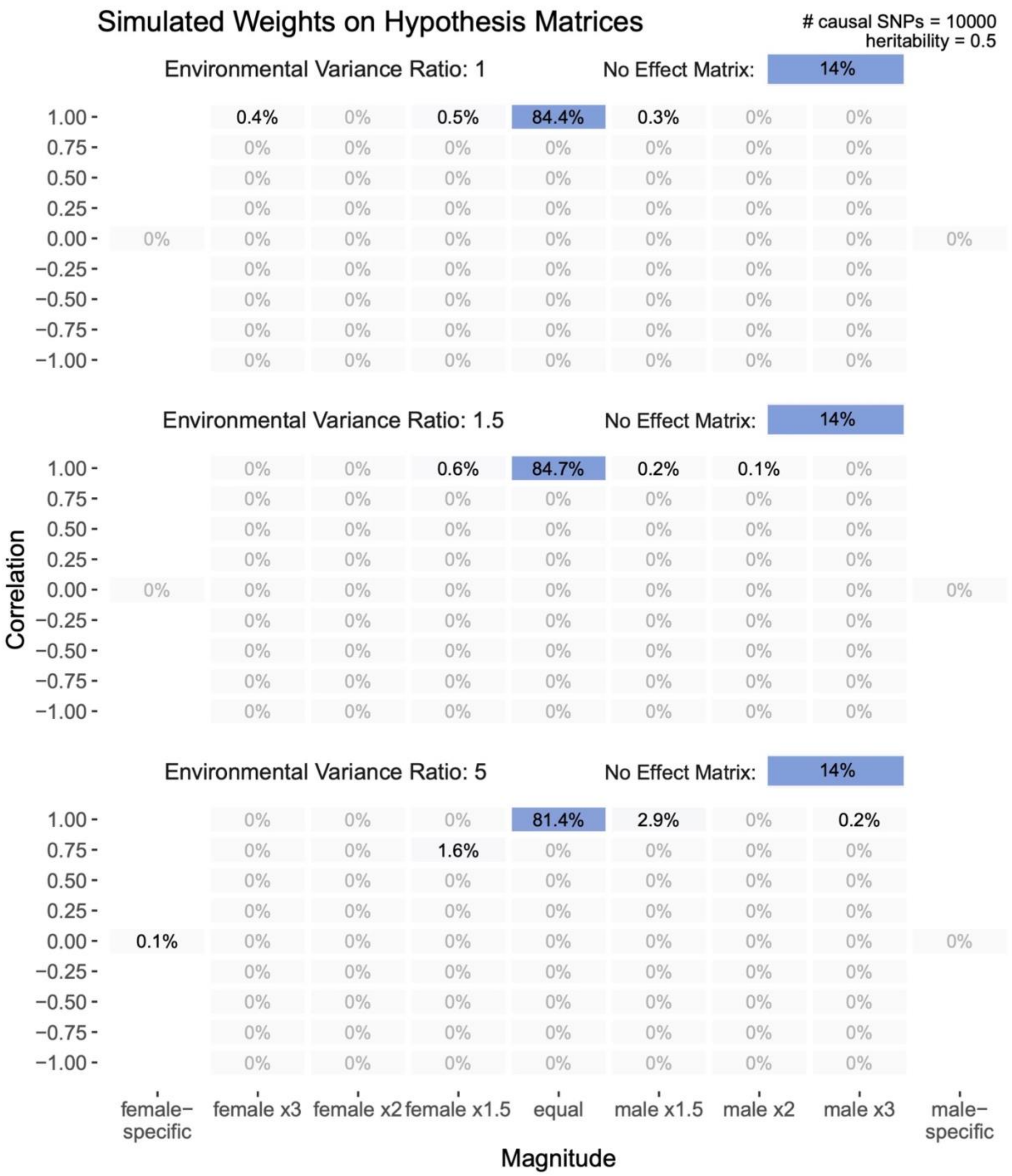
Simulated Weights on Hypothesis Matrices
### causal SNPs = 10000
heritability = 0.5
Environmental Variance Ratio: 1
No Effect Matrix: 14%
0.4%
0.5%
84.4%
0.3%
Environmental Variance Ratio: 1.5
No Effect Matrix: 14%
0.6%
84.7%
0.2%
0.1%
Environmental Variance Ratio: 5
No Effect Matrix: 14%
81.4%
2.9%
0.2%
1.6%
0.1%
Correlation
1.00
0.75
0.50
0.25
0.00
−0.25
−0.50
−0.75
−1.00
female−specific
female x3
female x2
female x1.5
equal
male x1.5
male x2
male x3
male−specific
Magnitude

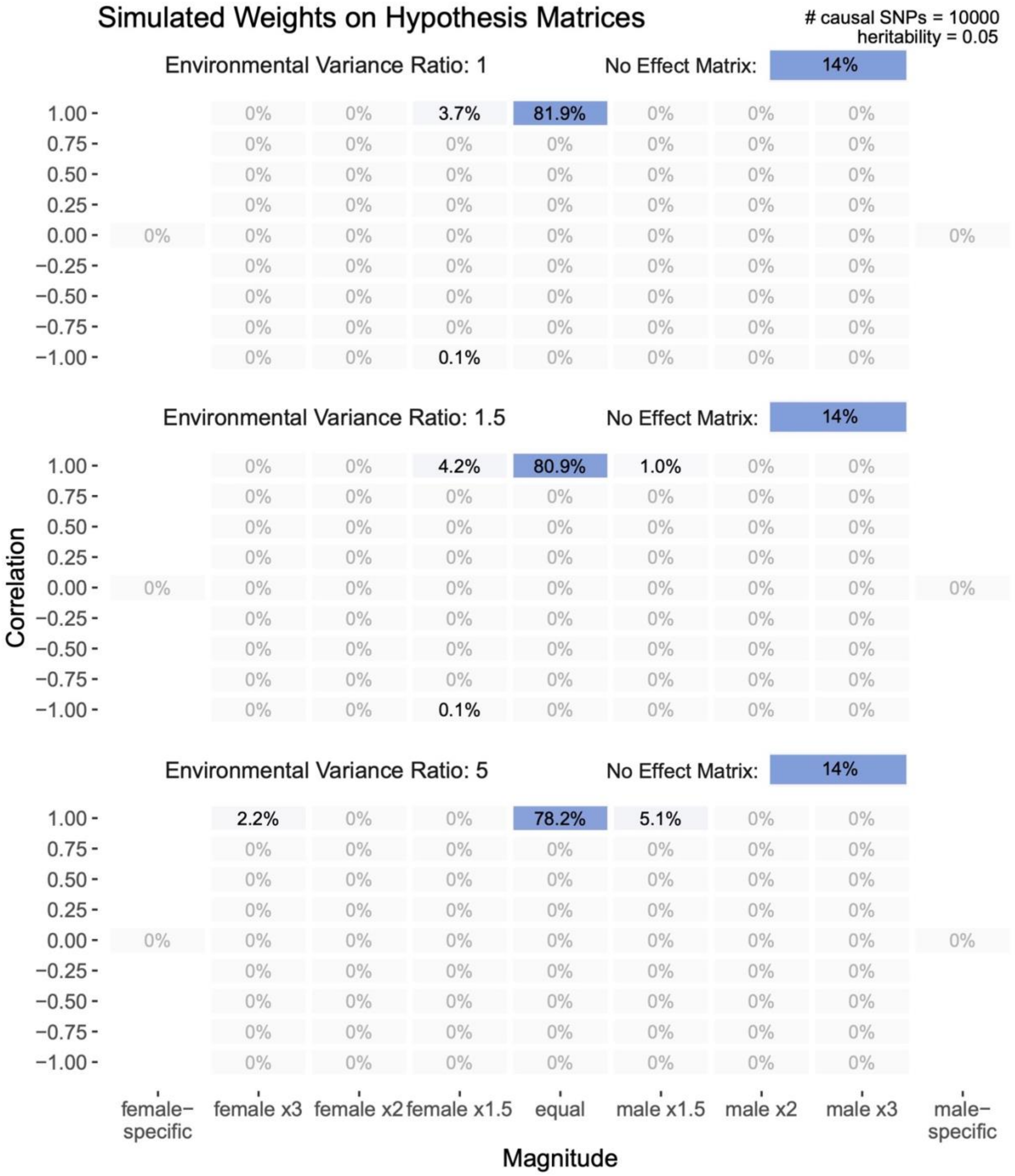

**Figure S7: Simulation of covariance structure, related to Figure 3**

We compared the distribution of mixture weights estimated in *mash* when inputting effect estimates and standard errors are sampled from a pre-specified variance-covariance matrix. The matrices and associated proportions sampled from a particular matrix is shown on the left.

We set the female to male environmental variance ratio as 1.2:1 and the heritability to 0.5. **(A)** We simulate a null model in which all effect estimates are drawn from a matrix representing equal effects between males and females. **(B)** 86% of the true effects have equal effects in males and females and 14% are sampled from a relationship in which effects in females are perfectly correlated but have twice the variance of that in males. **(C)** 86% of the true effects have equal effects in males and females and 14% are sampled from a matrix corresponding to female-private effects.

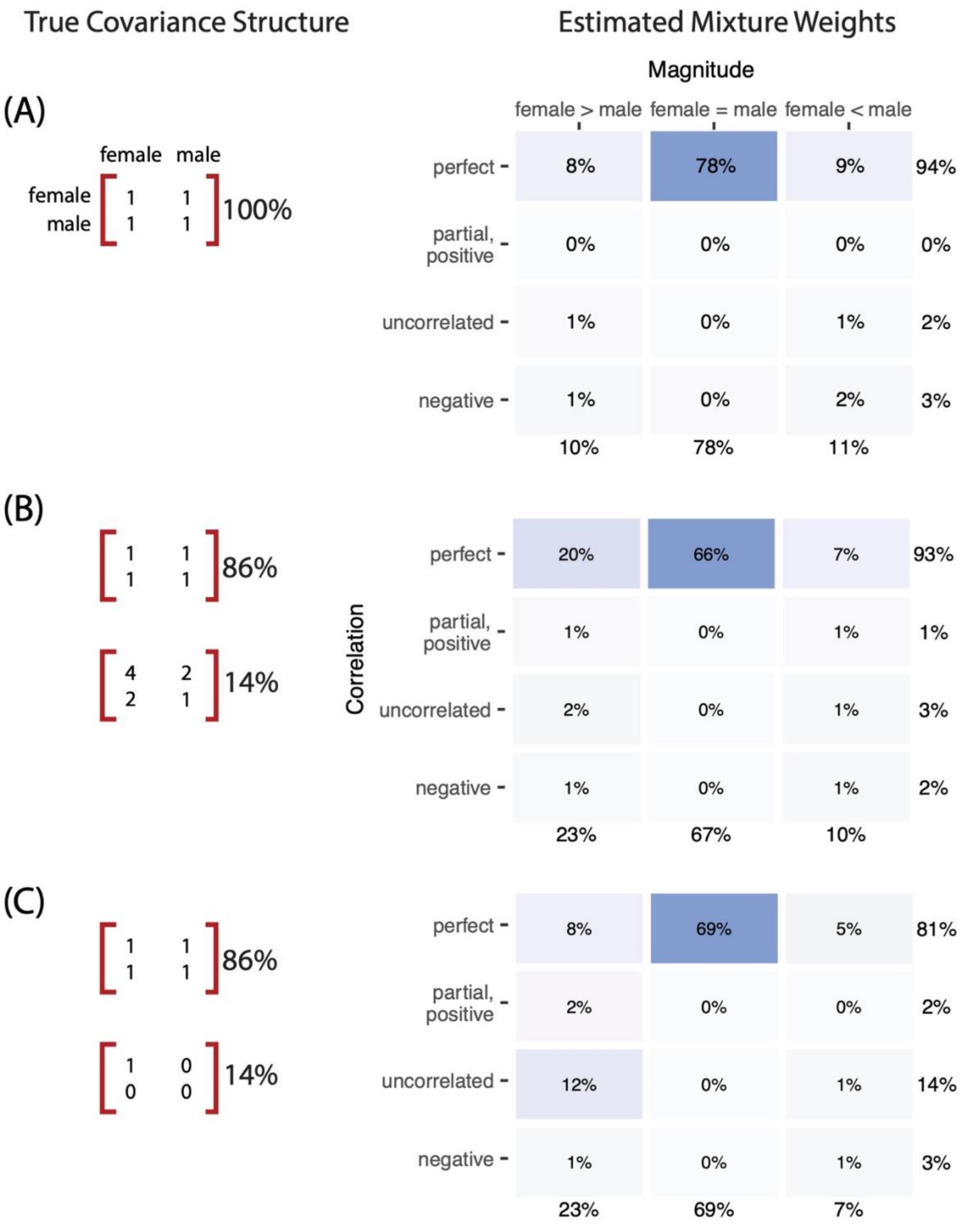
True Covariance Structure
Estimated Mixture Weights
Magnitude
female > male
female = male
female < male
Correlation
perfect
partial, positive
uncorrelated
negative
(A)
female male
female
male
1 1
1 1
100%
8% 78% 9% 94%
0% 0% 0% 0%
1% 0% 1% 2%
1% 0% 2% 3%
10% 78% 11%
(B)
1 1
1 1
86%
4 2
2 1
14%
20% 66% 7% 93%
1% 0% 1% 1%
2% 0% 1% 3%
1% 0% 1% 2%
23% 67% 10%
(C)
1 1
1 1
86%
1 0
0 0
14%
8% 69% 5% 81%
2% 0% 0% 2%
12% 0% 1% 14%
1% 0% 1% 3%
23% 69% 7%

**Figure S8: Proportion of non-trivial weights on hypothesis matrices, related to Figure 3 and Figure 4**

We examine the proportion of weight on non-trivial hypothesis matrices across 27 complex traits. Here, we define non-trivial as matrices with correlation < 1 or female ≠ male magnitude. The heritability ratio represented by point size is estimated by taking the larger of the two sex-specific heritabilities and dividing it by the smaller. We refer to the 1:1 line to see if more traits are represented more by unequal magnitude or imperfect correlation.

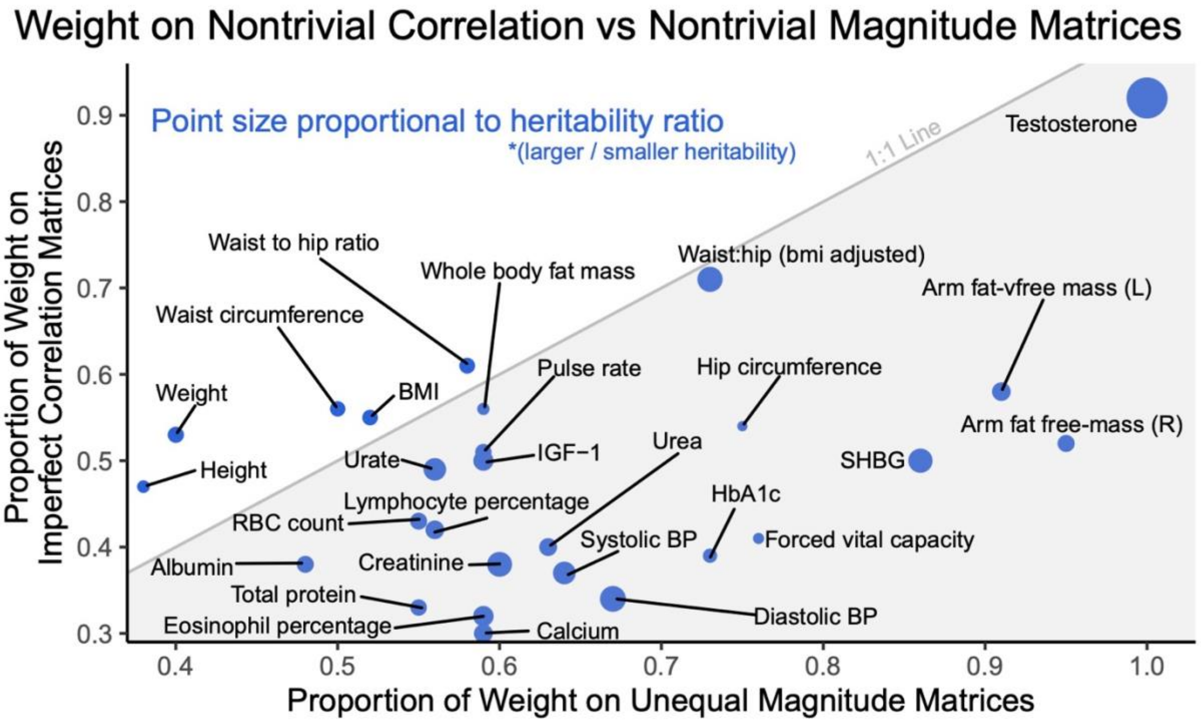

**Figure S9: Across traits, phenotypic variance sex ratio correlates with the phenotypic mean sex ratio, related to Figure 4**

The solid gray line shows a linear fit, excluding two outliers: testosterone and waist:hip ratio adjusted for BMI.

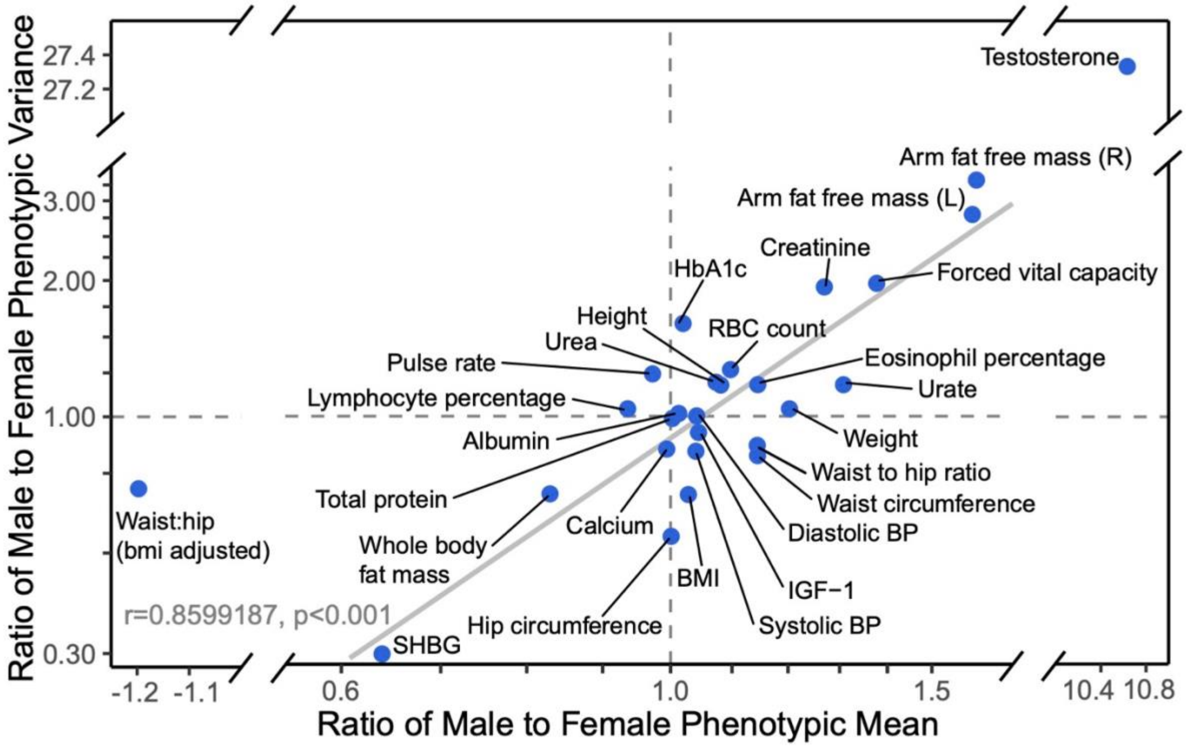

**Figure S10: Process of generating posterior estimates from *mash*, related to Figure 3**

Posterior mean and local false sign rate (lfsr) estimates are generated using *mash* from sex-specific summary statistics. We fit the model on the average of 100 mixture weights. The process is described in the **Text S7** section.

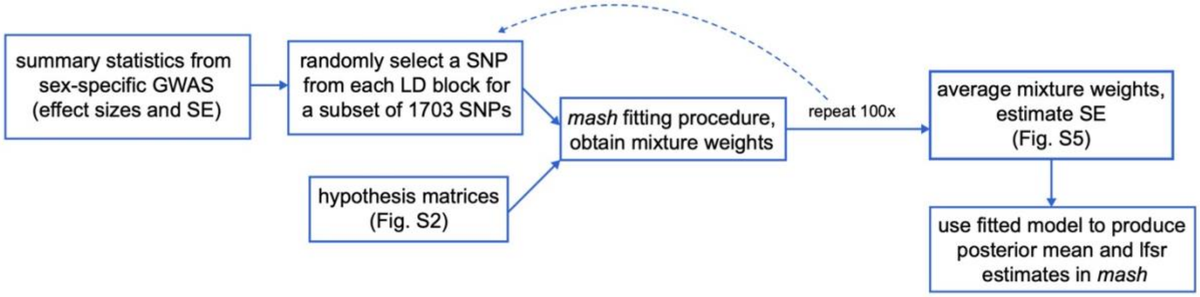

**Figure S11: Process of comparing models for complex phenotype prediction, related to Figure 3**

The process is described in the **Text S8**. **(A)** The sample test set is obtained by sampling 25k males and 25k females. We then create the training set, by removing the test set from the original list of sample IDs. **(B)** After obtaining summary statistics from GWAS, we perform clumping to adjust for LD. Covariance-aware effect estimates are acquired by inputting the sex-specific summary statistics through *mash* to generate posterior mean and lfsr estimates **Fig. S10**. lfsr estimates are converted into pseudo p-values by matching to original p-values after ordering both from smallest to largest. We also conduct PGS analysis over four p-value thresholds on each of the five models, thus obtaining 20 PGS. **(C)** Prediction accuracy is measured based on $R^2$, from the linear regression of phenotype value on covariates and the polygenic scores. The incremental $R^2$ is also estimated by subtracting $R^2$ from the null model $R^2$, based on the regression of the phenotype value on just the covariates.

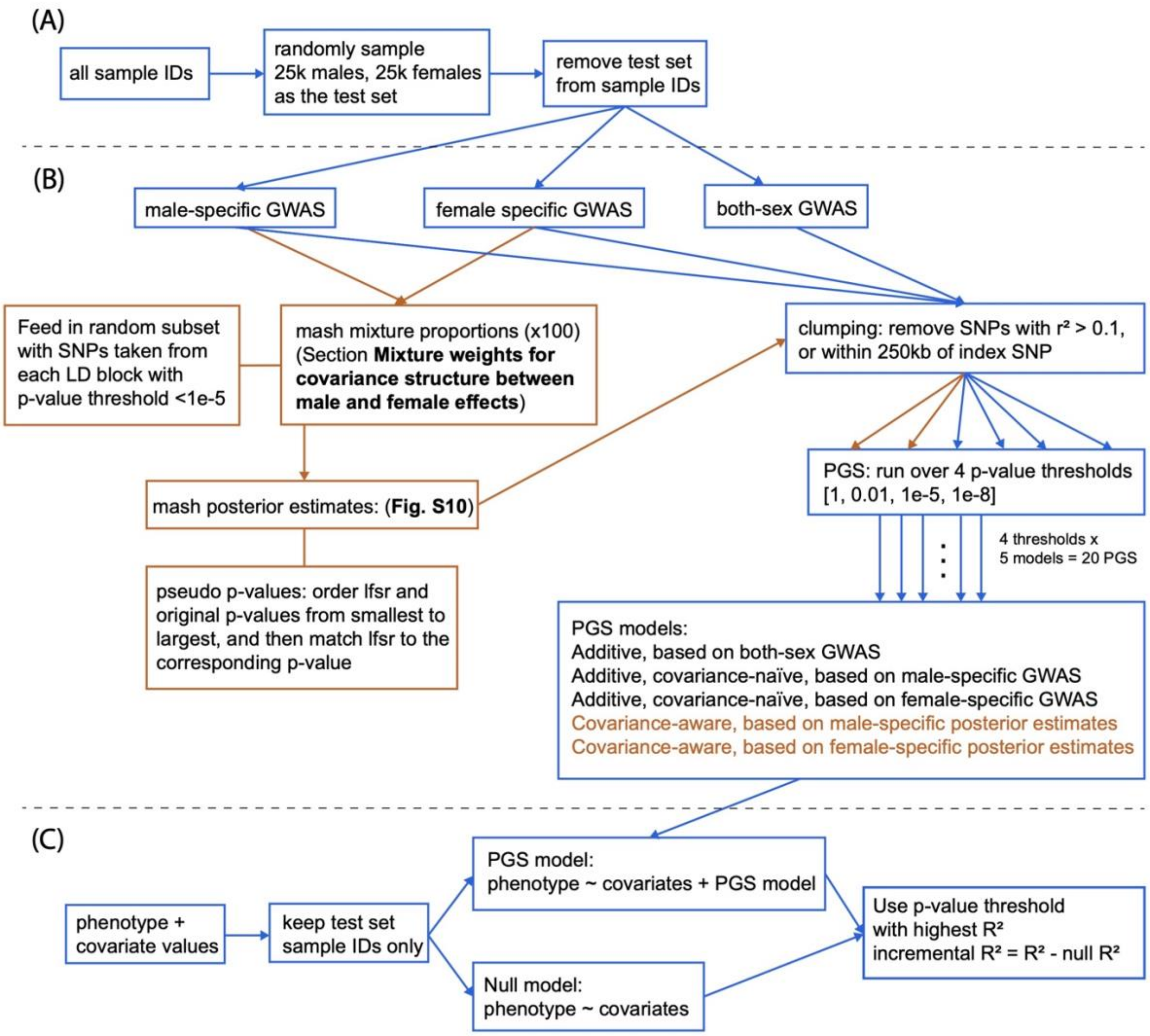

**Figure S12: Evaluating evidence for systematic amplification, related to Figure 2**

We regressed the male (green) and female (orange) trait values to polygenic scores estimated in an independent sample of males (top) and females (bottom) separately. Each line corresponds to a separate regression of trait values in one sex to polygenic scores estimated in one sex. Each point represents the mean value in one decile of the polygenic score. The fitted line and associated R2 are estimated from the raw, non-binned data.

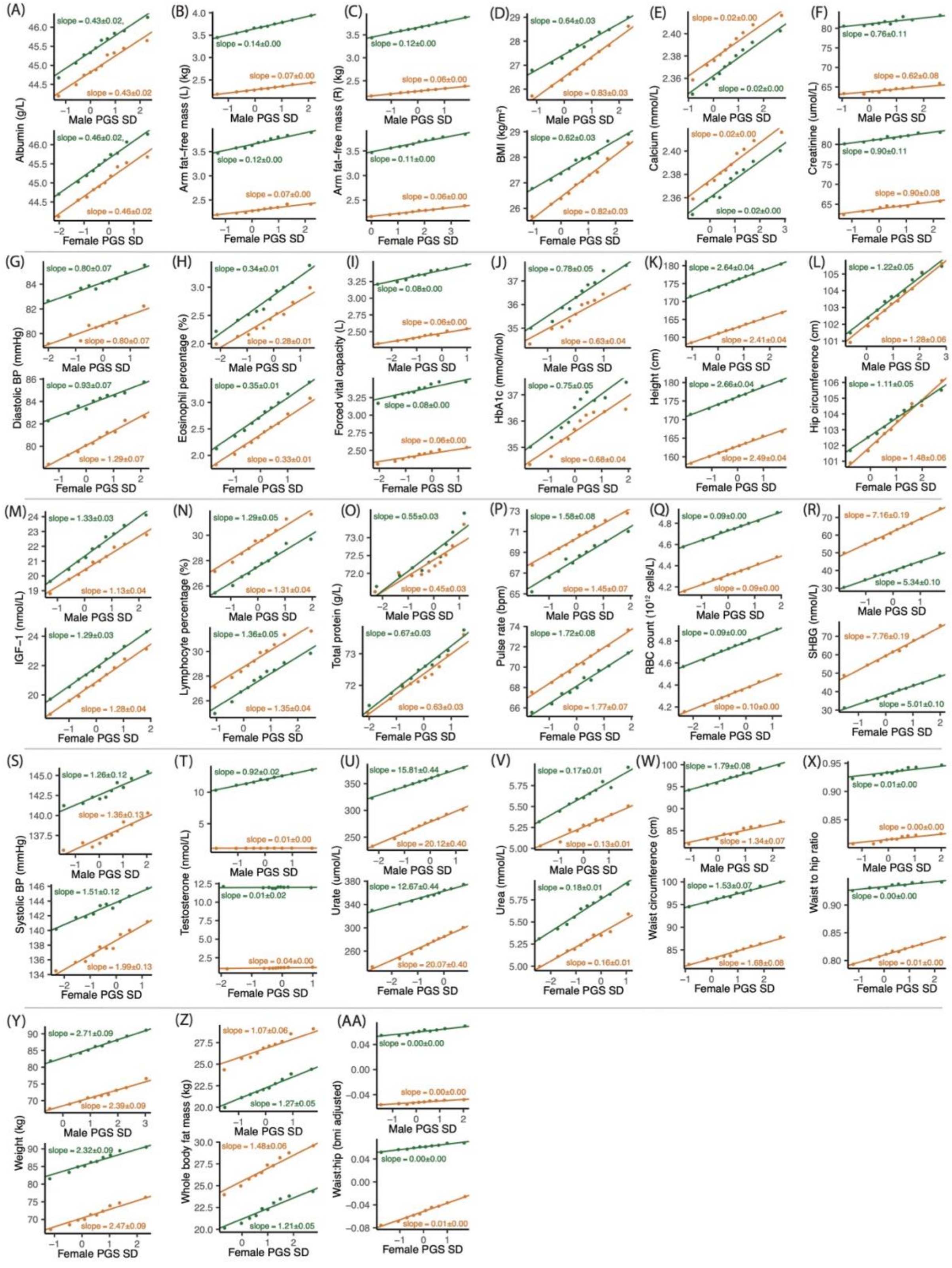
(A)
slope = 0.43±0.02,
slope = 0.43±0.02
slope = 0.46±0.02,
slope = 0.46±0.02
Albumin (g/L)
(B)
slope = 0.14±0.00
slope = 0.07±0.00
slope = 0.12±0.00
slope = 0.07±0.00
Arm fat–free mass (L) (kg)
(C)
slope = 0.12±0.00
slope = 0.06±0.00
slope = 0.11±0.00
slope = 0.06±0.00
Arm fat–free mass (R) (kg)
(D)
slope = 0.64±0.03
slope = 0.83±0.03
slope = 0.62±0.03
slope = 0.82±0.03
BMI (kg/m²)
(E)
slope = 0.02±0.00
slope = 0.02±0.00
slope = 0.02±0.00
slope = 0.02±0.00
Calcium (mmol/L)
(F)
slope = 0.76±0.11
slope = 0.62±0.08
slope = 0.90±0.11
slope = 0.90±0.08
Creatinine (umol/L)
(G)
slope = 0.80±0.07
slope = 0.80±0.07
slope = 0.93±0.07
slope = 1.29±0.07
Diastolic BP (mmHg)
(H)
slope = 0.34±0.01
slope = 0.28±0.01
slope = 0.35±0.01
slope = 0.33±0.01
Eosinophil percentage (%)
(I)
slope = 0.08±0.00
slope = 0.06±0.00
slope = 0.08±0.00
slope = 0.06±0.00
Forced vital capacity (L)
(J)
slope = 0.78±0.05
slope = 0.63±0.04
slope = 0.75±0.05
slope = 0.68±0.04
HbA1c (mmol/mol)
(K)
slope = 2.64±0.04
slope = 2.41±0.04
slope = 2.66±0.04
slope = 2.49±0.04
Height (cm)
(L)
slope = 1.22±0.05
slope = 1.28±0.06
slope = 1.11±0.05
slope = 1.48±0.06
Hip circumference (cm)
(M)
slope = 1.33±0.03
slope = 1.13±0.04
slope = 1.29±0.03
slope = 1.28±0.04
IGF−1 (nmol/L)
(N)
slope = 1.29±0.05
slope = 1.31±0.04
slope = 1.36±0.05
slope = 1.35±0.04
Lymphocyte percentage (%)
(O)
slope = 0.55±0.03
slope = 0.45±0.03
slope = 0.67±0.03
slope = 0.63±0.03
Total protein (g/L)
(P)
slope = 1.58±0.08
slope = 1.45±0.07
slope = 1.72±0.08
slope = 1.77±0.07
Pulse rate (bpm)
(Q)
slope = 0.09±0.00
slope = 0.09±0.00
slope = 0.09±0.00
slope = 0.10±0.00
RBC count (10^12 cells/L)
(R)
slope = 7.16±0.19
slope = 5.34±0.10
slope = 7.76±0.19
slope = 5.01±0.10
SHBG (nmol/L)
(S)
slope = 1.26±0.12
slope = 1.36±0.13
slope = 1.51±0.12
slope = 1.99±0.13
Systolic BP (mmHg)
(T)
slope = 0.92±0.02
slope = 0.01±0.00
slope = 0.01±0.02
slope = 0.04±0.00
Testosterone (nmol/L)
(U)
slope = 15.81±0.44
slope = 20.12±0.40
slope = 12.67±0.44
slope = 20.07±0.40
Urate (umol/L)
(V)
slope = 0.17±0.01
slope = 0.13±0.01
slope = 0.18±0.01
slope = 0.16±0.01
Urea (mmol/L)
(W)
slope = 1.79±0.08
slope = 1.34±0.07
slope = 1.53±0.07
slope = 1.68±0.08
Waist circumference (cm)
(X)
slope = 0.01±0.00
slope = 0.00±0.00
slope = 0.00±0.00
slope = 0.01±0.00
Waist to hip ratio
(Y)
slope = 2.71±0.09
slope = 2.39±0.09
slope = 2.32±0.09
slope = 2.47±0.09
Weight (kg)
(Z)
slope = 1.07±0.06
slope = 1.27±0.05
slope = 1.48±0.06
slope = 1.21±0.05
Whole body fat mass (kg)
(AA)
slope = 0.00±0.00
slope = 0.00±0.00
slope = 0.00±0.00
slope = 0.01±0.00
Waist:hip (bmi adjusted)
Male PGS SD
Female PGS SD

**Figure S13: The predictive utility of G-by-Sex aware polygenic scores, related to Figure 3**

We compared the incremental squared correlation (incremental $R^2$) of the phenotypic values and a predictor for three predictors. Incremental $R^2$ was obtained by taking the difference in $R^2$ between a prediction with covariates and the polygenic score and a prediction with only covariates. For each of the three models, we chose the highest incremental $R^2$ across four p-value thresholds (**Text S8**). The additive (yellow), and sex-specific covariance-naïve (blue), and sex-specific covariance aware (red) models were calculated from effect estimates generated in GWAS, with the latter two in sex-stratified samples. The sex-specific, covariance aware model (red) refers to a polygenic score constructed using *mash* posterior effect estimates, which considers the covariance in genetic effects between the sexes. Sex-specific models contained a median sample size of around 123K males and 147K females, whereas the additive model retained the full training set of around 270K individuals. Incremental $R^2$ values and ±3 SE-wide error bars were computed using 20-fold cross-validation.

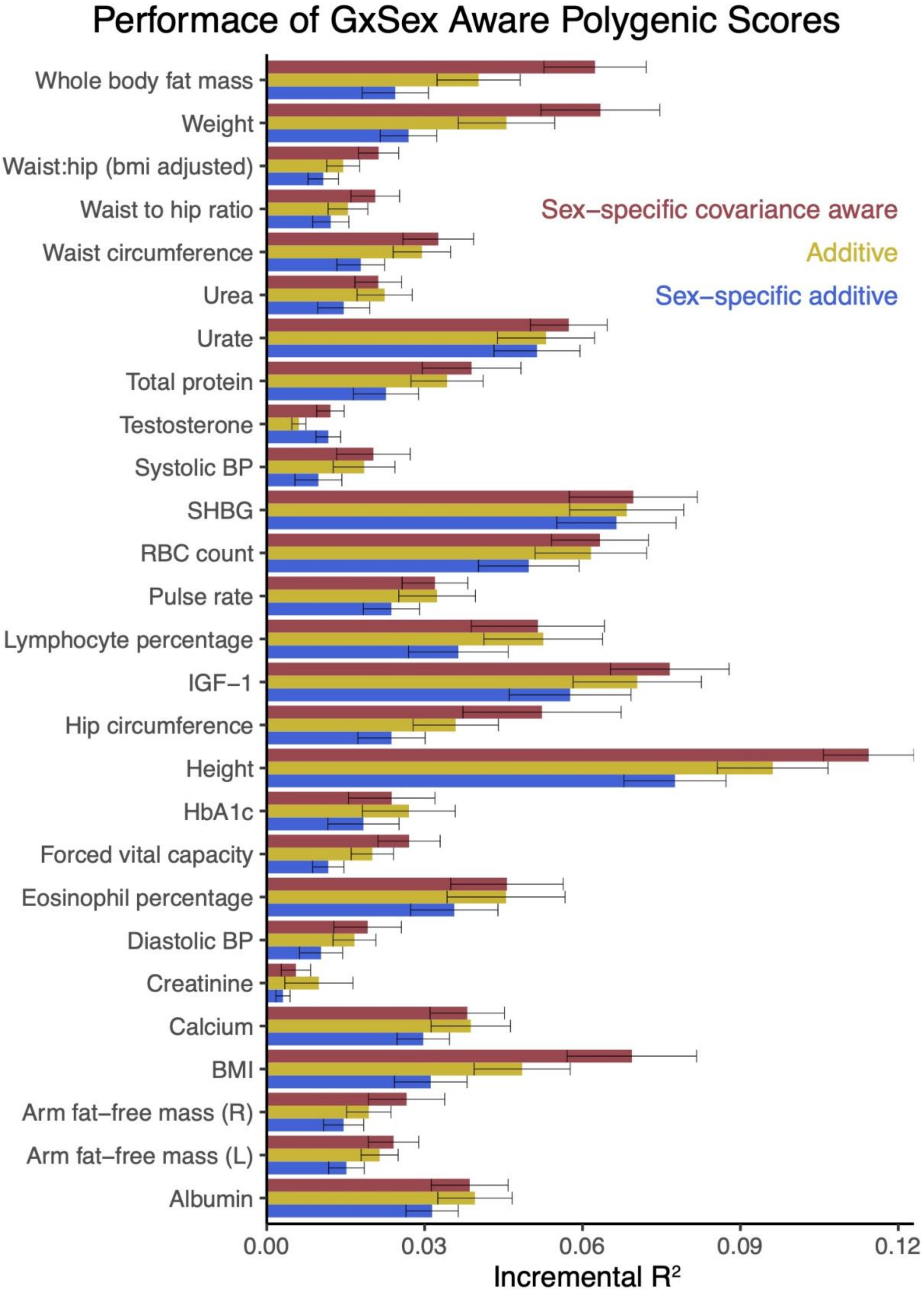
Performace of GxSex Aware Polygenic Scores
Whole body fat mass
Weight
Waist:hip (bmi adjusted)
Waist to hip ratio
Waist circumference
Urea
Urate
Total protein
Testosterone
Systolic BP
SHBG
RBC count
Pulse rate
Lymphocyte percentage
IGF−1
Hip circumference
Height
HbA1c
Forced vital capacity
Eosinophil percentage
Diastolic BP
Creatinine
Calcium
BMI
Arm fat−free mass (R)
Arm fat−free mass (L)
Albumin
Sex−specific covariance aware
Additive
Sex−specific additive
0.00
0.03
0.06
0.09
0.12
Incremental R²

**Figure S14: Genetic effect and testosterone levels – using additive model PGS, related to Figure 5**

The relationship between genetic effect and testosterone level bins, separated by sex, is depicted for all traits. Genetic effect is estimated from the slope of the regression of phenotypic values and polygenic scores from the additive model, multiplied by the polygenic score standard deviation (**Fig. S13**). The hollow data point represents a testosterone bin with overlapping range between males and females.

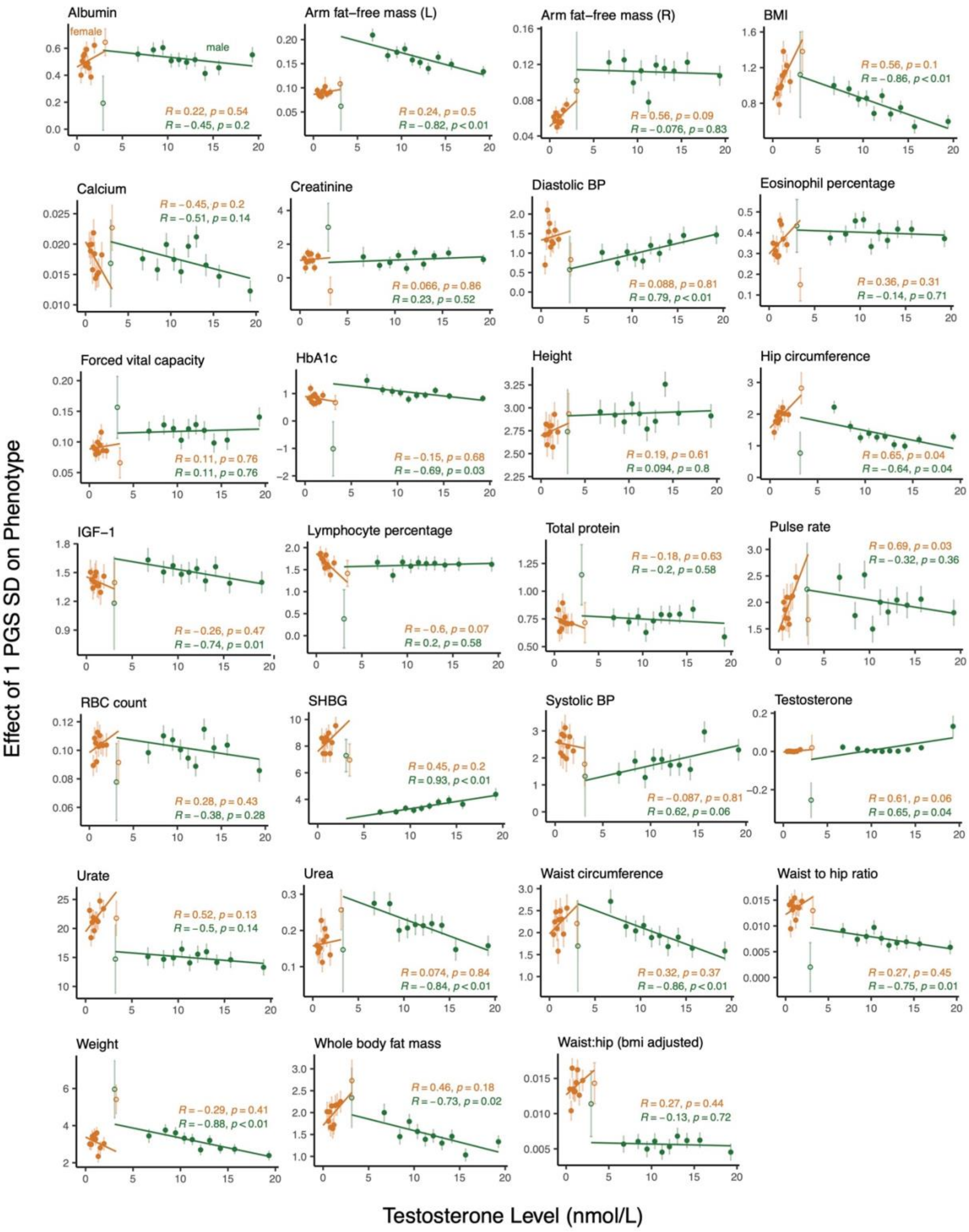
Effect of 1 PGS SD on Phenotype
Testosterone Level (nmol/L)
female
male
Albumin
R = 0.22, p = 0.54
R = −0.45, p = 0.2
Arm fat−free mass (L)
R = 0.24, p = 0.5
R = −0.82, p < 0.01
Arm fat−free mass (R)
R = 0.56, p = 0.09
R = −0.076, p = 0.83
BMI
R = 0.56, p = 0.1
R = −0.86, p < 0.01
Calcium
R = −0.45, p = 0.2
R = −0.51, p = 0.14
Creatinine
R = 0.066, p = 0.86
R = 0.23, p = 0.52
Diastolic BP
R = 0.088, p = 0.81
R = 0.79, p < 0.01
Eosinophil percentage
R = 0.36, p = 0.31
R = −0.14, p = 0.71
Forced vital capacity
R = 0.11, p = 0.76
R = 0.11, p = 0.76
HbA1c
R = −0.15, p = 0.68
R = −0.69, p = 0.03
Height
R = 0.19, p = 0.61
R = 0.094, p = 0.8
Hip circumference
R = 0.65, p = 0.04
R = −0.64, p = 0.04
IGF−1
R = −0.26, p = 0.47
R = −0.74, p = 0.01
Lymphocyte percentage
R = −0.6, p = 0.07
R = 0.2, p = 0.58
Total protein
R = −0.18, p = 0.63
R = −0.2, p = 0.58
Pulse rate
R = 0.69, p = 0.03
R = −0.32, p = 0.36
RBC count
R = 0.28, p = 0.43
R = −0.38, p = 0.28
SHBG
R = 0.45, p = 0.2
R = 0.93, p < 0.01
Systolic BP
R = −0.087, p = 0.81
R = 0.62, p = 0.06
Testosterone
R = 0.61, p = 0.06
R = 0.65, p = 0.04
Urate
R = 0.52, p = 0.13
R = −0.5, p = 0.14
Urea
R = 0.074, p = 0.84
R = −0.84, p < 0.01
Waist circumference
R = 0.32, p = 0.37
R = −0.86, p < 0.01
Waist to hip ratio
R = 0.27, p = 0.45
R = −0.75, p = 0.01
Weight
R = −0.29, p = 0.41
R = −0.88, p < 0.01
Whole body fat mass
R = 0.46, p = 0.18
R = −0.73, p = 0.02
Waist:hip (bmi adjusted)
R = 0.27, p = 0.44
R = −0.13, p = 0.72

**Figure S15: Genetic effect and testosterone levels – using sex-specific model PGS, related to Figure 5**

The same type of figures as **Fig. S14** are displayed with the exception of utilizing PGS from the sex-specific, covariance-naïve model for estimation of genetic effect.

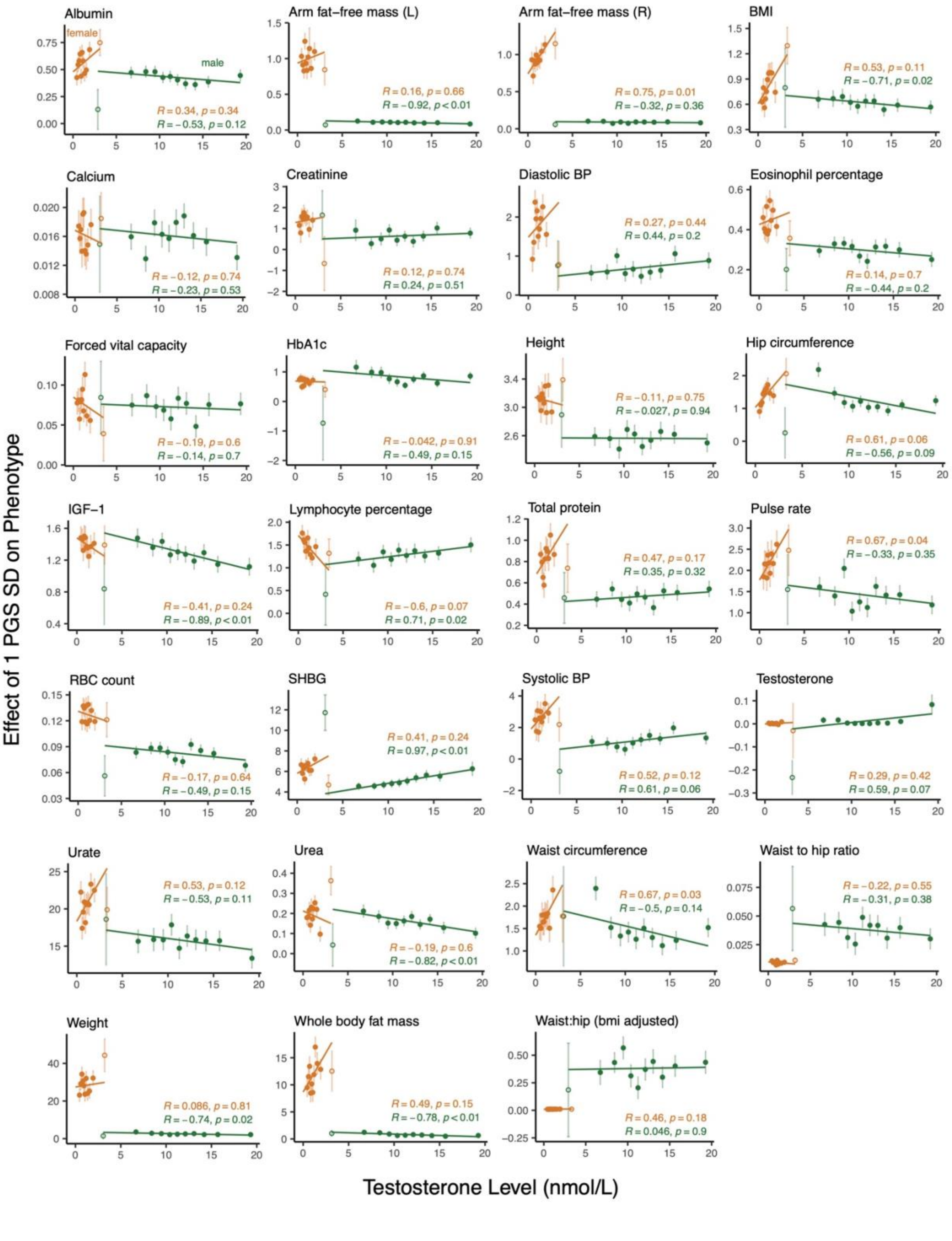
Albumin
female
male
R = 0.34, p = 0.34
R = −0.53, p = 0.12
Arm fat−free mass (L)
R = 0.16, p = 0.66
R = −0.92, p < 0.01
Arm fat−free mass (R)
R = 0.75, p = 0.01
R = −0.32, p = 0.36
BMI
R = 0.53, p = 0.11
R = −0.71, p = 0.02
Calcium
R = −0.12, p = 0.74
R = −0.23, p = 0.53
Creatinine
R = 0.12, p = 0.74
R = 0.24, p = 0.51
Diastolic BP
R = 0.27, p = 0.44
R = 0.44, p = 0.2
Eosinophil percentage
R = 0.14, p = 0.7
R = −0.44, p = 0.2
Forced vital capacity
R = −0.19, p = 0.6
R = −0.14, p = 0.7
HbA1c
R = −0.042, p = 0.91
R = −0.49, p = 0.15
Height
R = −0.11, p = 0.75
R = −0.027, p = 0.94
Hip circumference
R = 0.61, p = 0.06
R = −0.56, p = 0.09
IGF−1
R = −0.41, p = 0.24
R = −0.89, p < 0.01
Lymphocyte percentage
R = −0.6, p = 0.07
R = 0.71, p = 0.02
Total protein
R = 0.47, p = 0.17
R = 0.35, p = 0.32
Pulse rate
R = 0.67, p = 0.04
R = −0.33, p = 0.35
RBC count
R = −0.17, p = 0.64
R = −0.49, p = 0.15
SHBG
R = 0.41, p = 0.24
R = 0.97, p < 0.01
Systolic BP
R = 0.52, p = 0.12
R = 0.61, p = 0.06
Testosterone
R = 0.29, p = 0.42
R = 0.59, p = 0.07
Urate
R = 0.53, p = 0.12
R = −0.53, p = 0.11
Urea
R = −0.19, p = 0.6
R = −0.82, p < 0.01
Waist circumference
R = 0.67, p = 0.03
R = −0.5, p = 0.14
Waist to hip ratio
R = −0.22, p = 0.55
R = −0.31, p = 0.38
Weight
R = 0.086, p = 0.81
R = −0.74, p = 0.02
Whole body fat mass
R = 0.49, p = 0.15
R = −0.78, p < 0.01
Waist:hip (bmi adjusted)
R = 0.46, p = 0.18
R = 0.046, p = 0.9
Effect of 1 PGS SD on Phenotype
Testosterone Level (nmol/L)

**Figure S16: Genetic effect and PGS for testosterone, related to Figure 5**

We used mendelian randomization to assess the causal relationship between genetic effect and testosterone levels, similar to **Fig. S14**. **(A)** The correlation for each sex and all traits between genetic effect and PGS for testosterone across the bins is shown, with traits in descending order by the difference of correlation estimates between the sexes. **(B)** The plots illustrate the relationship between amplification of total genetic effect of each trait and PGS based on sex-specific summary statistics for testosterone. Genetic effect is estimated from the slope of the regression of phenotype values on the PGS based on additive summary statistics in each PGS bin for testosterone, multiplied by the PGS standard deviation. The x-axis takes the mean PGS for testosterone from each bin.

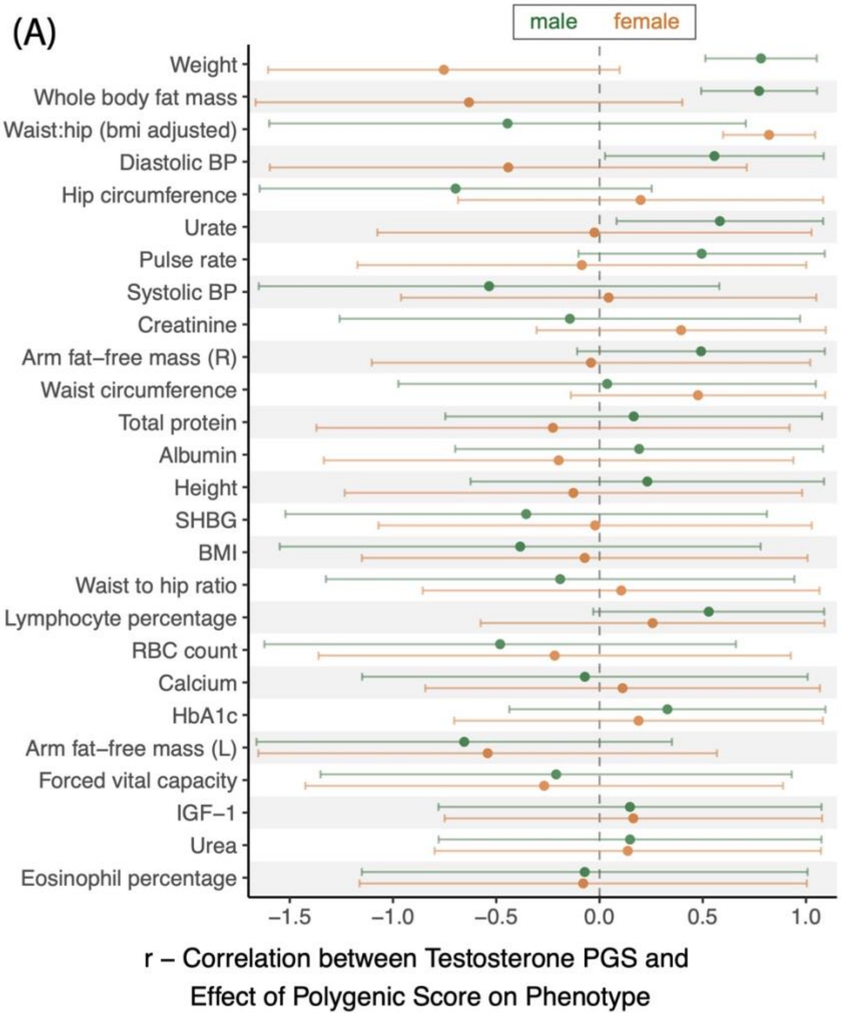

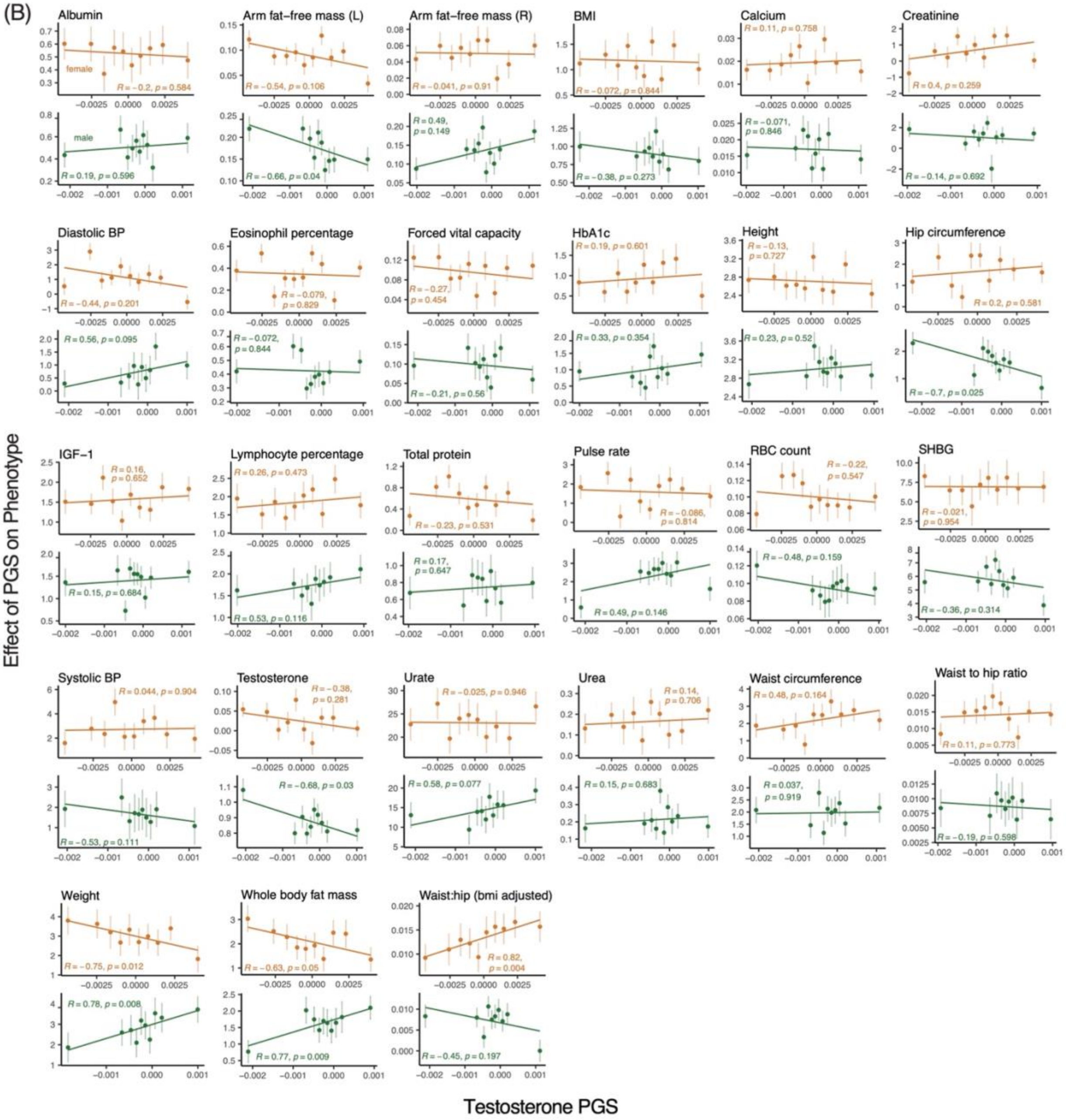

**Figure S17: Genetic effect residualized for age and testosterone levels, related to Figure 5**

In each bin, we examined the relationship between the polygenic effect residualized for mean age and the mean testosterone level. This plot shows the correlation between the two across ten bins.

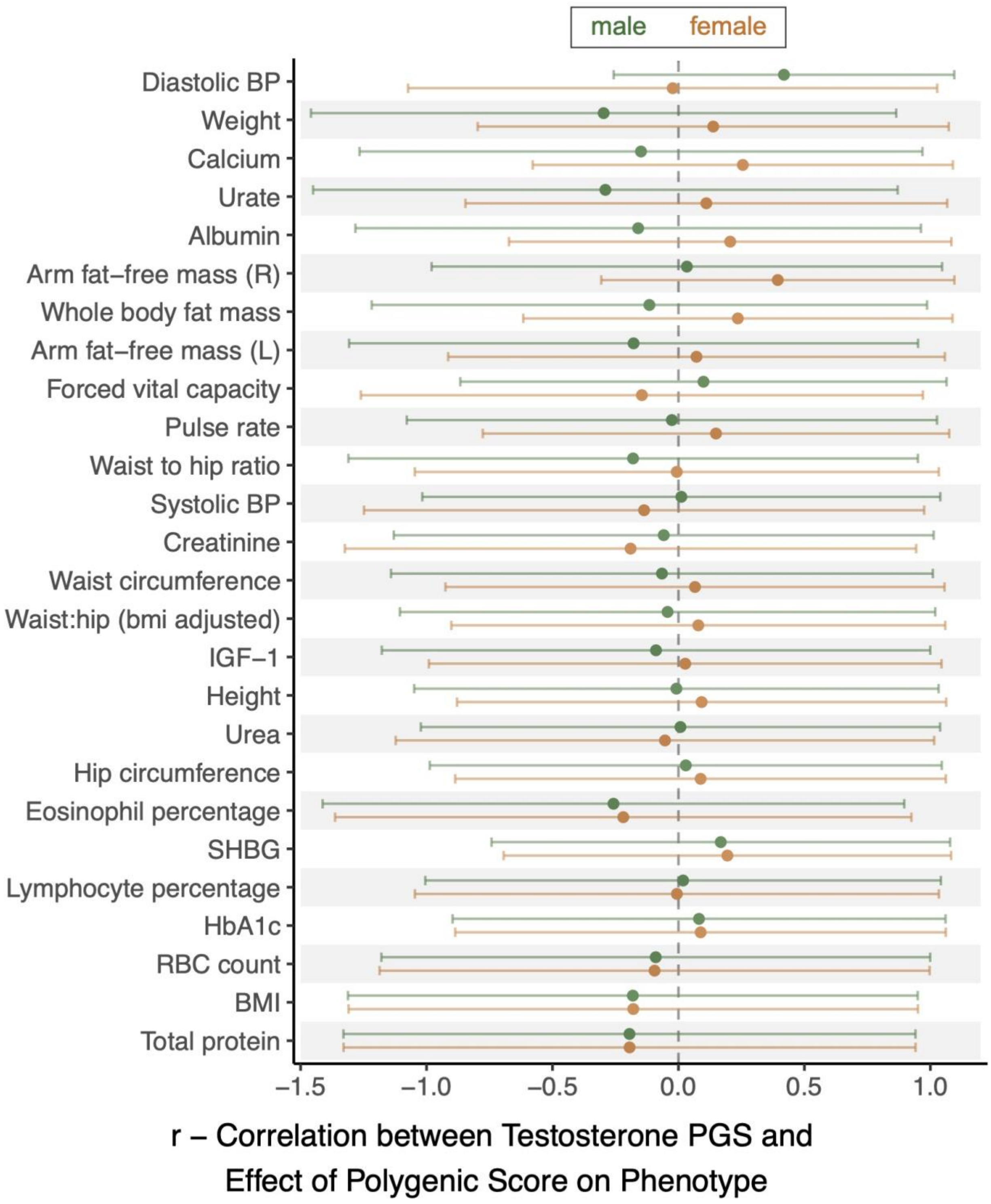
male
female
Diastolic BP
Weight
Calcium
Urate
Albumin
Arm fat−free mass (R)
Whole body fat mass
Arm fat−free mass (L)
Forced vital capacity
Pulse rate
Waist to hip ratio
Systolic BP
Creatinine
Waist circumference
Waist:hip (bmi adjusted)
IGF−1
Height
Urea
Hip circumference
Eosinophil percentage
SHBG
Lymphocyte percentage
HbA1c
RBC count
BMI
Total protein
−1.5
−1.0
−0.5
0.0
0.5
1.0
r – Correlation between Testosterone PGS and
Effect of Polygenic Score on Phenotype

**Figure S18: Example of shared polygenic and environmental amplification in muscle, related to Figure 6**

An example of equal amplification of genetic and environmental effect (**"Are polygenic and environmental effects jointly amplified?"** in main text**)** is depicted in muscle after resistance training. The example follows the model in **Fig. 6A**, in which the genetic effect (genetic regulation) and environmental effect (resistance exercise), both affect a core pathway (involving IGF-1) for muscle. The effect size of the core pathway is then amplified by a modulator such as testosterone, producing the sexual-dimorphism shown in muscle.

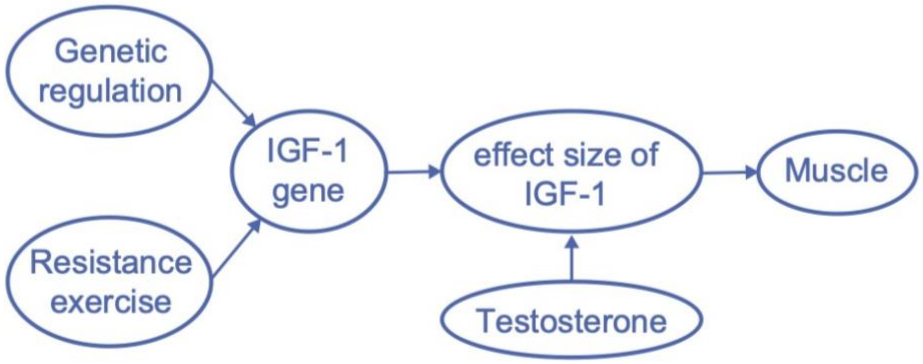

**Figure S19: Competing models for sex differences in trait variance, related to Fig. 6**

We compare two different models for sex differences in trait variance, greater-male and estrus-mediated variability as discussed in Zajitschek et al.[21] and **Text S11**, with our model of pervasive amplification linking genetic and environmental effects ("**Are polygenic and environmental effects jointly amplified?"** in main text**). (A)** We illustrate the difference between the variability between ($V_{between}$) and the variability within ($V_{within}$) using individual distributions. We define the three models according to the relationship between male and female $V_{between}$ and $V_{within.}$ **(B)** The three models are superimposed as different colored lines on **Fig. 6B** to compare the distribution of traits across the models. Traits in blue, green, and yellow are consistent (within 90% CI) with the model of pervasive amplification, greater-male variability, and estrus-mediated variability alone, respectively. Traits in grey are consistent with more than one model. Traits in black are inconsistent with either model.

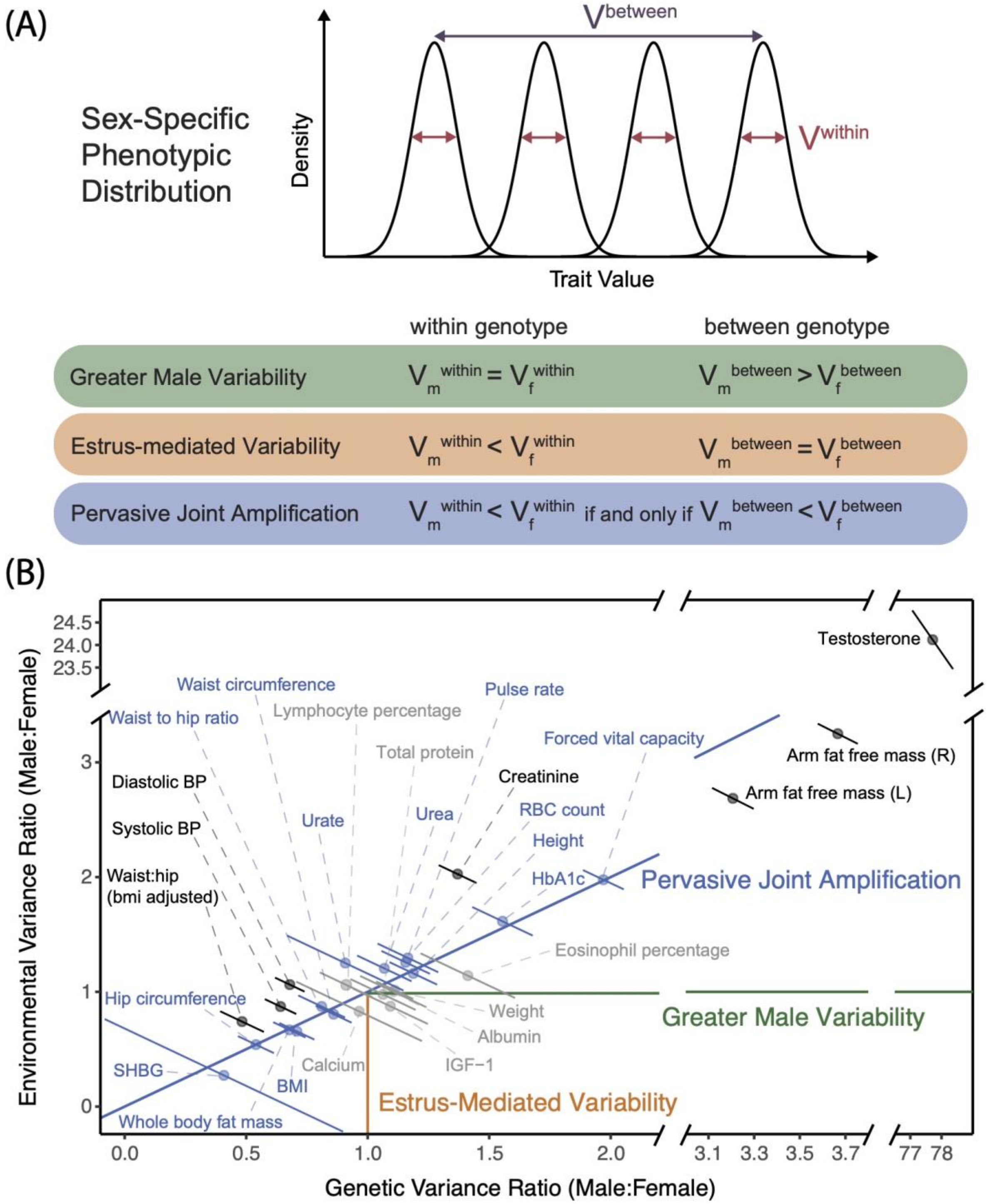
(A)
Sex-Specific Phenotypic Distribution
Density
Trait Value
V between
V within
within genotype
between genotype
Greater Male Variability
$V_m^{within} = V_f^{within}$
$V_m^{between} > V_f^{between}$
Estrus-mediated Variability
$V_m^{within} < V_f^{within}$
$V_m^{between} = V_f^{between}$
Pervasive Joint Amplification
$V_m^{within} < V_f^{within}$ if and only if $V_m^{between} < V_f^{between}$
(B)
Environmental Variance Ratio (Male:Female)
Genetic Variance Ratio (Male:Female)
Testosterone
Waist circumference
Lymphocyte percentage
Pulse rate
Waist to hip ratio
Total protein
Forced vital capacity
Arm fat free mass (R)
Diastolic BP
Creatinine
Arm fat free mass (L)
Systolic BP
Urate
Urea
RBC count
Height
Waist:hip (bmi adjusted)
HbA1c
Pervasive Joint Amplification
Eosinophil percentage
Hip circumference
Weight
Greater Male Variability
Albumin
SHBG
Calcium
IGF-1
BMI
Whole body fat mass
Estrus-Mediated Variability

**Figure S20: Z-scores for strength of sexually-antagonistic selection, related to Figure 7**

Shown are Z-scores for the strength of sexually-antagonistic selection, *A* in **Eq 1**, computed for various gnomAD subsamples (categorized by ancestry group). Different panels show results obtained with different p-value thresholds on marginal significance of sex-specific association in the GWAS (in either sex), including $10^{-3}$ **(A)**, $10^{-5}$ **(B)**, and $10^{-8}$ **(C)**. Bars show 90% resampling confidence intervals.

(A)

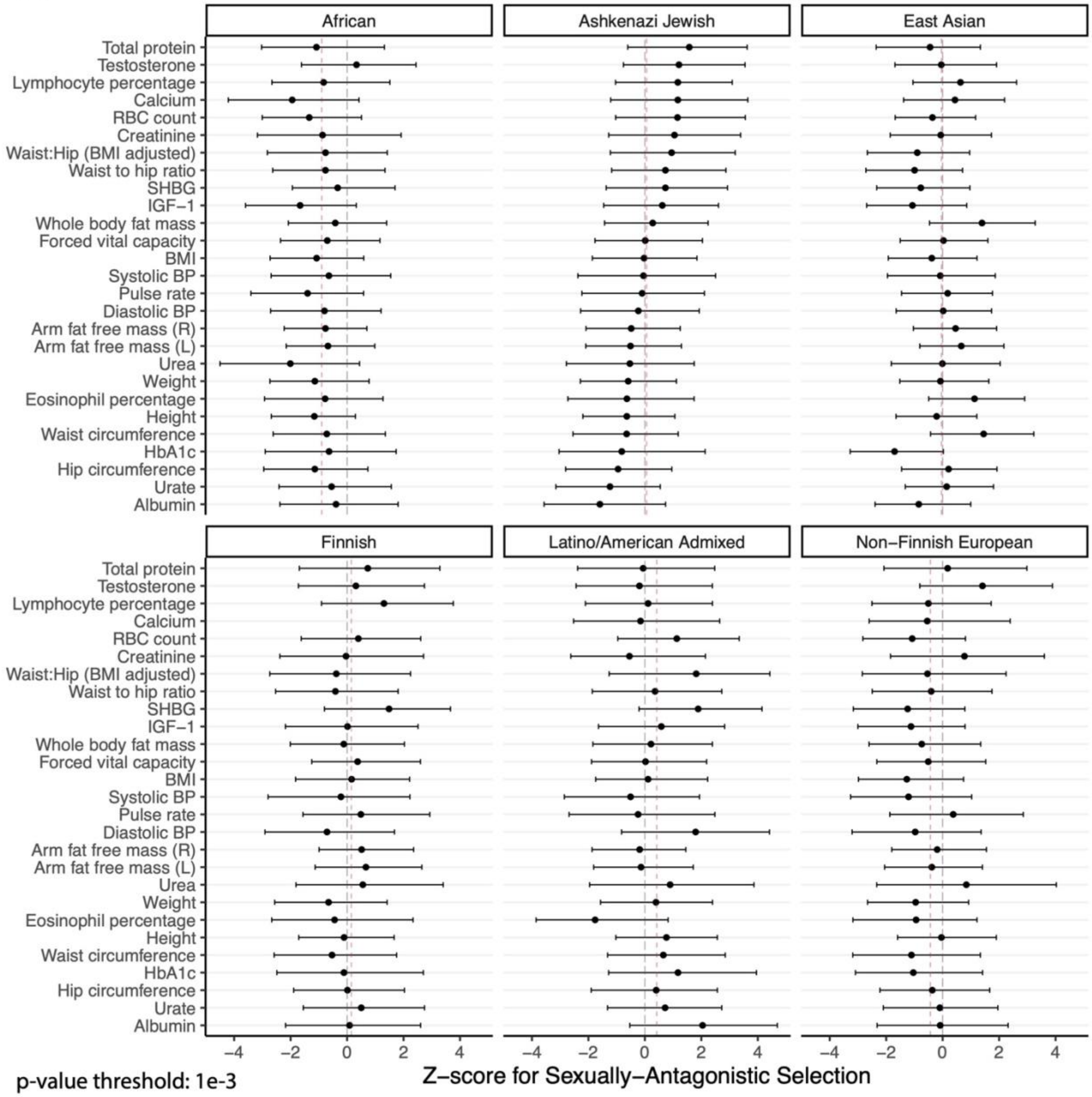

p-value threshold: 1e-3

- - - - mean Z score for all traits in the ancestry group

(B)

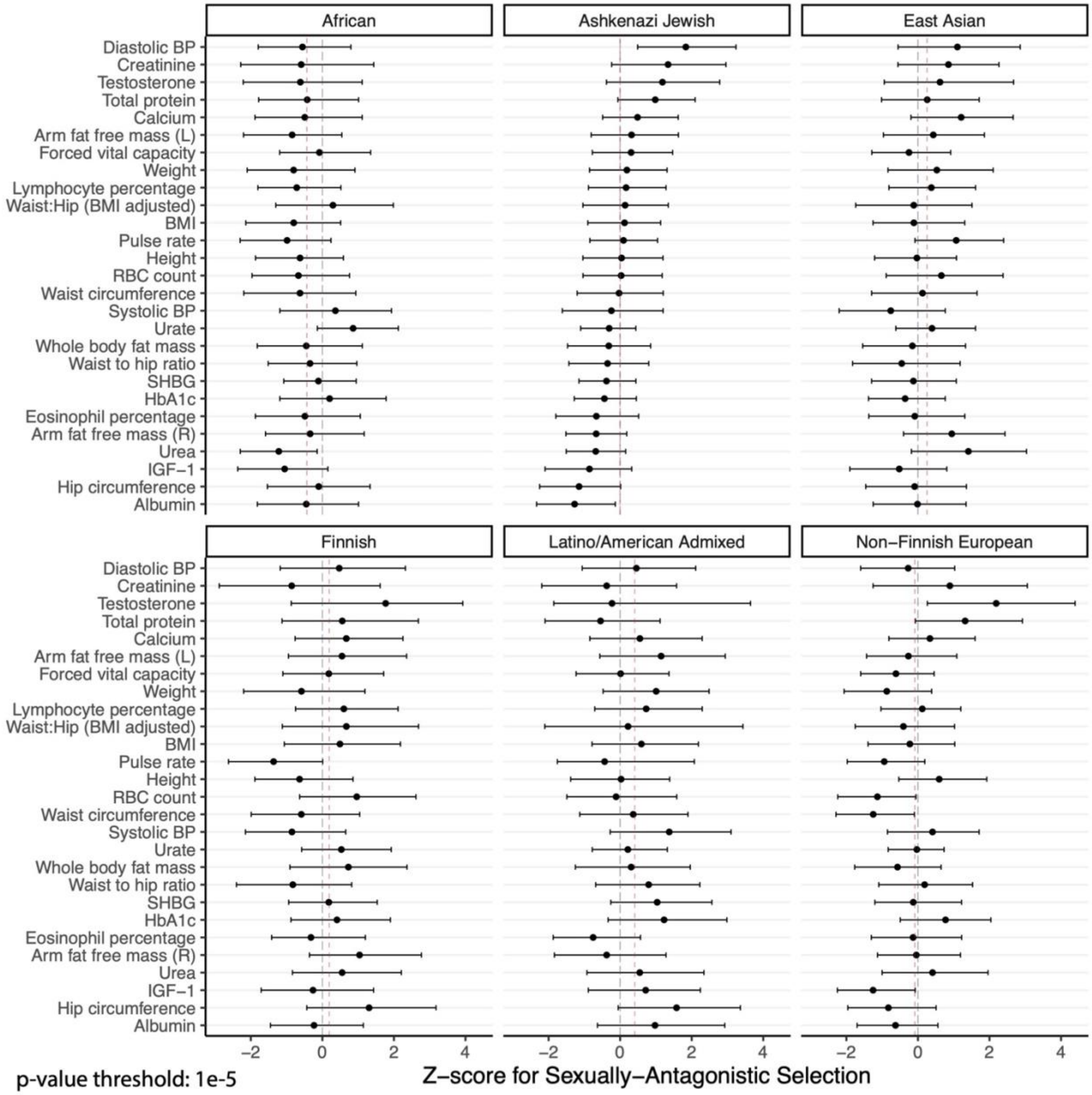

p-value threshold: 1e-5

---- mean Z score for all traits in the ancestry group

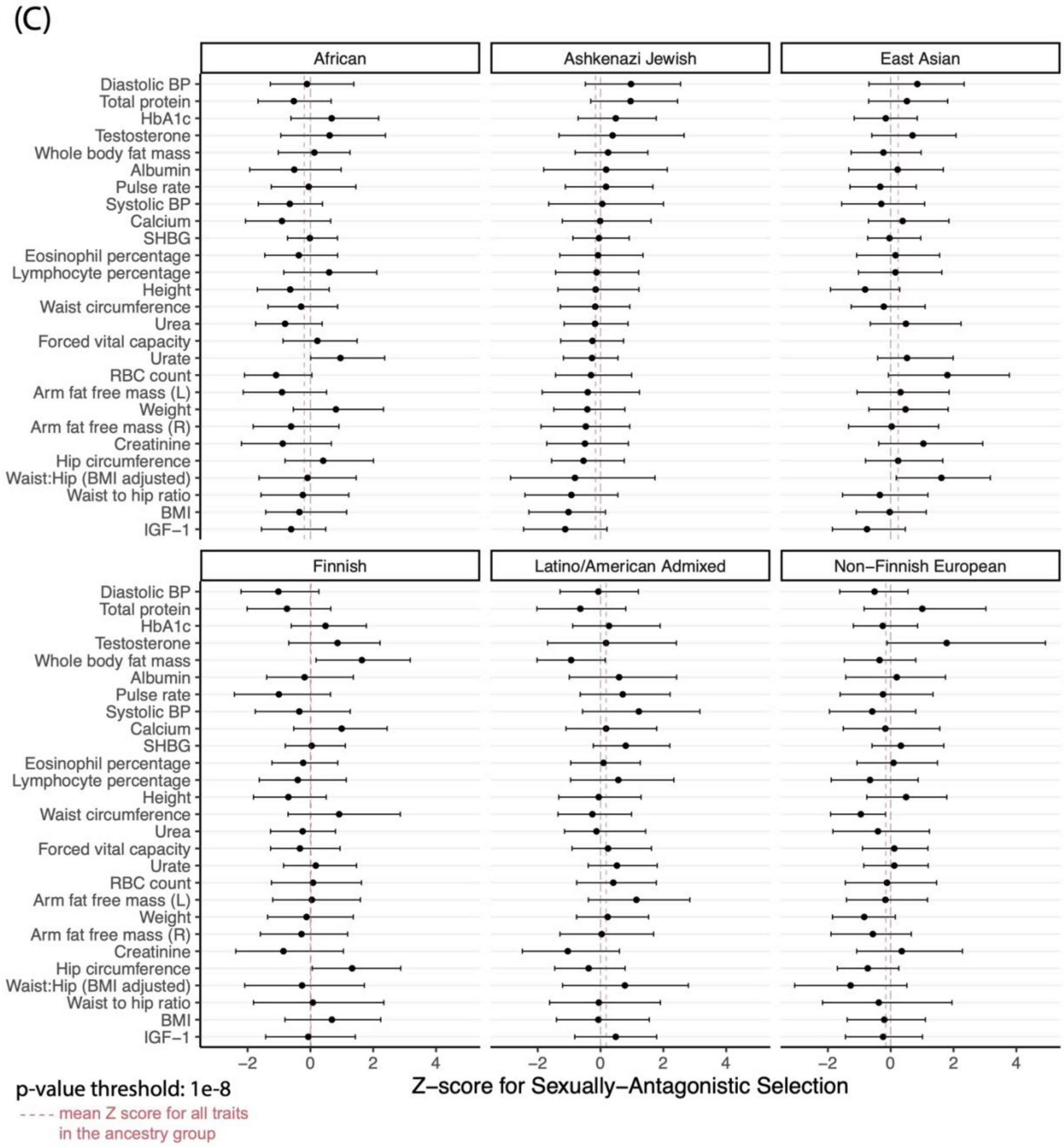

**Figure S21: Z-scores using UK Biobank data, related to Figure 7**

This figure parallels **Fig. 7** and **Fig. S20** but uses male-female allele frequency differentiation data from the UK BioBank. Here, the statistic $L_{ST}$ replaces $F_{ST}$, as explained in the section "**Estimating the potential for sexually-antagonistic selection on standing variation**" of the main text. The p-value cutoff for marginal sex-specific GWAS associations included in this analysis was set to $10^{-5}$.

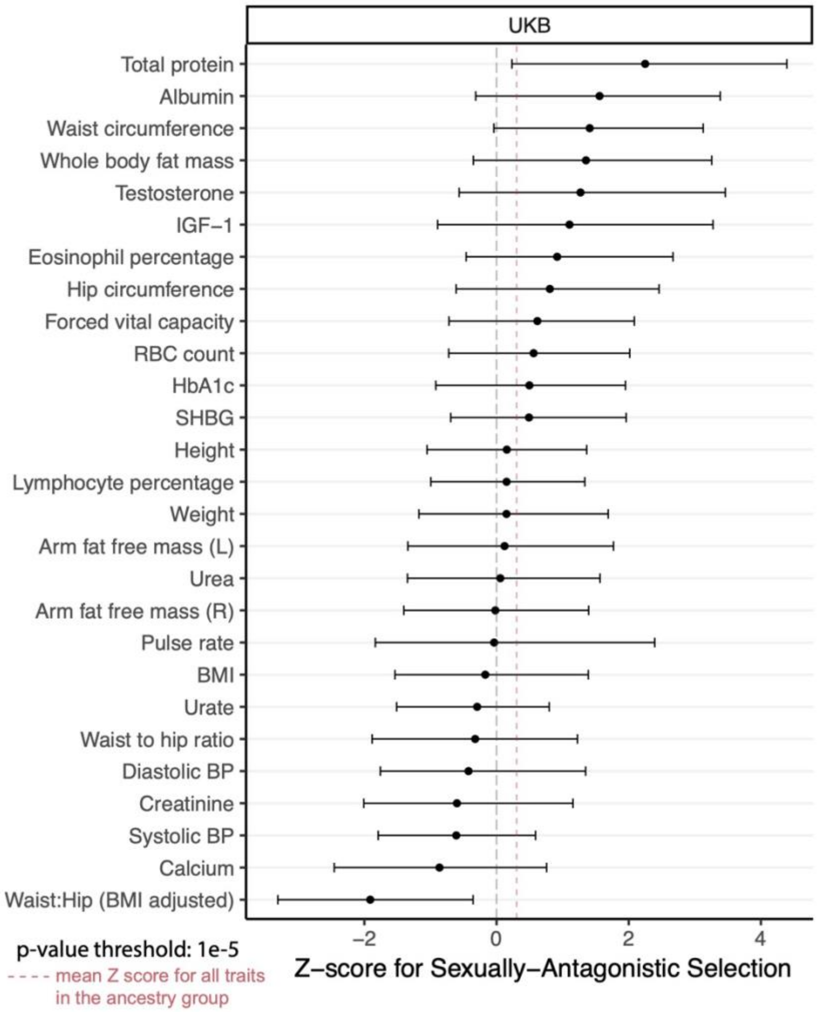

### Supplementary Tables

**Table S1: Filtering site with small sample sizes, related to Figure 7**

For the sexually-antagonistic selection analysis, we removed all sites with less than 1,000 alleles in the sample in order to narrow our research on sites with sufficiently large sample sizes. There were 2,285,169 sites before filtering.

| Ancestry | # Sites After Filtering |
|---|---|
| African/African American | 2,282,394 |
| Amish | 0 |
| Latino/American Admixed | 2,282,394 |
| Ashkenazi Jewish | 2,247,203 |
| East Asian | 2,255,778 |
| Finnish | 2,249,604 |
| Middle Eastern | 0 |
| Non-Finnish European | 2,283,952 |
| South Asian | 2,167,009 |
| Other | 2,080,336 |

**Table S2. Filtering monoallelic sites, related to Figure 7**

For the sexually-antagonistic selection analysis, and for each ancestry, we removed sites which were monoallelic in one or both sexes.

| Ancestry | # Sites after filtering |
|---|---|
| African/African American | 924,946 |
| Latino/American Admixed | 519,496 |
| Ashkenazi Jewish | 161,271 |
| East Asian | 300,323 |
| Finnish | 207,162 |
| Non-Finnish European | 1,104,008 |
| South Asian | 330,518 |
| Other | 267,812 |

**Table S3: Filtering sites that were not overlapping between the gnomAD dataset and the GWAS dataset, related to Figure 7**

For the sexually-antagonistic selection analysis, any sites which did not occur in both the filtered GWAS data from UKB and the and filtered gnomAD data were excluded.

| Ancestry | # Sites before filtering by GWAS-gnomAD overlap | # Sites after filtering | # Sites removed |
|---|---|---|---|
| African/African American | 924,946 | 886,467 | 38,479 |
| Latino/American Admixed | 519,496 | 506,760 | 12,736 |
| Ashkenazi Jewish | 161,271 | 160,358 | 913 |
| East Asian | 300,323 | 295,801 | 4,522 |

| | | | |
|---|---|---|---|
| Finnish | 207,162 | 204,914 | 2,248 |
| Non-Finnish European | 1,104,008 | 1,051,015 | 52,993 |
| South Asian | 330,518 | 325,444 | 5,074 |
| Other | 267,812 | 264,682 | 3,130 |

**Table S4: Filtering by p-value threshold, related to Figure 7**

For the sexually-antagonistic selection analysis, we used three different thresholds of GWAS p-values to further restrict which sites were used for analysis by focusing on sites with strong correlation with the trait of interest. Shown are the results of filtering for height, as an example. Results for other traits are highly similar.

| Ancestry | p-value threshold | # Sites before this filtering step | # Sites after filtering |
|---|---|---|---|
| Ashkenazi Jewish | 1e-3 | 160,358 | 2,749 |
| Ashkenazi Jewish | 1e-5 | 160,358 | 578 |
| Ashkenazi Jewish | 1e-8 | 160,358 | 298 |
| Finnish | 1e-3 | 204,914 | 3,975 |
| Finnish | 1e-5 | 204,914 | 852 |
| Finnish | 1e-8 | 204,914 | 373 |

**Table S5: Identification of SNPs susceptible to mis-mapping to sex chromosomes—Kasimatis et al., related to Figure 7**

We compare our results for excluding sites due to mis-mapping to the results from Kasimatis et al.[18], who removed from their analysis sites in regions with 90% identity or higher with a region in a sex chromosome (as we have, based on BLAT[25] matches), as well as sites overlapping probes of length 40bp or higher.

| 90% BLAT identity threshold | This study, susceptible to mismapping | This study, not identified as susceptible |
|---|---|---|
| Kasimatis et al., suceptible to mis-hybridization | 15,565 (2.51%) | 4,963 (0.80%) |
| Kasimatis et al., not identified as susceptible | 469,864 (75.78%) | 129,648 (20.91%) |

**Table S6: Characterizing GxSex based on independent analysis of individual sex-heterogenous SNPs, related to Figure 3**

We simulated GWAS data study arising from the covariance structure inferred for five traits. We then followed the method described in **Text S6** for calling SNPs as sex-heterogenous SNPs and categorizing them.

| Generative covariance structure | # sex-heterogenous SNPs | % sex-specific effects | % effects in the opposite direction | % effects in the same directions |
|---|---|---|---|---|
| Height | 994.7 (13.72) | 62.3% (0.6%) | 37.7% (0.6%) | 0.0% (0.0%) |
| BMI | 1527.7 (50.74) | 52.1% (2.7%) | 26.8% (0.9%) | 21.1% (0.9%) |
| Creatinine | 1243.6 (7.67) | 52.8% (0.4%) | 30.7% (0.4%) | 16.4% (0.4%) |
| IGF-1 | 1392.3 (10.82) | 52.2% (0.5%) | 29.1% (0.7%) | 18.7% (0.4%) |
| Systolic BP | 1349.2 (15.68) | 51.7% (0.3%) | 28.4% (0.3%) | 20.0% (0.4%) |
